# BacSC: A general workflow for bacterial single-cell RNA sequencing data analysis

**DOI:** 10.1101/2024.06.22.600071

**Authors:** Johannes Ostner, Tim Kirk, Roberto Olayo-Alarcon, Janne Gesine Thöming, Adam Z. Rosenthal, Susanne Häussler, Christian L. Müller

**Affiliations:** Computational Health Center, Helmholtz Munich, Ingolstädter Landstraße 1, 85764, Neuherberg, Germany; Institut für Statistik, Ludwig-Maximilians-Universität München, Geschwister-Scholl-Platz 1, 80539, Munich, Germany; Institute for Molecular Bacteriology, TWINCORE, Centre for Experimental and Clinical Infection Research, Hannover, Germany; Department of Clinical Microbiology, Rigshospitalet, Copenhagen, Denmark; Department of Microbiology and Immunology, University of North Carolina, Chapel Hill, NC, USA; Department of Molecular Bacteriology, Helmholtz Centre for Infection Research, Braunschweig, Germany; Center for Computational Mathematics, Flatiron Institute, 162 Fifth Avenue, New York, 10010, NY, USA

**Keywords:** bacterial single-cell RNA sequencing, phenotypic heterogeneity, statistical analysis, data processing, computational pipeline, data thinning, synthetic data generation, scanpy

## Abstract

Bacterial single-cell RNA sequencing has the potential to elucidate within-population heterogeneity of prokaryotes, as well as their interaction with host systems. Despite conceptual similarities, the statistical properties of bacterial single-cell datasets are highly dependent on the protocol, making proper processing essential to tap their full potential. We present BacSC, a fully data-driven computational pipeline that processes bacterial single-cell data without requiring manual intervention. BacSC performs data-adaptive quality control and variance stabilization, selects suitable parameters for dimension reduction, neighborhood embedding, and clustering, and provides false discovery rate control in differential gene expression testing. We validated BacSC on a broad selection of bacterial single-cell datasets spanning multiple protocols and species. Here, BacSC detected subpopulations in *Klebsiella pneumoniae*, found matching structures of *Pseudomonas aeruginosa* under regular and low-iron conditions, and better represented subpopulation dynamics of *Bacillus subtilis*. BacSC thus simplifies statistical processing of bacterial single-cell data and reduces the danger of incorrect processing.

## 1 Introduction

Single-cell RNA sequencing (scRNA-seq) has revolutionized genetic analysis of eukaryotic cell compendia by allowing researchers to extract individual cells’ gene expression profiles and obtain new insights on intracellular mechanisms, as well as the structure and dynamics within entire populations of cells [1–3]. These advances have led, among others, to a better understanding of immune responses [4], disease progression [5], or advancements in drug development [6]. Consequently, similar insights into microbial heterogeneity are expected from scRNA-seq of bacterial populations, opening up new avenues for assessing antimicrobial resistance, evolutionary pathways, or within-population differences in response to external conditions [7]. In addition, bacterial scRNA-seq yields new ways to analyze interactions between the isogenic microbiome and host systems, for example in toxin regulation [8, 9], formation of metabolic niches [10], and the analysis of microbial spatial heterogeneity [11].

Applying scRNA-seq technologies to bacteria has however proven to be challenging, e.g. due to low overall transcript abundance, the short half-life of bacterial mRNA, and difficulties in cell lysis due to sturdier cell walls [12–15]. Recently, multiple protocols have been developed that enable scRNA-seq of bacteria on larger scales by tackling these challenges in different ways [13, 14, 16–19]. For example, ProBac-seq [18] uses a library of oligonucleotide probes to target mRNAs, while BacDrop [13] uses a two-stage cell barcoding procedure to increase cell numbers.

Datasets from scRNA-seq contain gene expression counts for each UMI (unique molecular identifier) and are typically sparse, high-dimensional and noisy, requiring specialized methods and particular care in their statistical processing to obtain biologically meaningful representations [20, 21]. This process has been extensively discussed for eukaryotic cells, leading to well-documented benchmarks [22, 23], best practices [24–26], and methods to select adequate hyperparameters [27–29] for each step of the statistical analysis pipeline. For bacterial scRNA-seq, no such guidelines exist yet, prompting the use of default parameters and methods without prior assessment of their statistical validity and suitability for the data at hand. This may, however, lead to suboptimal or even flawed representations of the data, which can severely impact the quality of biological insights gained from downstream analyses.

Each step in a typical statistical processing pipeline developed for the analysis of eukaryotic scRNA-seq [24, 26] poses new statistical challenges when applied to bacterial scRNA-seq data:

- In quality control, differences in sparsity and sequencing depth have to be accounted for when filtering out low-quality genes or cells [30].
- Variance stabilization is a crucial step to ensure comparability for all sequenced cells, but scaling the data to a common sequencing depth and the choice of an imputation value for zero replacement must be done with the statistical properties of the data in mind [21, 22].
- The number of principal components used for low-dimensional data representation, as well as the number of neighbors and minimal distance used in UMAP embeddings, are hyperparameters that are commonly chosen in a heuristic fashion, but have a significant impact on downstream analysis and visual representation of the data [27, 31].
- The resolution parameter in cell type clustering is also often determined by visual trial-and-error procedures [32].
- Finally, recent studies show that differential expression testing between cell types suffers from a double-dipping issue that inflates the false discovery rate [29] if not accounted for.

In this study, we address these challenges by developing a standard workflow for processing bacterial scRNA-seq gene expression data that does not require the selection of modeling choices or manual tuning of parameters. We introduce BacSC, a computational pipeline for automatic processing of scRNA-seq data that is applicable to datasets generated by various bacterial scRNA-seq protocols. BacSC reevaluates the validity of methods used in each of the steps outlined above in the context of bacterial scRNA-seq, adjusts methods if necessary, and automatically chooses suitable hyperparameters in a data-driven way. To this end, BacSC provides tools for data integration and quality control of bacterial scRNA-seq data, and performs a simple, yet powerful variance stabilizing transform that is suitable for scRNA-seq data with varying sequencing depth and high zero inflation. Using techniques from data thinning [31, 33] and knockoff generation [27, 34], BacSC is able to select suitable parameters and perform dimensionality reduction, neighborhood embedding, and cell-type clustering without requiring user intervention. BacSC also offers FDR control for differential expression testing of bacterial scRNA-seq data through contrasting p-values with synthetic null data[29, 35].

To validate the steps taken in BacSC, we compared the statistical properties of 13 datasets generated with ProBac-seq [18, 36] and BacDrop [13], emphasizing their low sequencing depth, high zero inflation, and differences in marginal gene distribution. As a proof of concept, BacSC was able to distinguish the same cell types as previously shown through analysis with default or manually chosen parameters for all datasets with known biological structure. BacSC additionally showed improved ability to describe the transitional nature of cell competence in *B. subtilis*, was able to give a more clear distinction of cells expressing mobile genetic elements in *K. pneumoniae*, and discovered new cellular subpopulations in *K. pneumoniae* and *P. aeruginosa*. When applied to a combination of *P. aeruginosa* cells grown under regular and iron-reduced conditions, BacSC was able to simultaneously integrate cells from both conditions based on their gene expression profiles and detect differential expression of genes related to iron acquisition.

BacSC is available as a modular framework in Python that seamlessly integrates into the scanpy [37] workflow and allows for direct downstream analysis with other tools from the scverse [38]. BacSC is available on GitHub (https://github.com/bio-datascience/BacSC).

## 2 Results

### 2.1 Explorative comparison of bacterial scRNA-seq technologies reveals differences in key statistical properties

To ensure the cross-platform and cross-species applicability of BacSC, we gathered a total of 13 bacterial scRNA-seq datasets that were generated with two different sequencing protocols, ProBac-seq [18, 36], and BacDrop [13] (see section 3). The datasets encompass five bacterial species (*Pseudomonas aeruginosa, Bacillus subtilis, Klebsiella pneumoniae, Escherichia coli, Enterococcus faecium*), further distinguished by strain, growth environment, or treatment condition (Table 1).

**Table 1.** Description of datasets used to benchmark BacSC. All datasets are named by the convention *species condition protocol*. Datasets from ProBac-seq are marked with the suffix ” PB”, datasets from BacDrop are marked with ” _BD”

| Dataset | Species/strain | Condition | Protocol | Source |
| --- | --- | --- | --- | --- |
| Pseudomonas_balanced_PB | *P. aeruginosa* PAO1 | balanced growth | ProBac-seq | This study |
| Pseudomonas_li_PB | *P. aeruginosa* PAO1 | Low Iron environment | ProBac-seq | This study |
| Ecoli_balanced_PB | *E. coli* MG1655 | balanced growth | ProBac-seq | This study |
| Bsub_minmed_PB | *B. subtilis* 168 | minimal media | ProBac-seq | McNulty et al. [18] |
| Bsub_damage_PB | *B. subtilis* 168 | DNA damage induced by Mitomycin C | ProBac-seq | This study |
| Bsub_MPA_PB | *B. subtilis* 168 | MPA energy stress | ProBac-seq | This study |
| Klebs_antibiotics_BD | *K. pneumoniae* MGH66 | 6 samples, treated with one of 3 antibiotics (2 samples each): Meropenem, Gentamicin, Ciprofloxacin | BacDrop | Ma et al. [13] |
| Klebs_untreated_BD | *K. pneumoniae* MGH66 | Untreated culture (2 samples) | BacDrop | Ma et al. [13] |
| Klebs_BIDMC35_BD | *K. pneumoniae* BIDMC35 | Untreated culture | BacDrop | Ma et al. [13] |
| Klebs_4species_BD | *K. pneumoniae* MGH66 | Untreated culture | BacDrop | Ma et al. [13] |
| Ecoli_4species_BD | *E. coli* 10B | Untreated culture | BacDrop | Ma et al. [13] |
| Efaecium_4species_BD | *E. faecium* EnGen0052 | Untreated culture | BacDrop | Ma et al. [13] |
| Pseudomonas_4species_BD | *P. aeruginosa* PAO1 | Untreated culture | BacDrop | Ma et al. [13] |

The number of genes per dataset was mostly dependent on the species (Figure 2A), and ranged between 5,572 (*P. aeruginosa*) and 2,350 (*E. faecium*). The sequencing depth per cell was highly dependent on the sequencing method, with data from BacDrop showing a median sequencing depth between 2 and 43, while all datasets generated with ProBac-seq had at least a median sequencing depth of 150 (Figure 2B). In contrast, datasets generated with BacDrop generally encompassed a higher number of cells (median 9,936) than datasets from ProBac-seq (median 3,773).

**Fig. 1.**
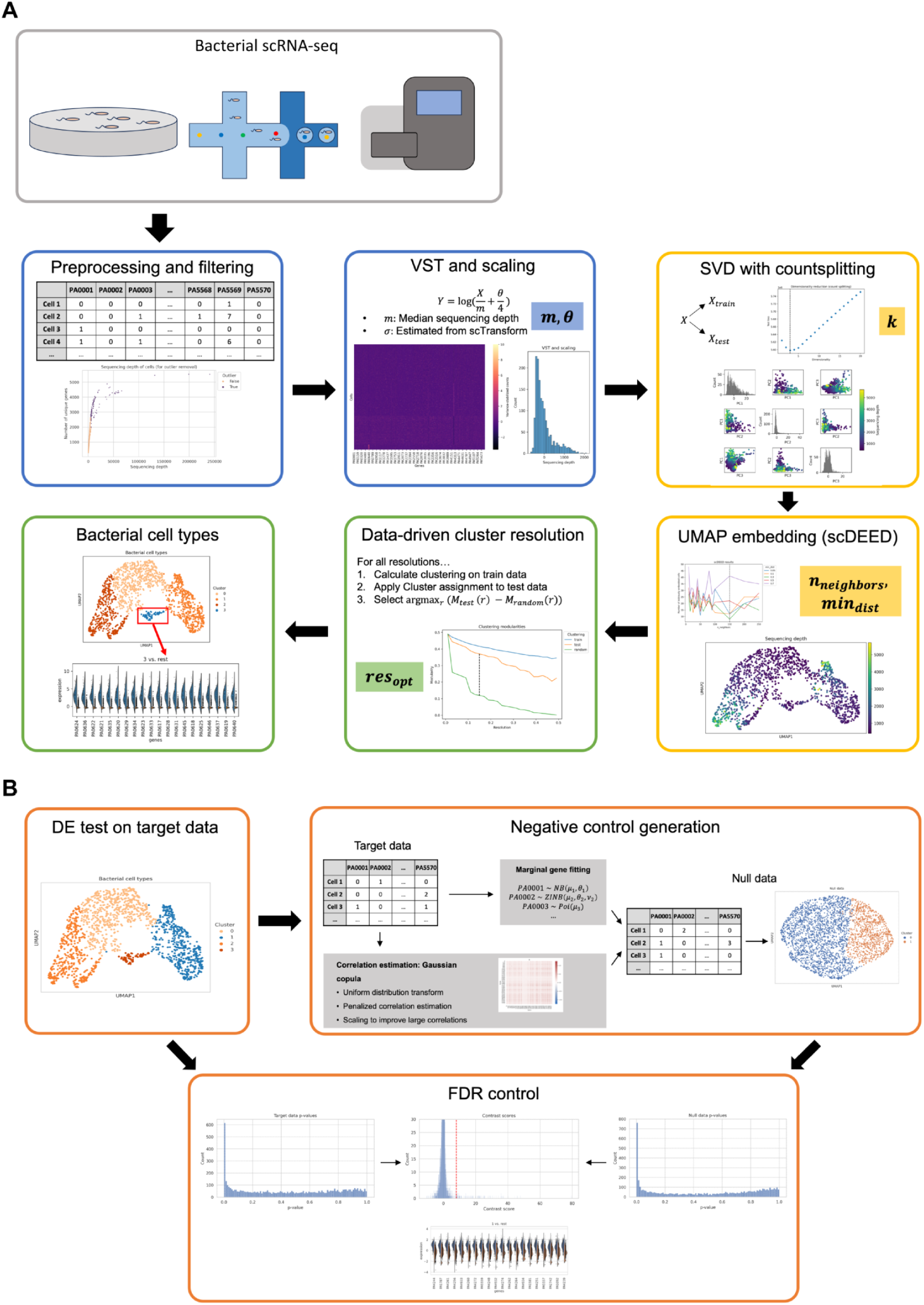
Conceptual visualization of the BacSC pipeline. **(A)** Bacterial single-cell RNA sequencing produces a table of read counts. Sterting with this table, BacSC first filters out outlier cells, before performing a variance stabilizing transform (blue boxes). Next, the latent data dimensionality is determined through count splitting, and suitable parameters for UMAP visualizations are determined by scDEED (yellow boxes)-Finally, BacSC determines an adequate resolution for clustering, and is able to discover bacterial cell types (green boxes). The colored rectangles show the most important key parameters calculated **in** the respective step of BacSC. **(B)** For differential expression testing, BacSC generates synthetic null data with the same marginal distributions as the target data through a Gaussian copula approach. P-values of a DE test on the target data are then contrasted with p-values on the synthetic null data to obtain differentially expressed genes at a desired FDR level.

**Fig. 2.**
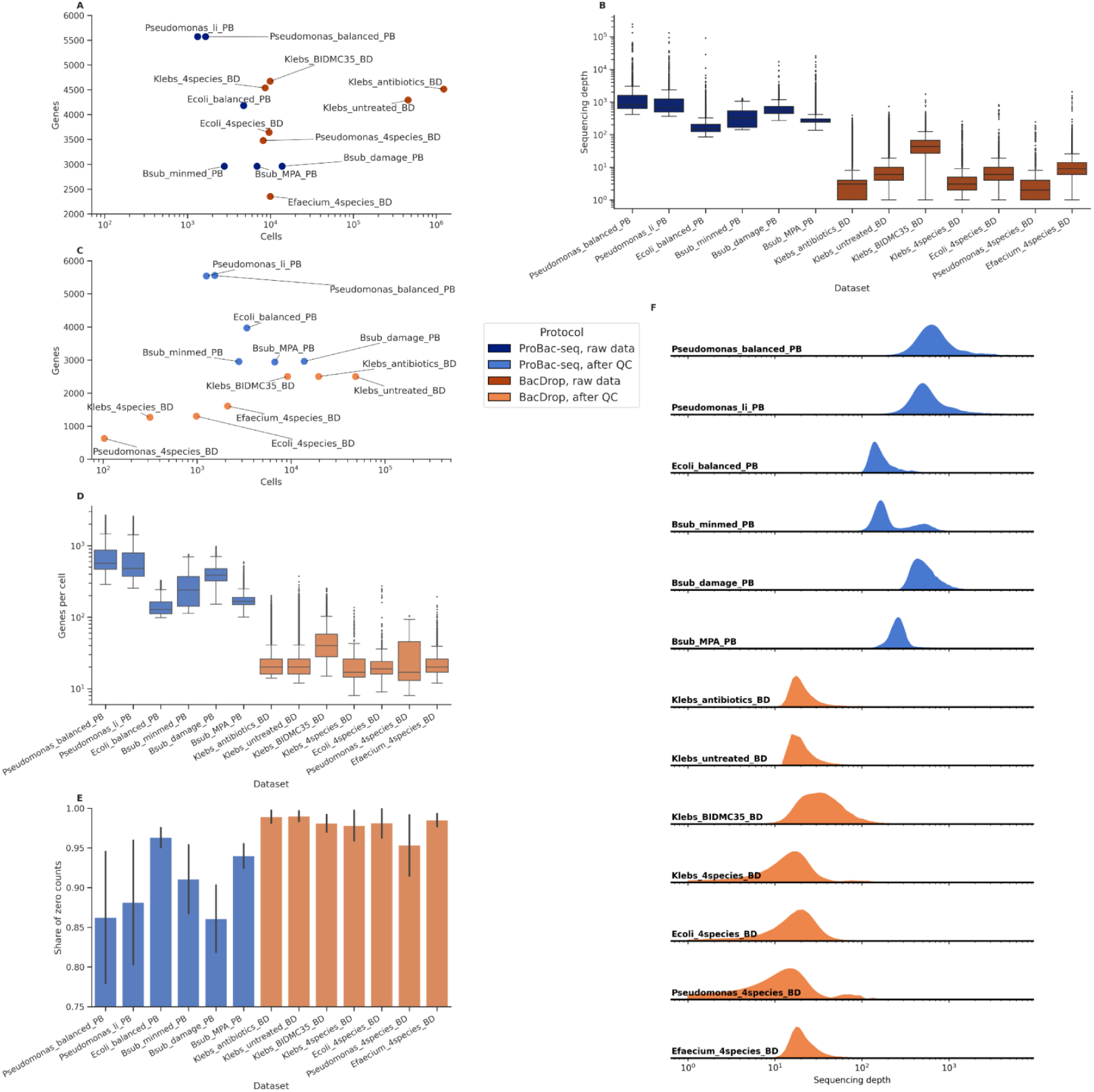
Explorative comparison of bacterial scRNA-seq technologies reveals differences in key statistical properties. **(A)** Number of genes and cells before quality control. **(B)** Sequencing depth per cell before quality control. The box depicts the 25% and 75% quartiles of the data, as well as the median; whiskers extend to 1.5 times the interquartile range of the data. **(C)** umber of genes and cells after quality control. **(D)** Number of expressed genes per cell after quality control. The box depicts the 25% and 75% quartiles of the data, as well as the median; whiskers extend to 1.5 times the interquartile range of the data. **(E)**Share of zero counts over all cells in the raw data matrices after quality control. Errorbars show the empirical standard deviation. **(F)** Density plots of sequencing depth for each dataset after quality control.

After filtering out cells with abnormally low or high expression and genes without reads in more than one cell (See section 2.2), both protocols could be easily distinguished by the number of genes detected, with all datasets from ProBac-seq encompassing at least 2,922 genes, while datasets from BacDrop contained a maximum of 2,500 genes (Figure 2C, Table E1). This was in part due to the subsetting to 2,500 highly variable genes, which was only performed on the *Klebs antibiotics BD, Klebs untreated BD*, and *Klebs BIDMC35 BD* datasets. The BacDrop data from the four species comparison comprised a much lower numbers of genes (628 - 1606) without selection of highly variable genes. The number of cells generally differed more within the BacDrop data (103 - 48,511), while the ProBac-seq datasets had much more stable cell numbers (1,910 - 13,801; Figure 2C, Table E1).

BacDrop only detected between 24 and 47 unique genes per cell on average, while ProBac-seq covered at least 49 genes for each cell in every dataset (Figure 2D). Consequently, ProBac-seq had less zero entries in the filtered read count matrices, with zeroes making up between 86% and 97% of all entries, while BacDrop showed zero inflation numbers between 95% and 99.2% (Figure 2E). After quality control, we observed similar discrepancies between protocols in sequencing depth. ProBac-seq not only covered more genes per cell, but was also able to capture more transcripts, with median sequencing depths ranging from 103 to 794.5. BacDrop datasets only had a median sequencing depth of 45 or less after quality control (Figure 2F; Table E1). We therefore reasoned that the usage of multiple probes per gene and subsequent aggregation through max-pooling in ProBac-seq (see Methods, [36]) leads to higher genome coverage and sequencing depth for each cell.

### 2.2 Description of the BacSC pipeline

At its core, statistical processing of scRNA-seq data extracts information from raw transcriptome reads by filtering, normalization, dimension reduction, and clustering steps [24, 26]. BacSC selects suitable methods and automates the choice of hyperparameters for each step without the need for manual intervention (except for quality control; Figure 1). section 2.2 briefly describes each step, while we give more detailed descriptions in the ”QUANTIFICATION AND STATISTICAL ANALYSIS” part of the STAR methods. First, the data is subjected to quality control to filter out barcodes with abnormally low or high gene expression (Figure 1A). Because our exploratory analysis showed that bacterial single-cell data differs heavily in terms of average sequencing depth, number of expressed genes, and zero inflation, this step is highly dependent on the experimental protocol used. Therefore, BacSC leaves this step as the only point where manual intervention is necessary, but provides tools for outlier detection through median absolute deviation (MAD) statistics [30] and aggregating probe-based data from ProBac-seq. As with eukaryotic scRNA-seq data, the main data object after quality control in each dataset is a *n* × *p*-dimensional count table *X*, containing the read counts of *p* features for *n* cells.

Next, the read count data must be normalized and scaled. Because bacterial scRNA-seq data shows greatly reduced sequencing depth and increased zero inflation compared to eukaryotic scRNA-seq, special care has to be taken in this step [39, 40]. BacSC first scales each cell individually to have the same number of reads, and subsequently log-transforms the data. The pseudocount introduced in this step is gene-specific [22], with overdispersion parameters calculated through sctransform [41] (Figure 1A). Finally, each gene is scaled to have zero mean and unit variance over all cells.

After variance stabilization, the data is reduced to a lower-dimensional representation by singular value decomposition (SVD) on the data. The embedding dimensionality *k* in this step of the scRNA-seq processing workflow is often set manually, e.g. by finding an ”elbow” in the plot of SVD loadings [25]. BacSC instead uses a count-splitting approach to find a good value for *k*, which was described by Neufeld et al. [31]. For this, the raw counts after quality control are split into a training and test dataset, and the variance-stabilizing transform is applied to both datasets. Then, the latent dimensionality *k* with minimal reconstruction error between the *k*-dimensional embedding of the training data and the full test data is chosen (Figures 1A, B1).

UMAP (Uniform Manifold Approximation and Projection) plots [42] are a popular tool for two-dimensional visualization of scRNA-seq data to preserve the local structure and point out global differences in higher-dimensional data. The algorithm is largely dependent on three parameters - the latent dimensionality *k*, the number of neighbors *n_neighbors_* considered for each cell, as well as the minimal distance *min_dist_* between points. These parameters are often adjusted manually until a satisfactory picture arises. To eliminate this manual step, BacSC uses the negative-control approach described by scDEED [27] to determine the latter two latent parameters. scDEED calculates a reliability score - the correlation between the distance vectors from a cell to its neighbors before and after UMAP embedding - and compares them to the distribution of contrast scores on a randomly permuted dataset (Figures 1A, B1). It then selects the parameter combination for which the amount of cells with abnormally low reliability scores is minimized.

Cell clusters in scRNA-seq data are typically detected through the the Louvain [43] and Leiden [44] algorithms. Both algorithms aim to maximize the modularity of partition over all cells with respect to a resolution parameter *res*. Once again, this parameter is usually chosen manually to fit the structure observed in the UMAP or PCA embeddings. Computational determination of a feasible resolution parameter that robustly detects cell clusters without creating too many subclusters is, however, not straightforward. BacSC uses the train and test dataset obtained from count splitting and introduces a new gap statistic based on the difference in modularity between two clusterings on the test data - one calculated on the train data and one assigned randomly. Maximizing this gap statistic allows to find a value for *res* for which the obtained clustering on the train data also generalizes well to the structure of the test data Figures 1A, B2).

Bacterial single-cell sequencing allows to characterize heterogeneity within bacterial populations in unprecedented detail. The discovery of subpopulations and the description and interpretation of different cell types in bacterial populations is therefore still at an early stage. To characterize previously unknown cell types, automatic selection of signature genes for each cluster is often achieved through differential expression (DE) testing [24]. For this task, BacSC provides capabilities for DE testing that takes the recently popularized problem of ”double dipping” for DE testing of cell types into account [29, 45, 46]. In short, using the same information (gene expression) to define a clustering as well as the subsequently determining DE genes to characterize these clusters results in an inflated false discovery rate (FDR). BacSC solves this issue by adapting the ClusterDE method [29] for FDR control. Due to the highly sparse nature of bacterial single-cell data, BacSC uses a modified version of scDesign2 [47] to generate the synthetic null data. Further, BacSC also adapts ClusterDE to achieve better results for highly uneven cluster proportions (Figures 1B, B3).

To validate our pipeline, we applied BacSC to all datasets described in Table 1. For quality control, we manually set dataset-specific filtering parameters on minimal sequencing depth and MAD cutoff (Table E2), based on visual inspection of the distribution of sequencing depth and number of unique genes per cell. After variance stabilization, we further reduced the *Klebs antibiotics BD, Klebs untreated BD*, and *Klebs BIDMC35 BD* datasets to 2,500 highly variable genes based on their standardized variances [48]. All other steps of BacSC do not require any manual intervention, and were thus performed automatically. The determined data distribution, as well as parameters for latent dimensionality, number of neighbors, minimal distance, and clustering resolution are shown in Table E2.

### 2.3 BacSC uncovers new biological structures in datasets obtained from different bacterial scRNA-seq protocols

#### 2.3.1 Transitions between cellular states in *B.subtilis* are pronounced by BacSC

To show the validity of the transformations and parameters selected in BacSC, we first investigated the *Bsub minmed PB* dataset (Figures 3, D18). This data was generated by [18] to validate the ProBac-seq method. The original analysis with default parameters in Seurat [48] discovered four distinct subpopulations with multiple subclusters and different functionality. In the first two dimensions of the PCA embedding suggested by BacSC, three larger subpopulations were immediately apparent (Figure 3A), while a fourth cluster with only 20 cells emerged in the UMAP embedding with BacSC’s selected parameters (Figure 3B). Clustering with the automatically determined resolution resulted in five cell type clusters (Figure 3B).

**Fig. 3.**
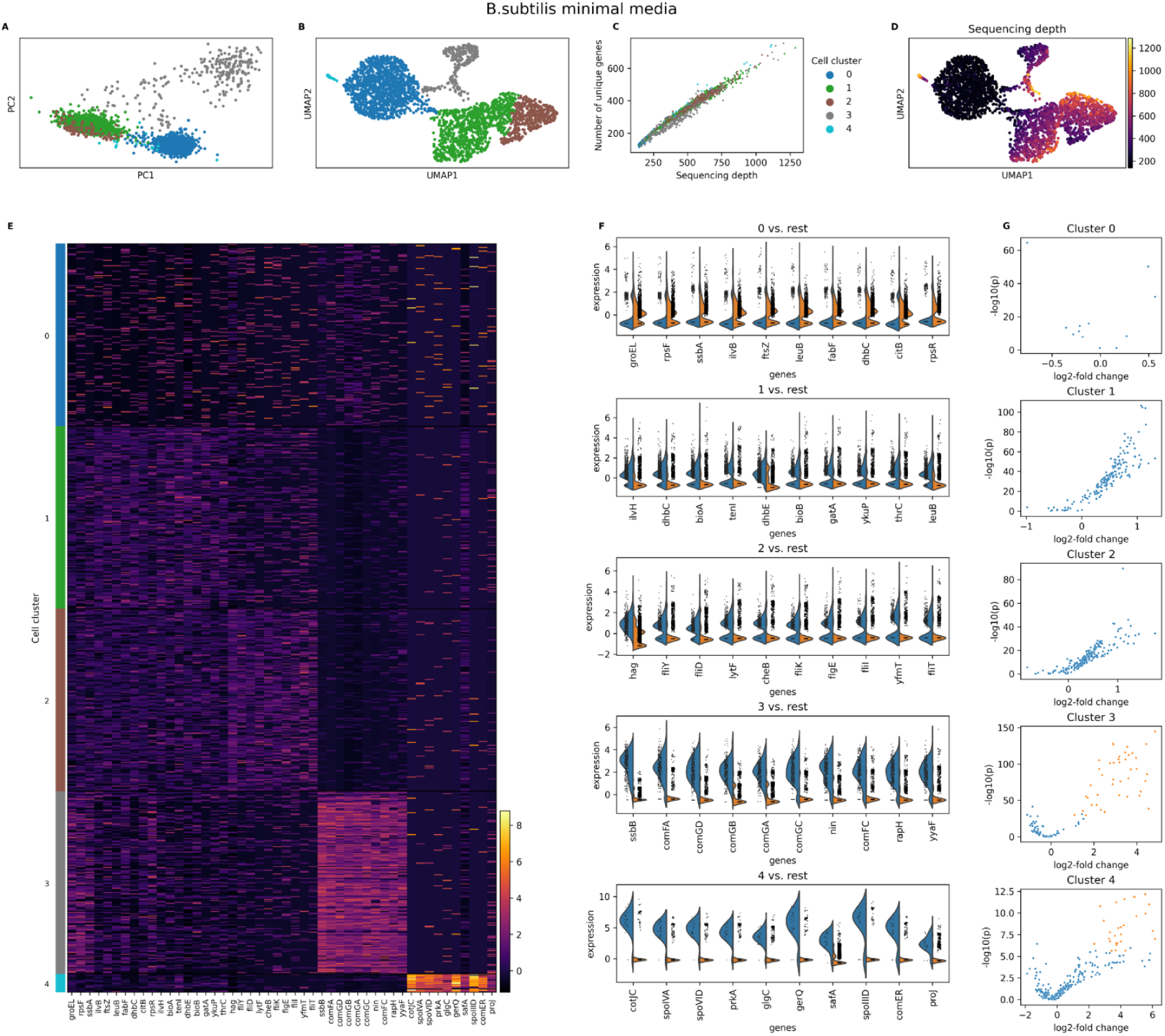
’Iransitions between cellular states in *B.subtilis* are pronounced by BacSC. Analysis of the *Bsub_minmed_PB* dataset with BacSC. **(A)** Scatterplot of first two dimensions of PCA embedding with cell type clusters highlighted. **(B)** UMAP plot based on the parameters determined by BacSC, colored by cell cluster. **(C)** Scatterplot of sequencing depth versus number of unique genes per cell, colored by cell cluster. **(D)** Umap plot as in (B), colored by sequencing depth. **(E)** Heatmap of normalized gene expression. For each cluster, the 10 genes with the highest contrast scores are shown. For better visibility of small clusters, at most 200 cells per cluster are shown. **(F)** Violin plots of normalized gene expression for DE tests of each cell type (blue) against the rest of the cell population (orange). For each cluster, the 10 genes with the highest contrast score are shown. (G) Volcano plots for DE tests as in (F). The x-axis shows the log-fold change for gene expression, the y-axis shows the - log_10_-transformed (uncorrected) p-value. Only genes that are expressed in at least 80% of cells in the respective cluster are shown. Orange genes are differentially expressed at the a = 0.05-level.

Because of the ”double-dipping” issue described above, DE testing produced large numbers for genes with very small p-values for each cell type (Figure D18I). Counteracting this through the p-value correction in BacSC revealed characteristic genes for each cell type (Figure 3E-G), but only the two smallest clusters (3 and 4) had genes significant at a FDR level of *α* = 0.05 (Figure D18J, Table E4).

Cell type 4 showed increased expression of many sporulation genes (*spoIVA, spoVID, spoIID*), while the marker genes in cell type 3 contained many genes associated with cell competence (*comFA, comGD, comGB, comGA, comGC, comFC*). These subpopulations were also found as clusters 9, respectively 6/8 in [18]. Cell type 0 contained cells with very low sequencing depths (Figure 3C, D), and many genes were significantly underexpressed at an FDR level of 0.1 (Table E4). The genes with the highest contrast scores for this cell type partially overlapped with genes found in clusters 0 and 3 in the original publication. Similarly, cell type 1 contained many upregulated genes at an FDR of 0.2. For cell type 2, many structural flagella components (*fliY, fliD, fliK, fliI, fliT*) were among the genes with the highest contrast scores, but only differentially expressed at an FDR level of 0.26. The region containing cell types 1 and 2 from BacSC therefore corresponds to clusters 1, 2, 3, and 5 from [18].

Notably, the UMAP from BacSC showed continuous streams of cells between the cell types, especially between cell types 0, 1, and 3 (Figure 3B), which were not visible in the original analysis [18]. We suspected these cells to be in a transitional phase between two cell states. The development of competent cells (cell type 3) is known to be procedural [49], which explains the transition of cells in and out of this cell type.

#### 2.3.2 BacSC shows clear differences in response of *K. pneumoniae* to different antibiotics

To showcase the applicability of BacSC to data from different bacterial scRNA-seq protocols, we revisited an analysis of six samples of *Klebsiella pneumoniae* generated with BacDrop [13]. The *Klebs antibiotics BD* dataset contains two replicates for each of three antibiotic treatments, ciprofloxacin, meropenem, and gentamicin.

Despite the high sparsity of the data (99.2%, Table E1), BacSC was able to successfully integrate all six samples. The first two principal components already showed heterogeneity in the data in the form of three clear subpopulations (Figure 4A). This was enhanced through the UMAP plot and data clustering (Figure 4B), which revealed two major clusters of cells that split up into two, respectively three cell types, and three small cell clusters. For all cell types, a subset of genes was differentially expressed at FDR levels of 0.2 or lower (Table E5).

**Fig. 4.**
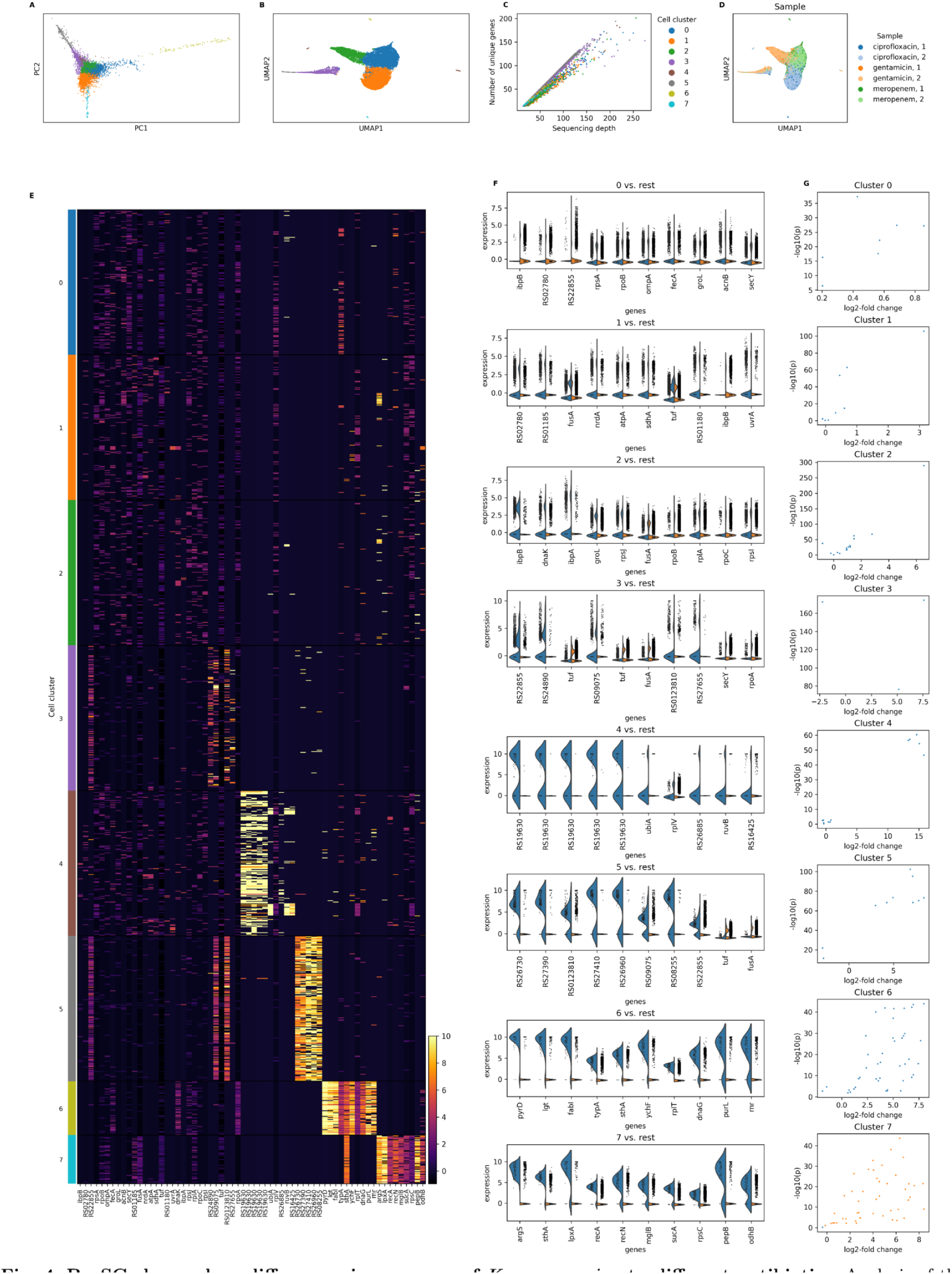
BacSC shows clear differences in response of *K. pneumoniae* to different antibiotics. Analysis of the *Klebs_antibiotics_BD* dataset with BacSC. **(A)** Scatterplot of first two dimensions of PCA embedding with cell type clusters highlighted. **(B)** UMAP plot based on the parameters determined by BacSC, colored by cell cluster. **(C)** Scatterplot of sequencing depth versus number of unique genes per cell, colored by cell cluster. **(D)** Umap plot as in (B), colored by sample identity (antibiotic and replicate). **(E)** Heatmap of normalized gene expression. For each cluster, the 10 genes with the highest contrast scores are shown. For each cluster, the 10 genes with the highest contrast scores are shown. For better visibility of small clusters, at most 200 cells per cluster are shown. **(F)** Violin plots of normalized gene expression for DE tests of each cell type (blue) against the rest of the cell population (orange). For each cluster, the 10 genes with the highest contrast score are shown. Genes in (E) and (F) are annotated with gene symbols wherever possible, otherwise locus tags are shown. (G) Volcano plots for DE tests as in (F). The x-axis shows the log-fold change for gene expression, the y-axis shows the - log_10_-transformed (uncorrected) p-value. Only genes that are expressed in at least 50% of cells in the respective cluster are shown. Orange genes are differentially expressed at the *a* = 0.05-level.

The cell types contained in the largest cellular subpopulation (0, 1, and 2) almost perfectly matched the separation by antibiotics shown in Figure 4D. Within these clusters, cells from both samples were distributed evenly, suggesting no residual batch effects. Cell types 3 and 5 made up all cells in the second large subpopulation, which contained a higher number of unique expressed genes than the rest of the dataset (Figure 4C). Both of these clusters showed significant differential expression of IS903B transposase-related genes (*RS09075, RS22855*), which matches the subpopulation of mobile genetic elements (MGE) described by [13]. Contrary to the original analysis, this subpopulation separated more from the bulk of the cells in BacSC’s UMAP embedding (Figure 4B). The small subpopulations (Cell types 4, 6, 7) were all characterized by a few genes that were barely expressed in other cells.

### 2.4 Processing with BacSC discovers a distinct response of *P. aeruginosa* to a low-iron environment

#### 2.4.1 Bacterial cell types of exponentially grown *P. aeruginosa* are similar in growth conditions with differing iron availability

We next tested if BacSC could recover environment-specific microbial cell types from bacterial cultures grown under different external conditions. For this, we investigated the *Pseudomonas balanced PB* and *Pseudomonas li PB* datasets. Both datasets contain cells from *P. aeruginosa* in exponential growth in minimal media, and sequenced with ProBac-seq. For the first sample, cells were grown in regular minimal media (MOPS with 10 µM FeSO_4_), while for the second sample, bacteria were exposed to a mild iron limitation (0.5 µM FeSO_4_), which resembles a growth condition mimicking competition between host and pathogen for the essential trace element during infection.

We first processed each dataset individually with BacSC. The diagnostic plots for both datasets (D22, D23) showed that normalized sequencing depths, as well as latent dimensionality, neighborhood embedding, and clustering resolution parameters found by BacSC were very similar. The PCA and UMAP embeddings for both datasets also showed similar patterns (Figures 5A, B, C6A, B, C7A, B). The sequencing depth vs. genome coverage plots (Figures 5D, E) revealed that in both populations, a subset of cells had lower coverage at high sequencing depths. This subgroup was identified as cluster 1 in the cell type clustering. Both datasets further contained two larger subpopulations (cell types 0 and 2), and one smaller cluster (cell type 3).

**Fig. 5.**
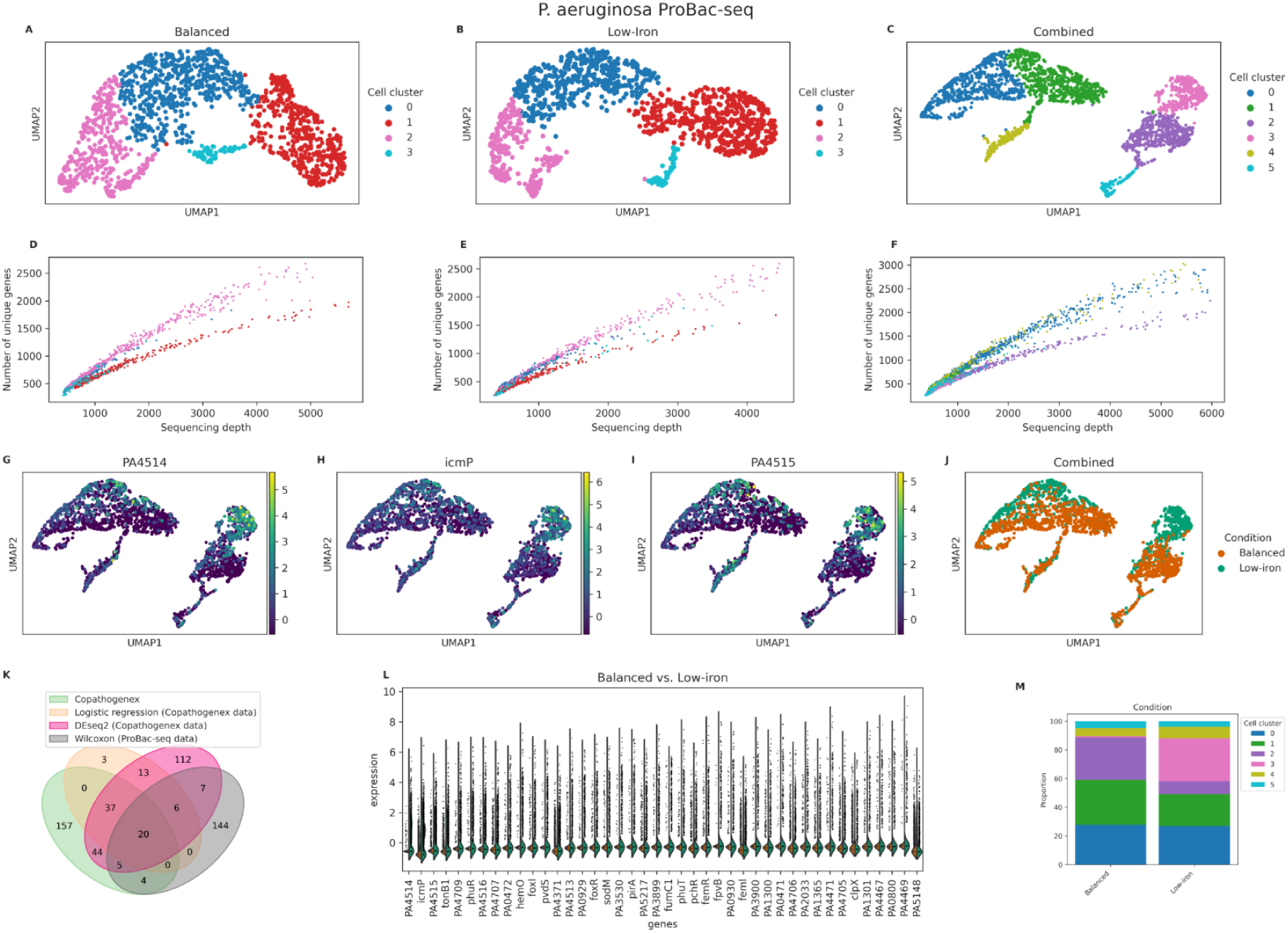
Processing with BacSC discovers a distinct response of *P. aeruginosa* to a low-iron environment. Analysis of the *Pseudomonas_balanced_PB, Pseudomonas_li_PB* datasets and a combination of both with BacSC. **(A-C)** UMAP plots based on the parameters determined by BacSC, colored by cell cluster for the balanced, low-iron, and combined datasets, respectively. **(D-F)** Scatterplots of sequencing depth versus number of unique genes per cell, colored by cell cluster, for all three datasets. **(G-1)** UMAP plots of the combined dataset, highlighting normalized expression values of the three genes most significantly associated with iron reduction (see subfigure L). **(J)** UMAP plot of the combined dataset, highlighting growth condition (balanced or low-iron) for each cell. **(K)** Venn diagram of differentially expressed genes found in Co-PATHOgenex and ProBac-seq data for Pseudomonas in balanced versus low-iron growth conditions. **(L)** Violin plots of differentially expressed genes in ProBac-seq and Co-PATHOgenex (at least one DE method, balanced vs. low-iron). (M) Stacked barplot of cluster proportions for cells from each growth condition.

The lower-coverage cell types in both datasets were characterized by 51 and 82 genes respectively, that were differentially expressed at an FDR of 0.05 (Tables E6, E7) when compared to the rest of the population. Of the 95 genes differentially expressed in either of the two datasets, 38 genes appeared in both, including 22 genes encoding components of the 30S and 50S subunits of the ribosome (*rpsA, rpsB, rplQ, rpsKD, rplFO, rplDWBCP, rpmC, rplEN, rpsJ, rpsG, rplJ, rplK, rpsRI*, Figures C6E-G, C7E-G), indicating increased translation activity. Cell type 3 also showed considerable overlap between DE genes at the 5%-level. Here, all 22 genes that were DE in the balanced growth sample were also among the 34 genes detected in the low-iron culture. Many of these genes encode the R-type pyocin R2 (*PA0617, PA0618, PA0619, PA0620, PA0622, PA0623, PA0640*, Figures C6E-G, C7E-G), a phage tail-like bacteriocin that specifically targets and kills competing bacteria by puncturing their cell membranes [50, 51]. For cell type 2, which contained cells with a large number of expressed genes, a large number of genes was detected to be DE at an FDR of 0.05, with underexpressed ribosomal genes showing the highest contrast scores, complementary to the set of DE genes in the low-coverage cell type. The remaining cell type 0 contained cells with low sequencing depth and showed no statistically significant DE genes.

#### 2.4.2 Combined data processing allows for the detection of genes related to iron acquisition

To analyze the differences between the cell populations from balanced and low-iron growth conditions, we created a combined dataset by concatenating the raw count matrices of both experiments. Processing with BacSC revealed a similar common structure as in the individual datasets (Figures Figure 5C, F, C8A-D), confirming the similarities detected in the previous section. While the R2 pyocin cluster (cell type 5) showed good mixing between both conditions, the cell populations with high expression of ribosomal genes distinctly separated and were even clustered into different cell types (2 and 3, (Figure 5C)). Additionally, a new cell type (cluster 4) emerged in the combined dataset, which was not detected in either of the individual datasets. Similar to cell type 0, this cluster showed reduced expression of ribosomal genes (*rplF, rplP, rplD, rplB*), as well as genes encoding for ATP-synthase and the TCA cycle component succinate dehydrogenase (*atpA, atpD, atpH, sdhA, sdhC*, Figure C8E-G), suggesting a low energy state. For cell types 0 and 1, a within-cluster shift of cells by condition was also visible (Figure 5J). As in the individual data set analyses, marker genes for all cell types except cell type 1 were detected by BacSC at FDR levels smaller than 0.2.

Plotting the cell type proportions for each sample showed that cell types 2 and 3 almost exclusively contained cells from one condition, while the other cell types showed no notable changes in proportionality between the balanced and low-iron conditions (Figure 5M). We confirmed this visual result by differential abundance testing with scCODA [52] and detected cell types 2 and 3 as differentially abundant at an FDR level of 0.2.

Finally, we examined the differences in gene expression between cells from both growth conditions. For this, we first performed DE testing between the balanced growth and low-iron cell populations with a Wilcoxon rank-sum test. Since this test setup does not suffer from double-dipping, we used the Benjamini-Hochberg correction [53] to account for multiple comparisons, revealing 186 genes with corrected p-values of less than 0.05. To verify our findings, we used bulk sequencing results from the Co-PATHOgenex study [54], also testing differential expression between cells grown in balanced and iron-reduced conditions. Of note, in this study an abrupt iron limitation was artificially induced by the addition of the iron chelator 2,2’-bipyridine shortly before harvest. We compared the gene set found by BacSC on the bacterial scRNA-seq data with three gene sets detected on the Co-PATHOgenex data with different DE tests - the method described in the Co-PATHOgenex paper, a logistic regression model, and DESeq2 [55], each at a significance level of 0.05. The gene set from BacSC had good overlap with the gene sets found in bulk data, as 42 of the 186 genes were detected by at least one other DE test, and the intersection of all four gene sets contained 20 genes (Figure 5K). Furthermore, 26 of the 42 genes detected in the bulk data were among the top 50 genes with the lowest adjusted p-values in the DE test on the bacterial scRNA-seq data (Table E3). Investigating the gene expression levels and function of these 42 genes, we found most of them to be overexpressed in the low-iron sample (Figure 5G-I, L). Furthermore, most of these genes (e.g. *PA4514, icmP, phuR*) are known to be related to iron reception (Table E3).

## 3 Discussion

The emergence of protocols for scRNA-seq of bacterial populations is about to transform microbiology research by allowing to evaluate the transcriptional profiles of bacteria at an unprecedented combination of scale and resolution. Despite their technological similarity, bacterial scRNA-seq datasets at their current state differ significantly from eukaryotic scRNA-seq data in terms of sparsity and sequencing depth. To facilitate the statistically sound processing of bacterial scRNA-seq data, we present BacSC, a computational pipeline that allows for easy, dataset-specific quality control and automatic variance stabilization, low-dimensional representation, neighborhood embedding, clustering, and differential expression analysis of such data.

By using a variance-stabilizing transform with gene-wise zero imputation parameters [22], BacSC is able to adequately normalize gene expression data with very large amounts of zero entries and low sequencing depth. We show that train-test splitting through data thinning [28, 33] and comparison to negative control data in scDEED [27] provides ways to select suitable parameters for dimensionality reduction, and neighborhood embedding. Furthermore, selecting a clustering resolution through our newly defined gap statistic based on count splitting of the raw expression data reveals biologically distinct subpopulations. To counteract FDR inflation when testing differential gene expression of bacterial cell types, we extend the ClusterDE method [29] to highly disproportionate cluster sizes. Additionally, our copula-based simulation setup adapts the approach from scDesign [47, 56] to bacterial scRNA-seq data. To this end, we add correlation shrinkage [57, 58] and an adjustment for underestimation of small gene-gene correlations.

Overall, BacSC is a highly flexible framework that performs statistical analysis of bacterial scRNA-seq data independent of the underlying sequencing protocol, while avoiding common statistical pitfalls. Through its capabilities for automated parameter selection, BacSC further allows for a set-and-forget approach to bacterial scRNA-seq data processing, greatly simplifying these tasks. We demonstrated this flexibility through application to 13 bacterial scRNA-seq datasets from two protocols across five different species. Despite large differences in size and sequencing depth per cell even after manual quality control, BacSC was able to integrate, cluster, and perform differential expression testing on each dataset without needing any further user intervention.

The detected cell types and their marker genes showed remarkable overlap with the clusters previously found through processing with default or manually selected parameters in multiple datasets [13, 18], confirming the correctness of BacSC’s findings. BacSC was further able to better depict dynamics between cellular subpopulations in *B. subtilis* and found new bacterial cell types in *K. pneumoniae*. Analyzing two datasets from *P. aeruginosa* grown in environments with different iron availability, BacSC found similar cell types, highlighting its robustness. After joint processing of both datasets with BacSC, differential expression testing correctly detected various genes related to iron acquisition.

Its modular structure and seamless integration in scanpy [37] allow users to easily apply the entire BacSC pipeline or parts of it to their own data, and perform downstream analysis with other methods provided in the scverse [38]. In our studies, we used these capabilities to test for differential abundance between cell type proportions with scCODA [52].

In addition to the described features, there are multiple areas where further improvements and extensions to BacSC are possible. While we developed and evaluated BacSC with bacterial scRNA-seq data in mind, the techniques used were designed for eukaryotic scRNA-seq analysis. Therefore, BacSC is in principal also suited for this type of data, expanding its application range beyond the usecases shown here. In its current state, BacSC uses methods that are seen as the baseline in scRNA-seq analysis [25].

While we adapted these techniques here to fit the properties of bacterial scRNA-seq data, there exist a plethora of approaches, each with their own assumptions, that often show improved capabilities on eukaryotic data [59]. Careful evaluation of these methods in the context of bacterial scRNA-seq requires further efforts.

Finally, our improvements on the synthetic data generation algorithm for differential expression testing currently only cover simulation of one homogeneous cell population. An extension to match the capabilities of scDesign2 and scDesign3 [47, 56] in simulating multiple cell types, batches, trajectories, and spatial information is an open challenge.

By eliminating the need to manually select suitable techniques and parameters, BacSC removes sources of errors and allows for more efficient data processing. We therefore believe that BacSC provides an easily applicable framework that facilitates proper statistical analysis of bacterial scRNA-seq data.

## Supporting information

Key resources table

## Acknowledgments

We thank Sine Lo Svenningsen for providing the *E. coli* strain MAS1081. Furthermore, we thank Petra Hagendorff for her assistance in using the Chromium Controller and Astrid Dröge for her support in preparing sequencing libraries.

C.L.M. acknowledges core funding from the Institute of Computational Biology, Helmholtz Zentrum München. C.L.M. and S.H. received funding from the Deutsche Forschungsgemeinschaft (DFG, German Research Foundation) in the framework of the Priority Program SPP2389 “Emergent functions of bacterial multicellularity” (HA 3299/9-1, AOBJ: 687646). Furthermore, S.H. received funding under Germany’s Excellence Strategy – EXC 2155 “RESIST” – Project ID 390874280, from the Novo Nordisk Foundation (NNF 18OC0033946), from the SFB/TRR-298-SIIRI – Project-ID 426335750 and the Ministry of Science and Culture of Lower Saxony (Niedersächsisches Ministerium für Wissenschaft und Kultur) BacData, ZN3428. R.O.A. and C.L.M. were funded by the StressRegNet consortium within the Bavarian research network bayresq.net funded through the Bavarian State Ministry of Science and Arts, Germany.

## Author contributions

J.O. and C.L.M. designed the structure and individual steps of the BacSC pipeline, and conceived improvements to existing methods. T.K., J.G.T. and S.H. generated data containing *P. aeruginosa* and *E. coli* with ProBac-seq, A.Z.R. provided all datasets from *B. subtilis*. J.O. implemented the pipeline and conducted all applications and tests. J.O., T.K., R.O.A., J.G.T., and A.Z.R. analyzed the results from BacSC in a biological context, R.O.A. further performed analysis of the Co-PATHOgenex data. J.O. wrote the manuscript with help from all other authors. All authors read and approved the manuscript.

## Supplemental information

- Supplemental pdf:

– Additional dataset analysis
– Supplemental figures B1-B5, C6-C17, D18-D31
– Supplemental tables E1-E17
- Pa probes.xslx: Probes used in ProBac-seq of *P. aeruginosa*

## STAR Methods

### RESOURCE AVAILABILITY

#### Lead Contact

Further information and requests for resources and reagents should be directed to and will be fulfilled by the lead contact, Johannes Ostner.

#### Materials Availability

Materials generated in this study are freely available at public repositories (see key resources table) or by contacting the lead contact.

#### Data and Code Availability

Single-cell RNA-seq data have been deposited at GEO and are publicly available as of the date of publication. Accession numbers are listed in the key resources table. Intermediate datasets have been deposited at zenodo and are publicly available as of the date of publication. DOIs are listed in the key resources table. This paper analyzes existing, publicly available data. The accession numbers for these datasets are listed in the key resources table. All original code has been deposited at GitHub (https://github.com/bio-datascience/BacSC) and is publicly available as of the date of publication. DOIs are listed in the key resources table. (additional citations in the key resources table: [60–62])

### EXPERIMENTAL MODEL AND STUDY PARTICIPANT DETAILS

For validating the performance of BacSC, we analyzed previously published scRNA-seq datasets. For ProBac-seq data analysis, we used the *Bsub minmed PB* dataset from the original publication (GEO: GSE223752) [18]. For BacDrop data analysis, we selected seven datasets provided in the original publication [13], and used the read count matrices published by the authors (GEO: GSE180237). The *Klebs BIDMC35 BD*, *Klebs 4species BD*, *Ecoli 4species BD*, *Efaecium 4species BD*, and *Pseudomonas 4species BD* datasets were used as provided. For the *Klebs untreated BD*, and *Klebs antibiotics BD* datasets, we concatenated the count matrices from multiple samples before analysis wth BacSC.

Furthermore, in this study we generated additional datasets using ProBac-seq, encompassing two experiments on *Bacillus subtilis*, two samples of *Pseudomonas aeruginosa*, as well as one sample of *Escherichia coli*.

#### ProBac-seq of B. subtilis

For the *Bsub damage PB* dataset, cells were grown to mid-log phase in spizizen’s minimal media (SMM) and Mitomycin C (MMC, 0.5*µg/ml* final concentration) was added to wildtype *B.subtilis* (strain 168) as reported by [63]. The *Bsub MPA PB* data contains *B.subtilis* cells grown in SMM as described by [64, 65] to mid-log phase and challenged with Mycophenolic acid (MPA, 40*µg/ml* final concentration).

#### ProBac-seq of E. coli and P. aeruginosa

For the samples *Ecoli balanced PB*, *Pseudomonas balanced PB* and *Pseudomonas li PB* MOPS (morpholinepropanesulfonic acid) minimal medium (supplemented with 100 ng/µl thiamine) with 0.2 % glucose as the sole carbon source was used [66]. To induce a mild iron limitation on *Pseudomonas li PB*, the FeSO_4_ concentration was lowered to 0.5 µM instead of the regular 10 µM. Single colonies of *E. coli* MAS1081 [67, 68] and PAO1 were used to inoculate precultures with regular MOPS and were grown for 11-12 hours at 37°C with shaking at 180 rpm. After washing, main cultures in MOPS with normal iron or reduced iron content were inoculated at an OD_600_ of 0.00002 and grown for 10-14 generations. Bacteria were harvested in balanced growth conditions in early exponential phase (OD_600_ of 0.2-0.3).

### METHOD DETAILS

#### ProBac-seq of B. subtilis

For all *B. subtilis* datasets, ProBac-seq was performed as described in the original method [18, 36].

#### ProBac-seq of E. coli and P. aeruginosa

Further sample preparation for ProBac-seq was performed as previously described [18, 36], with slight modifications. In brief, 1 ml of each culture was used for fixation with 1 % formaldehyde for 30 min at room temperature. To increase the cell yield, all centrifugation steps were carried out at 7,000 x g for up to 5 min. Overnight storage in MAAM (4:1 V:V dilution of methanol to acetic acid) was omitted. All further steps were performed according to the protocol of the original method [18, 36]. PAO1-specific probes were designed and generated as previously described without additional UMI extension. The single-cell sequencing libraries were quality-checked and sequenced by the GMAK sequencing facility (HZI, Braunschweig, Germany) on a NovaSeq SP flow cell (100 cycles, 28-10-10-90) resulting in up to 170 million reads per sample. Raw fastq files were processed with CellRanger v7.1.0 [69] with the option *–expect-cells 10000*.

### QUANTIFICATION AND STATISTICAL ANALYSIS

This section describes statistical details for the individual steps in the BacSC pipeline. Statistical details and results from application of the BacSC pipeline to all datasets described in table 1 can be found in supplementary figures D18-D31 and supplementary tables E1-E17.

Processing starts with a raw counts matrix *X*_0_ ∈ N*^n^*^0^ *^×p^*^0^, which contains read counts of *p*_0_ genes for *n*_0_ droplets.

#### Quality control

For datasets generated with ProBac-seq, multiple probe reads for each gene are available. As described in the original publications [18, 36], we aggregated the probes by max-pooling. Furthermore, most datasets from ProBac-seq were already quality-controlled in CellRanger [69] and therefore needed less additional filtering. For all ProBac-seq datasets, we chose a minimum sequencing depth cutoff of 100. For data from BacDrop, we used the minimum sequencing depth cutoff of 15, as provided in the original publication [13]. For the three largest datasets (*Klebs untreated BD, Klebs antibiotics BD, Klebs BIDMC35 BD*), we also selected 2,500 highly variable genes after variance stabilization. BacSC further removes genes that were expressed in only a single cell, as variance stabilization for these genes is not possible. In contrast to eukaryotic scRNA-seq datasets, removal of mitochondrial genes is not required for bacterial scRNA-seq, as bacteria do not contain mitochondria. Still, other highly abundant types of RNA, such as rRNA and tmRNA, can be removed at this point. For the analysis presented here, we did not perform any removal of features beyond the preprocessing in CellRanger [69] for ProBac-seq or UMI-tools [70] for BacDrop.

Further outliers are detected by filtering cells based on median absolute deviations (MAD) of their log-transformed total counts and number of expressed genes [30]: *MAD*(*S*) = *median^n^_i=1_* (| log(*S_i_*) − *median* (log(*S*))|) where *S* is either the vector of sequencing depths 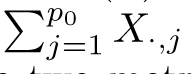 or number of expressed genes over all cells. A cell is considered an outlier if for either of the two metrics, |*S_i_* − *median*(*S*)| *> nmads* ∗ *MAD*(*S*), where *nmads* is the factor defined in table E2.

Table E2 gives an overview over the filtering parameters chosen for each dataset. After filtering, *X*_0_ is reduced to a matrix *X* ∈ N*^n×p^* of *p* genes and *n* cells.

#### Variance stabilization

For variance-stabilizing transformation (VST) of the filtered read counts, we follow the results from [22]. Assuming potential overdispersion of the count distribution, we use an approximation to the ideal VST determined by the delta method, a log-transformation in combination with common-sum scaling of the counts:

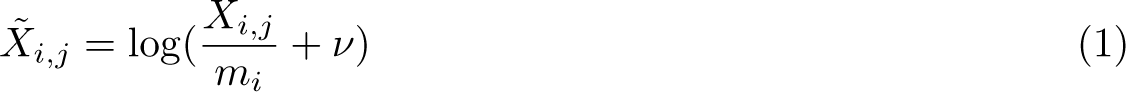

where 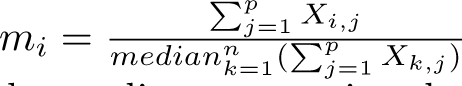 scales each cell’s counts to the median value of all sequencing depths. We chose the median sequencing depth as a scaling factor to gain robustness to outliers in sequencing depth.

Adding a pseudocount *ν* before log-transformation is necessary to handle zero entries in *X*. As described in [22], we set 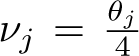 for each gene *j* = 1 *. . . p*, where *θ_j_* denotes the gene’s overdispersion factor. Calculating this overdispersion factor is not straightforward for genes with very low numbers of *mean*(*X*·*,j*)^2^ expressed genes, as the relation 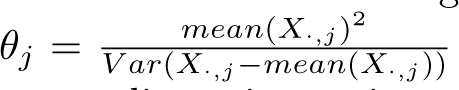 becomes very sensitive to single entries in *X*.

Instead, we make use of the gene overdispersion estimates provided by sctransform [41], which jointly models all genes, and thus produces more robust estimates of *θ_j_*. To this end, we apply sctransform to the count matrix *X*, extract the overdispersion estimates, and use them in equation 1.

After VST, we scale each gene individually to zero mean and unit variance by applying scanpy’s *scale* function [37], clipping large values at 10. This results in a normalized gene expression matrix *Y* ∈ R*^n×p^*.

#### Dimension reduction

The selection of the best embedding dimensionality *k_opt_* through data thinning was described for Poisson-distributed data in [31]. There, data thinning [33] is used to split the raw count data *X* into two *n* × *p*-dimensional datasets *X^train^* and *X^test^* by a random binomial split on each individual entry in *X*. The resulting train and test matrices are then both Poisson-distributed again. Because eukaryotic single-cell data is typically assumed to follow a Negative Binomial (NB) distribution for each gene, [28] extended the data-thinning approach to NB-distributed data. However, the lower read counts in bacterial scRNA-seq suggest that the data might follow a linear instead of a quadratic mean-variance pattern and are therefore Poisson-distributed.

To determine the distributional assumption for count splitting, we first calculate the mean *µ_j_* and variances *σ*^2^ of *X_·,j_* for each gene *j* = 1 *. . . p*. We then compare Pearson correlation coefficients *r* of a linear and a quadratic relation between *µ* and *σ*^2^. If *r_quadratic_ > r_linear_*, we assume *X* to be Negative Binomial distributed, otherwise it is Poisson-distributed. The raw data distribution for each dataset is shown in Table E2.

Depending on the chosen data distribution, *X* is split into two datasets by Poisson or NB count splitting (Figure B1A, B). In both cases, we set the split ratio *ɛ* = 0.5 to ensure an even split between train and test data and maximize the probability of obtaining a nonzero entry in train and test data if *X_i,j_ >* 1. We then determine all genes or cells that have only one nonzero entry in *X_train_* or *X_test_*, and remove them from both data splits. In line with [31], we apply the VST described in section 3 to both *X_train_*and *X_test_*, using the *θ* parameters determined on the whole data to speed up computation, and obtain transformed matrices *Y_train_* and *Y_test_*.

To determine *k_opt_*, we perform a singular value decomposition (SVD) *Y_train_* = *U* Σ*V ^T^* on the training data. For each *k* = 1 *. . .* 20, we then calculate the reconstruction loss as sum of squared differences between the test data and the *k*-dimensional approximation of the SVD of the train data (Figure B1C):

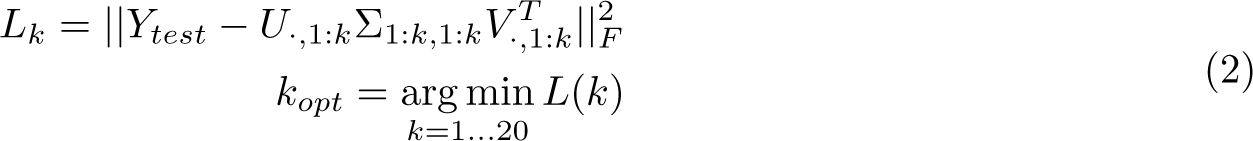

#### Data visualization

BacSC selects the latent parameters *n_neighbors_* and *min_dist_* for constructing a UMAP embedding of the data through scDEED [27]. For every combination of *n_neighbors_* (the number of neighbors for each cell in the neighborhood graph) and *min_dist_* (the effective minimum distance between points), scDEED defines a reliability score for each cell as the Pearson correlation between the euclidean distances to the 50% closest cells in PCA space and the euclidean distances to these cells after UMAP embedding. To obtain a baseline distribution, another set of reliability scores is calculated on a permuted dataset where each gene’s expression values are shuffled. scDEED then classifies the embedding of cells in the original dataset as ”trustworthy”, ”undefined”, or ”dubious” based on the 95% and 5% quantiles of the distribution of reliability scores in the permuted data (Figure B1D). Finally, the parameter combination with the smallest number of dubiously embedded cells is selected (Figure B1E, F).

As scDEED is only available in R, the BacSC pipeline includes a Python implementation of the method. For every dataset, we considered all pairwise combinations of parameters: *n_neighbors_* : (10, 15, 20, 25, 30, 35, 40, 45, 50, 60, 70, 80, 90, 100, 150, 200, 250); *min_dist_* : (0.05, 0.1, 0.3, 0.5, 0.7).

#### Clustering

The resolution parameters in Louvain and Leiden clustering are essential for defining the granularity of the resulting partition [44, 71]. Both algorithms aim to optimize the modularity or a similar metric of a partition on the neighborhood graph defined during UMAP generation:

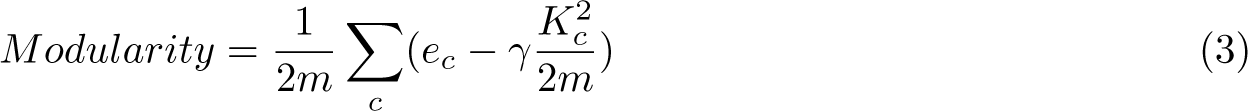

where *m* is the total number of edges in the neighborhood graph, *e_c_* is the number of edges within cluster *c*, and *K_c_* is the sum of degrees over all nodes in cluster *c*, and *γ* is the resolution parameter. Generally, a higher resolution parameter will lead to a more fine-grained clustering. While both algorithms effectively approximate an optimal clustering for a given value of *γ*, the choice of a ”good” resolution parameter is highly dependent on the structure and biological source of the data at hand [32, 72]. Larger datasets or datasets from more complex communities generally contain more subclusters and thus warrant a larger value of *γ* to detect all relevant subpopulations. On the other hand, choosing the resolution too large will result in non-robust clusterings that are highly sensitive to small perturbations of the data [73]. Furthermore, cluster assignments and the number of subpopulations are not monotonic in *γ*, complicating the evaluation of clustering quality [32]. In BacSC, we aim to automatically find a resolution parameter that results in an informative, but stable clustering of the cells.

To this end, we adapt the idea from [28] and use the train and test datasets obtained through count splitting for clustering evaluation. Starting with the variance-stabilized train and test data from dimensionality reduction, we generate the neighborhood graph for both datasets with the *k* and *n_neighbors_* parameters determined earlier. For each value of *γ* in a set of possible resolutions, we then perform Leiden or Louvain clustering on the training data, resulting in a cluster assignment *c_train_*. Since training and test data contain the same cells, we can now obtain a measure for the robustness of the clustering by calculating the modularity (3) for *c_train_* on the neighborhood graph of the test data (Figure B2A). We denote this value with *M_test_*. Since modularity generally decreases with the number of clusters, we cannot select the value of *γ* for which *M_test_* is maximal. Instead, we need to compare the test data resolution to a baseline score for each resolution value. Therefore, we generate a random cluster assignment on the test data by permuting the labels from *c_train_* and calculate *M_random_*, the modularity of the random clustering on the neighborhood graph of the test data. Finally, we select the resolution where the gap statistic between test modularity and random modularity is maximal (Figure B2B, C):

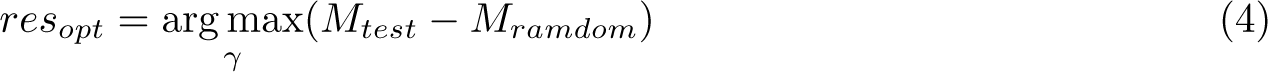

and perform a clustering with *res_opt_* on the full dataset to obtain cell type clusters (Figure B2D). For processing the datasets in this manuscript, we used the Leiden algorithm and modularity score and tested possible resolutions *γ* = (0.01, 0.03, 0.05, 0.49). The same procedure is however also applicable to Louvain clustering or other measures, e.g. the Constant Potts model [74].

Even though the resolution value determined by maximizing our gap statistic provides improvement over random cluster assignment while being robust to small data perturbations, it is by no means the only ”correct” resolution value. For some datasets, more fine-grained clusterings can give further insights into subpopulations of the data. Rather, *res_opt_*may serve as a baseline clustering resolution that gives an adequate first insight into the data.

#### Differential expression testing

Identifying genes with characteristic expression for cell clusters defined by the same gene expression values is an instance of reusing information, or ”double dipping” [46], and controlling the false discovery rate under such conditions is essential to achieve adequate results. The ClusterDE method [29] provides FDR control for DE testing of cell types in eukaryotic scRNA-seq by contrasting the p-values of interest with p-values calculated on a synthetically generated negative control dataset. In BacSC, we implement a modified version of the algorithm that takes the characteristics of bacterial single-cell data into account and allows for testing of highly disproportionate cell populations. The following description assumes a DE test of cell type *C* with *n_C_* cells against the union of all other cell types, containing *n_C̄_* = *n* − *n_C_* cells (Figure B3A). Tests of differential gene expression between two cell types are possible in the same manner, but the data needs to be subsetted to the clusters of interest first.

ClusterDE first generates negative control data with the same marginal gene distributions and genegene correlations as the original data, but no intrinsic cluster structure. This synthetic data generation is done with scDesign2 [47] or scDesign3 [56], which both use a Gaussian copula approach to generate synthetic scRNA-seq data. To account for the high sparsity and low sequencing depth of bacterial scRNA-seq data, we adapted the approach from scDesign2 in BacSC. In a first step, the marginal distribution of raw counts is determined for every gene *j*. As in scDesign2, we consider four possible distributions - Poisson (Poi), zero-inflated Poisson (ZIP), Negative Binomial (NB), and zero-inflated Negative Binomial (ZINB). If the gene’s empirical variance *σ*^2^ is larger than its empirical mean *µ_j_*, we determine the gene to be NB-or ZINB-distributed, otherwise its distribution is Poi or ZIP. We then fit the Poisson or NB distribution with and without zero-inflation to *X_·,j_* through maximum likelihood estimation via BFGS, as implemented in the *statsmodels* package [75]. Because of the large number of zeros, we experienced frequent convergence problems with NB estimation. To counteract this, we set the initial mean and dispersion parameters for both NB and ZINB to the mean and dispersion of all nonzero entries in *X_·,j_*, and the initial zero inflation in the ZINB model to the proportion of zeros in *X_·,j_*. If both the NB and ZINB models still do not converge, we instead use the estimates from the NB model with default starting parameters, regardless of convergence. We then perform a likelihood-ratio test between the log-likelihoods of the zero-inflated and regular model. If the null hypothesis of no difference in log-likelihood between both models is rejected at the *α* = 0.05 level, we model the gene with zero-inflation, otherwise we use the non-zero-inflated estimate. Denote the chosen distribution for gene *j* with its estimated parameters as as *D_j_*(*ϕ_j_*)

As in scDesign2, we now transform the discrete counts for each gene to continuous quantiles through a uniform approximation with the corresponding cumulative distribution function (CDF) *D*^^^*_j_* (*ϕ_j_*):

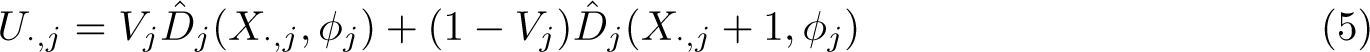

with *V_j_* ∼ *Uniform*(0, 1)*^n^*. We then transform these quantiles by the inverse CDF (denoted Φ*^−^*^1^) of a standard normal distribution and calculate their empirical correlation matrix *R* ∈ R*^p×p^*.

Contrary to eukaryotic scRNA-seq, where current datasets contain many more cells than genes, most of our bacterial scRNA-seq data is underdetermined, with *n < p* (Table E1). Therefore, the entries of the empirical covariance matrix must be shrunk to obtain a good estimate for *R* [57, 58]. To this end, we use a Python reimplementation of the covariance shrinkage proposed in [76].

The uniform approximation 5 in the copula transformation is necessary to allow the use of Gaussian copula for discrete count data, but shifts the count matrix by an average of 0.5. Since bacterial scRNA-seq data contains mostly zero or very small entries, this leads to considerably lower gene-gene correlations and gene variances in the generated data. To counteract this, we introduce a scaling factor *δ* on off-diagonal entries of *R* where the absolute absolute value of the original data’s gene-gene correlation *S* is larger than 0.1:

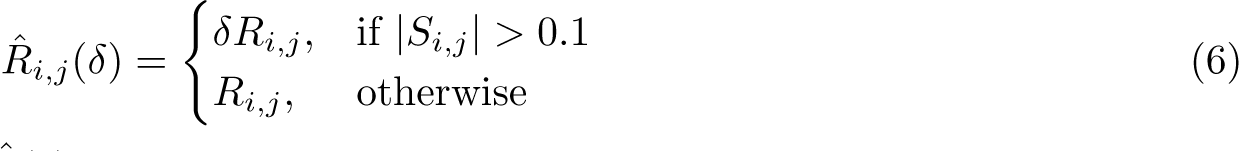

The scaled correlation matrix *R*^^^(*δ*) is not guaranteed to be positive definite though. To obtain a positive definite matrix *R*^∼^(*δ*) that is close to *R*^^^(*δ*), we calculate the eigendecomposition (*λ, v*) of *R*^^^(*δ*), increase all eigenvalues by −*λ_min_* + 10*^−^*^12^ if the smallest eigenvalue *λ_min_*is negative, and set *R*^∼^(*δ*) = *v diag*(*λ*^∼^) *v^−^*^1^ with the shifted eigenvalues *λ*^∼^. We then determine the ideal *δ* through a golden ratio optimizer [77] with initial bracket (1, 2) that minimizes the sum of squared differences between the scaled entries of *R*^∼^(*δ*) and *S*:

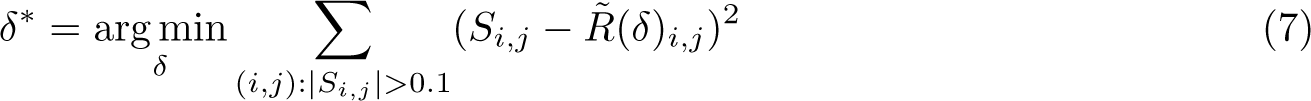

Scaling of the entries in *R* will slightly overestimate the gene means of the generated data (Figure B3B), but gives better results for large gene variances and gene-gene correlations (Figure B3C, D). To simulate synthetic null data with *n^′^* samples and no apparent cluster structure, we generate *n^′^* samples *Z*^^^ from a *Normal*(0*, R*^∼^(*δ^∗^*) distribution, and transform them back into the original space by the standard normal CDF and the inverse CDF of *D_j_*(*ϕ_j_*):

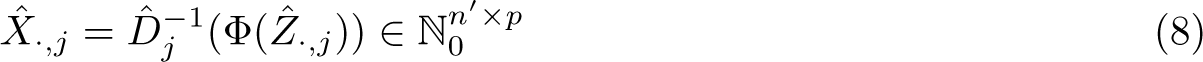

Using this procedure, we can obtain a synthetic null dataset with marginal distributions and gene-gene correlations similar to the target data, but no cluster structure. To allow for generation of negative control data that has the same numbers of cells in both groups as the original data, we set *n^′^* = 2*n* and subset *X*^^^ after processing. Analogous to ClusterDE, we process the synthetic null data in the same way as the original data. We use the same parameters for dimension reduction and neighborhood embedding as determined for the target data, but re-run sctransform on the null data to get new estimates for the gene-wise overdispersion *θ*. By finding a suitable resolution for the Leiden algorithm, we cluster *X*^^^ into exactly two parts, and randomly draw *n_C_* and *n_C̄_* cells from both clusters, respectively (Figure B3E).

FDR control in ClusterDE and BacSC is performed through contrast scores and the Clipper method [35]. We first obtain two sets of *n* p-values by performing the same DE test (e.g. Wilcoxon rank-sum) on the original data and on the drawn subset of the synthetic null data (Figure B3F, G). Next, we calculate the contrast score

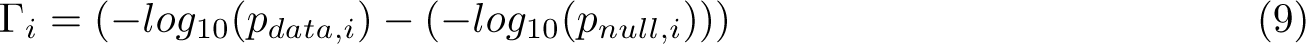

for each pair of p-values. Given a FDR level *α*, Clipper then finds a threshold *T* on the contrast scores

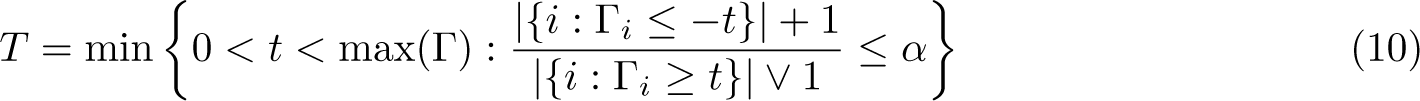

For genes with Γ*_i_ > T*, the expected FDR is less than *α* [34] and we denote them as differentially expressed (Figure B3H).

While differential expression testing with contrast scores is not computationally intensive, the generation of synthetic null data does require some computational power. Fortunately, a series of tests of each cell type’s gene expression against the union of all other cell types only requires generation of the synthetic null data once, as the same set of cells is included in every test and therefore marginal gene distributions and correlations are identical. Only the selection of *n_C_* and *n_C̄_* cells from *X*^^^ and subsequent steps have to be performed individually for each cell type.

## Appendix A Additional dataset analysis

This section contains biological interpretation of selected datasets that were not discussed in the main text.

### A.1 BacSC reveals effects of DNA damage in *B.subtilis*

One more impression on how external factors can change the composition of bacterial cell types is provided by the *Bsub damage PB* dataset by comparing this data to the same species grown in minimal media without DNA damage. First, the PCA plot of the DNA-damaged population did not exhibit the characteristic separation into three subpopulations as observed in the *Bsub minmed PB* dataset (Figure C10A). Instead, the UMAP embedding showed a much more homogeneous population structure C10B) with six different subclusters, and one separate cell type (cluster 6).

This cell type again contained competent cells, as indicated by an overexpression of *com* genes (FDR=0.1, Figure C10E, F, Table E10) although in a much lower concentration than in the experiment without DNA damage (0.9% vs. 9.4% of analyzed population). For cell types 1 and 2, BacSC found many genes to be up- or downregulated, respectively, at an FDR level of 0.1. Cell type 4 showed an overexpression of genes related to subtilosin A production (*albE, albF, albC, albA, albD*), while cell types 3 and 5 showed an overexpression of genes related to the SPbeta prophage (*yomS, yomP, yomR, …*), and prophage PBSX (*xtmA, xtmB, xkdE, xkdC, etc.*), albeit only at FDR levels larger than 0.5.

### A.2 BacSC discovers a new cell type in *K. pneumoniae*

The *Klebs untreated BD* data contains 48,511 cells after quality control and is thus the largest experiment of our analyzed datasets, but also one of the most sparse (99.1% zero entries, Table E1). The PCA plot generated by BacSC (Figure C12A) showed a separation of many cells that were later clustered as cell type 1 (Figure C12B). This cell type showed higher sequencing depth (Figure C12D) and a larger number of unique expressed genes per cell on average (Figure C12C).

Clustering revealed three distinct subpopulations (Figure C12B). Cell type 1 showed a distinct set of genes that were upregulated at an FDR of 0.05 (Figure C12E, G; Table E12). This cell type comprised 2,194 cells and was characterized by IS903B transposase genes (*RS22855*, Figure C12F). This MGE subpopulation was already described in the original publication, but separated more clearly from the rest of the population in the UMAP generated by BacSC (Figure C12B).

Cell type 0 made up the bulk of the cell population (44,236 cells) and was distinguished from the other cell types by no expression of IS903B transposase genes. The analysis with BacSC also found another cell type (Cluster 2), which was not described by [13]. Similar to the high-ribosomal cell type discovered in *P. aeruginosa*, this subpopulation was mostly characterized by a higher expression of ribosomal genes (*rplP, rplC, rpoC*).

## Appendix B Supplementary figures

**Fig. B1.**
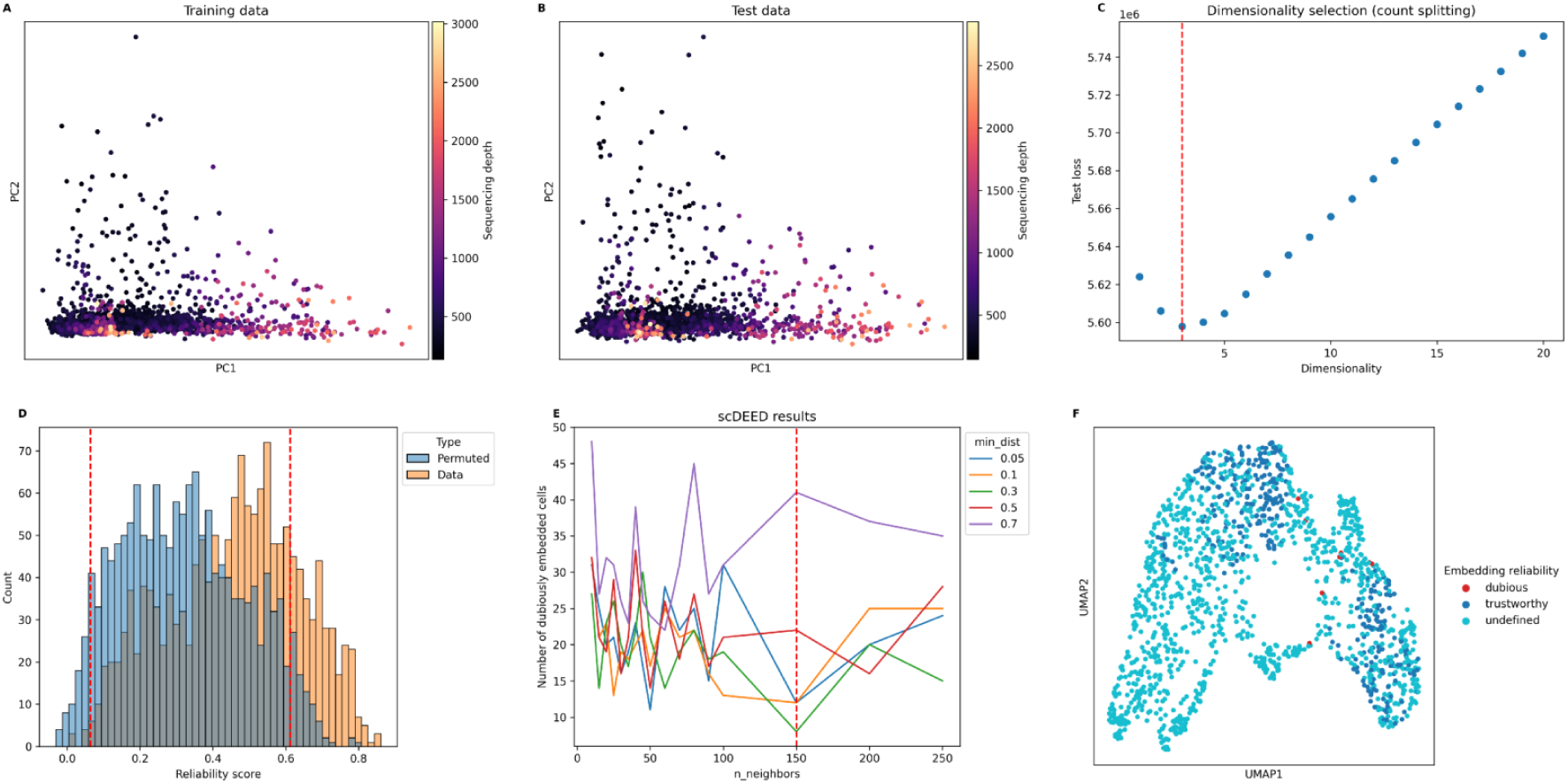
Dimensionality reduction techniques in BacSC. All plots were generated for the *Pseudomonas_balanced_PB* dataset. Count splitting generates a training dataset **(A)** and test dataset **(B)** with similar PCA embeddings and count distribution. **(C)** The latent dimensionality *kopt* of the dataset (dashed red line) is determined by minimizing the test loss. **(D)** Histogram of Null and target data reliability scores from scDEED. The dashed red lines denote the 5% and 95% quantiles of the distribution of null reliability scores *(nneighbors* = 150, *mindist* = 0.3). Cells with reliability scores smaller than the 5% quantile are marked *as* dubiously embedded, cells with reliability scores larger than the 95% quantile are marked *as* reliably embedded. **(E)** Number of dubiously embedded cells for each parameter combination tested **in** scDEED. The dashed red line indicates the chosen parameters *nneighbors* = 150, *mindist* = 0.3. **(F)** UMAP of the full dataset with parameters selected as in (E) and cells colored by their reliability classification from (D).

**Fig. B2.**
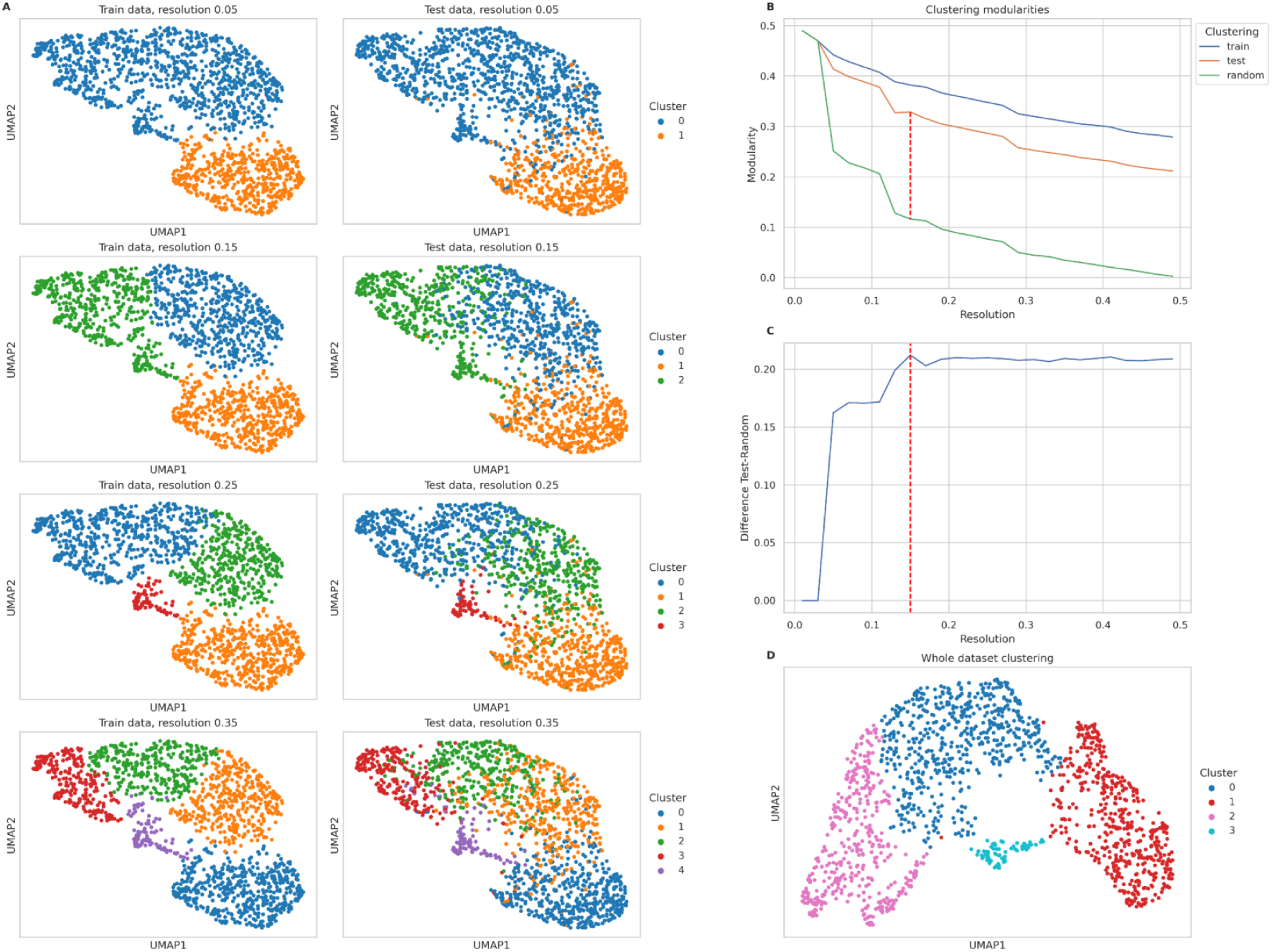
Selection of clustering resolution in BacSC. All plots were generated for the *Pseudomonas_balanced_PB* dataset. **(A)** Clustering on train data (left column) for different resolutions, and applied to test data (right column). **(B)** Modularity scores of train data clustering on train data (blue), test data (orange), and of randomly shuffled clustering on test data (green) for all tested resolutions. The dashed line indicates the largest value of the gap statistic between test and random resolution. This resolution value is selected by BacSC. **(C)** Gap statistic between test and random resolution for all tested values of the resolution parameter. The dashed line indicates the chosen resolution *res opt•* **(D)** UMAP of full dataset, clustered with the resolution parameter determined in (C) and (D).

**Fig. B3.**
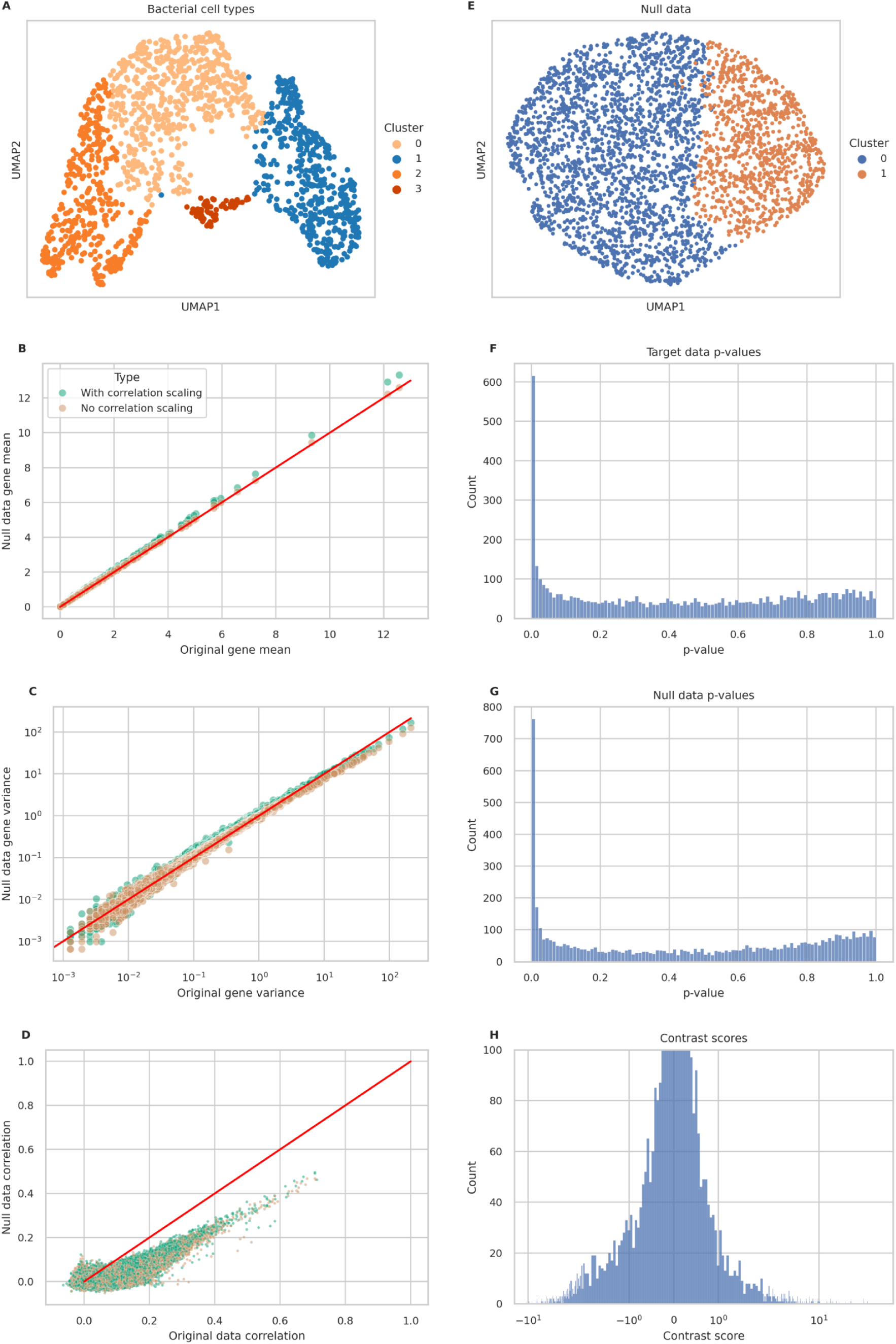
Differential expression testing in BacSC. All plots were generated for the *Pseudomonas_balanced_PB* dataset. **(A)** UMAP of the target data. In this figure, the testing of cluster 1 (blue) against the union of all other clusters (orange) is shown. **(B)** Comparison of gene means for synthetic null data with and without correlation scaling to the original data. The red line indicates a perfect match. **(C)** Comparison of gene variances for synthetic null data with and without correlation scaling to the original data. The red line indicates a perfect match. **(D)** Comparison of empirical gene-gene correlations (shrunk by the procedure outlined in [76]) for synthetic null data with and without correlation scaling to the original data. Only a random subset of 100,000 correlations is shown for each type of synthetic data. The red line indicates a perfect match. **(E)** UMAP of processed null dataset with clustering into two subsets. **(F)** Histogram of p-values for testing cell type 1 against all other cell types on the original (target) da (G) Histogram of p-values for testing cell type 1 against cell type O on the synthetic null data. **(H)** Histogram of contrast scores for testing cell type 1 against cell type 0. The y-axis was truncated at 100.

**Fig. B4.**
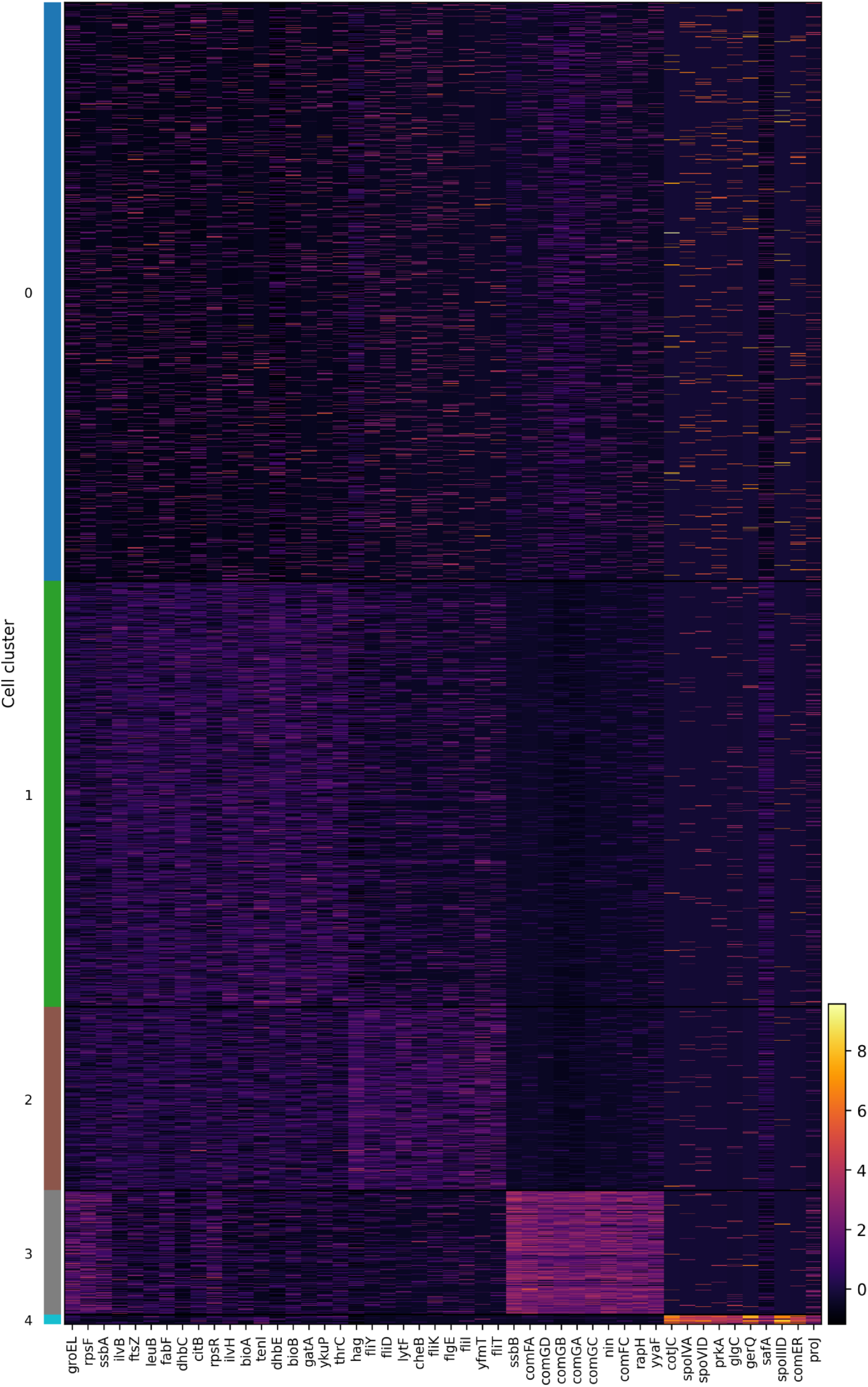
Full heatmap of normalized gene expression for the *Bsub_minmed_PB* dataset. This figure extends Figure 3E by showing all cells. For each cluster, the 10 genes with the highest contrast scores are shown.

**Fig. B5.**
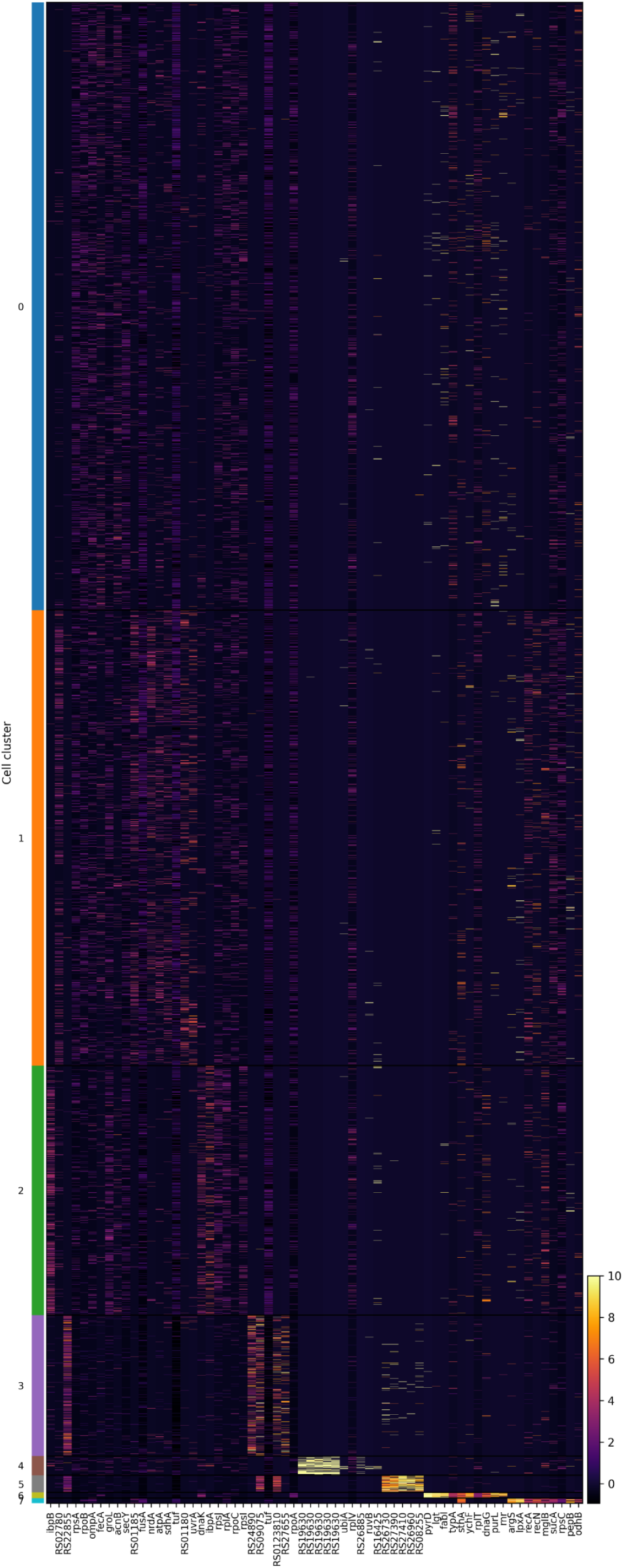
Full heatmap of normalized gene expression for the *Klebs_antibiotics_PB* dataset. This figure extends Figure 4E by showing all cells. For each cluster, the 10 genes with the highest contrast scores are shown.

## Appendix C Additional dataset analysis

This section contains results from BacSC in the style of figures 3 and 4 for all datasets shown in table 1 that were not already shown the main text.

**Fig. C6.**
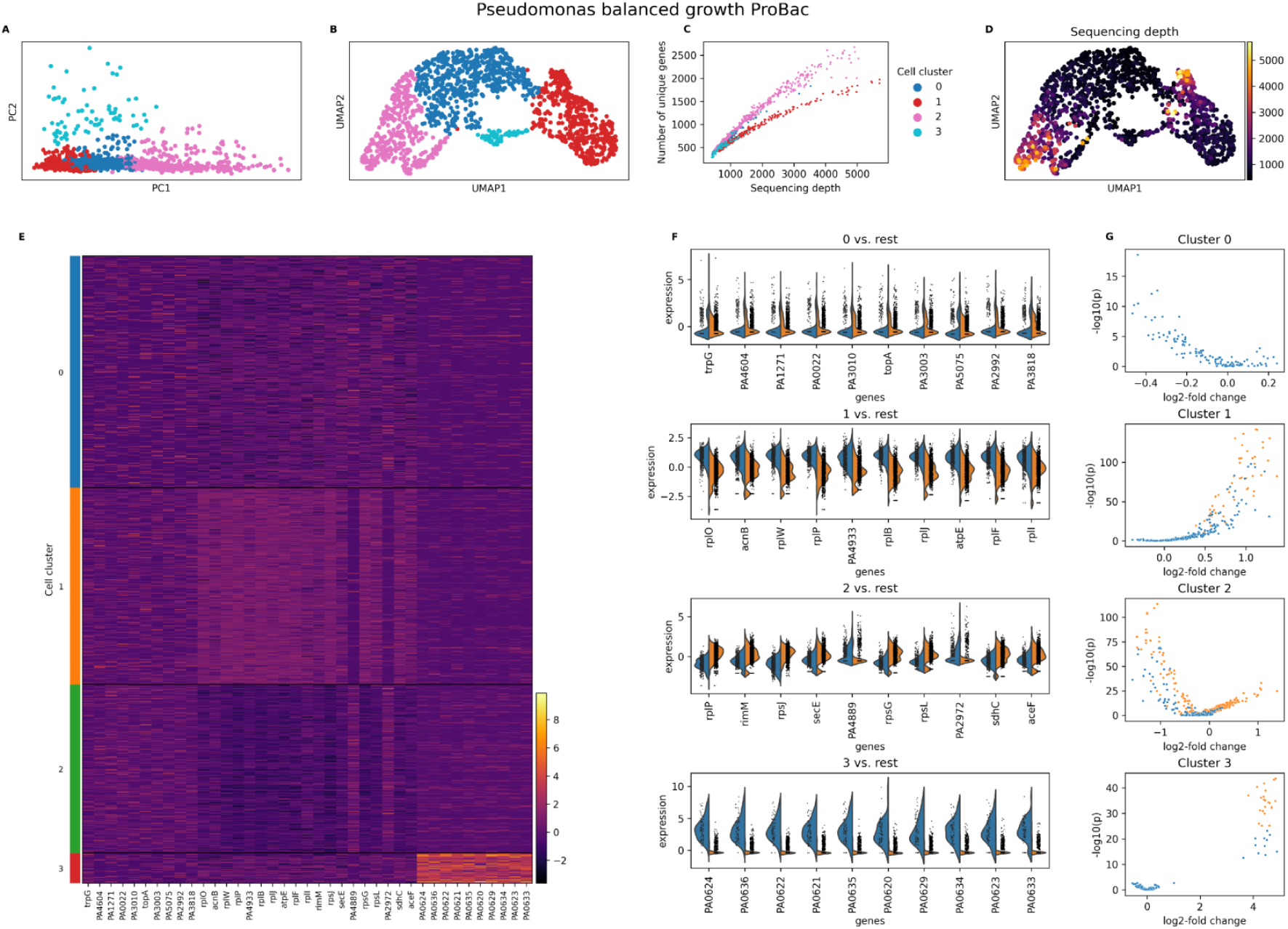
Analysis of the *Pseudomonas_balanced_PB* dataset with BacSC. **(A)** Scatterplot of first two dimensions of PCA embedding with cell type clusters highlighted. **(B)** UMAP plot based on the parameters determined by BacSC, colored by cell cluster. (**C)** Scatterplot of sequencing depth versus number of unique genes per cell, colored by cell cluster. (D) Umap plot as in (B), colored by sequencing depth. **(E)** Heatmap of normalized gene expression for all cells and characteristic genes for cell types. For each cluster, the 10 genes with the highest contrast scores are shown. **(F)** Violin plots of normalized gene expression for DE tests of each cell type (blue) against the rest of the cell population (orange). For each cluster, the 10 genes with the highest contrast score are shown. Genes in (E) and (F) are annotated with gene symbols wherever possible, otherwise locus tags are shown. **(G)** Volcano plots for DE tests as in (F). The x-axis shows the log-fold change for gene expression, the y-axis shows the - log_10_-transformed (uncorrected) p-value. Only genes that are expressed in at least 80% of cells in the respective cluster are shown. Orange genes are differentially expressed at the *a* = 0.05-level.

**Fig. C7.**
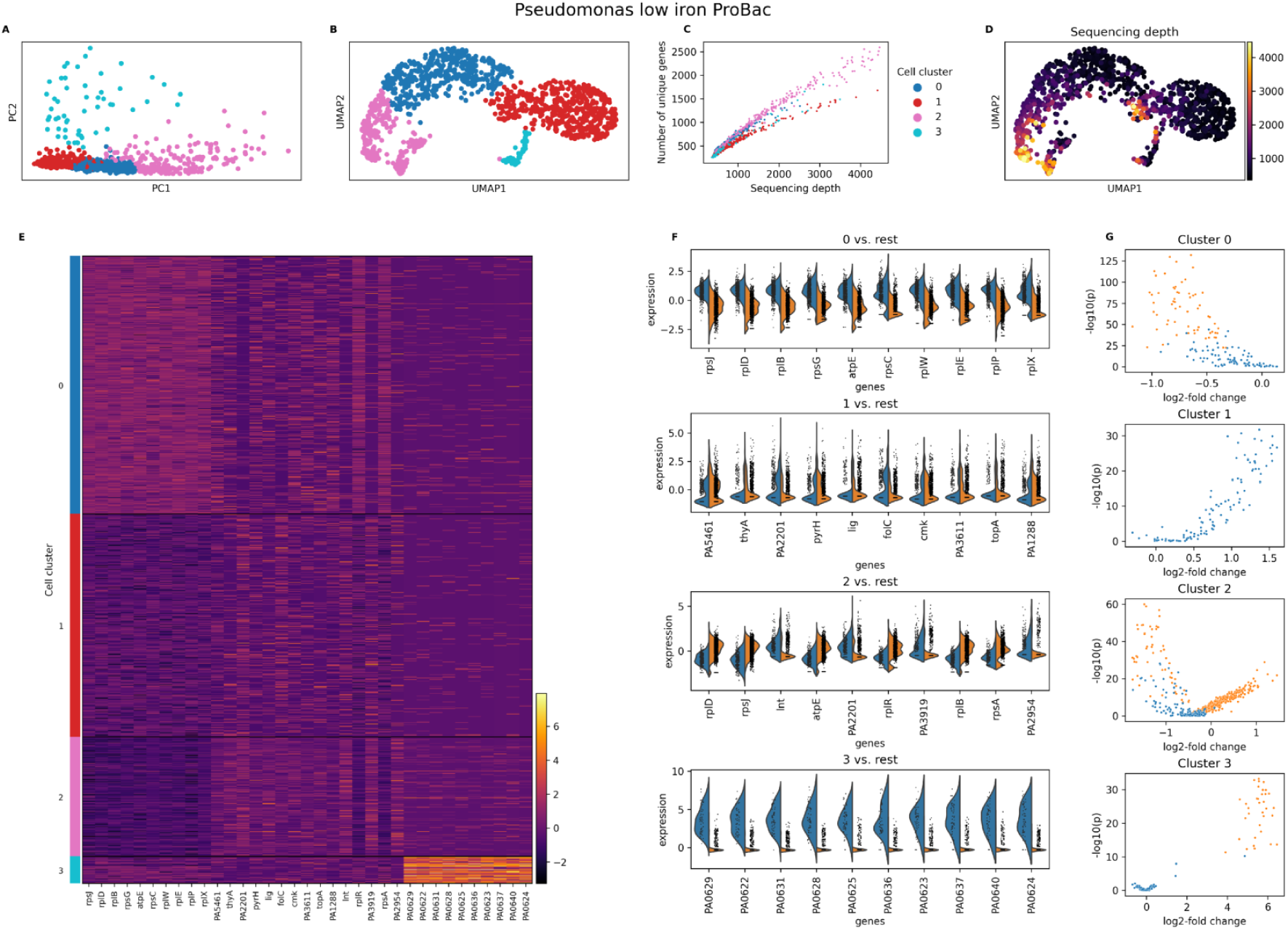
Analysis of the *Pseudomonas_li_PB* dataset with BacSC. **(A)** Scatterplot of first two dimensions of PCA embedding with cell type clusters highlighted. **(B)** UMAP plot based on the parameters determined by BacSC, colored by cell cluster. **(C)** Scatterplot of sequencing depth versus number of unique genes per cell, colored by cell cluster. **(D)** Umap plot as in (B), colored by sequencing depth. **(E)** Heatmap of normalized gene expression for all cells and characteristic genes for cell types. For each cluster, the 10 genes with the highest contrast scores are shown. **(F)** Violin plots of normalized gene expression for DE tests of each cell type (blue) against the rest of the cell population (orange). For each cluster, the 10 genes with the highest contrast score are shown. Genes in (E) and (F) are annotated with gene symbols wherever possible, otherwise locus tags are shown. **(G)** Volcano plots for DE tests as in (F). The x-axis shows the log-fold change for gene expression, the y-axis shows the - log_10_-transformed (uncorrected) p-value. Only genes that are expressed in at least 80% of cells in the respective cluster are shown. Orange genes are differentially expressed at the *a* = 0.05-level.

**Fig. C8.**
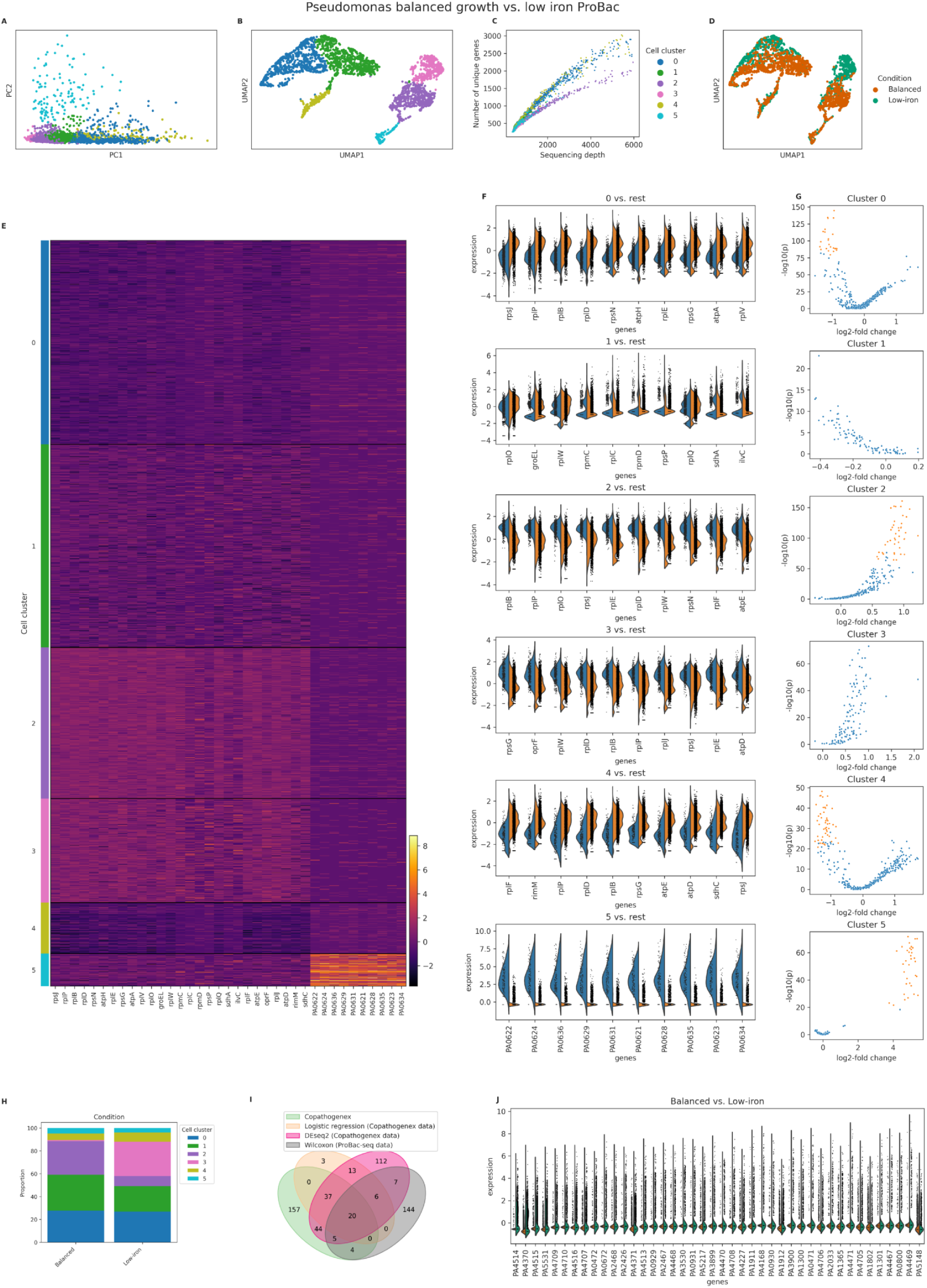
Analysis of the combined *Pseudomonas_balanced_PB* and *Pseudomonas_li_PB* dataset with BacSC. **(A)** Scatterplot of first two dimensions of PCA embedding with cell type clusters highlighted. **(B)** UMAP plot based on the parameters determined by BacSC, colored by cell cluster. **(C)** Scatterplot of sequencing depth versus number of unique genes per cell, colored by cell cluster. **(D)** UMAP plot as in (B), colored by sample (growth condition). **(E)** Heatmap of normalized gene expression for all cells and characteristic genes for cell types. For each cluster, the 10 genes with the highest contrast scores are shown. **(F)** Violin plots of normalized gene expression for DE tests of each cell type (blue) against the rest of the cell population (orange). For each cluster, the 10 genes with the highest contrast score are shown. Genes in and (F) are annotated with gene symbols wherever possible, otherwise locus tags are shown. (**G)** Volcano plots for DE tests as in (F). The x-axis shows the log-fold change for gene expression, the y-axis shows the - log_10_-transformed (uncorrected) p-value. Only genes that are expressed in at least 80% of cells in the respective cluster are shown. Orange genes are differentially expressed at the *a* = 0.05-level. **(H)** Stacked barplot of cluster proportions for cells from each growth condition. (I) Venn diagram of differentially expressed genes found in Co-PATHOgenex and ProBac-seq data for Pseudomonas in balanced versus low-iron growth conditions. (J) Violin plots of differentially expressed genes in ProBac-seq and Co-PATHOgenex (at least one DE method, balanced v 3’tow-iron).

**Fig. C9.**
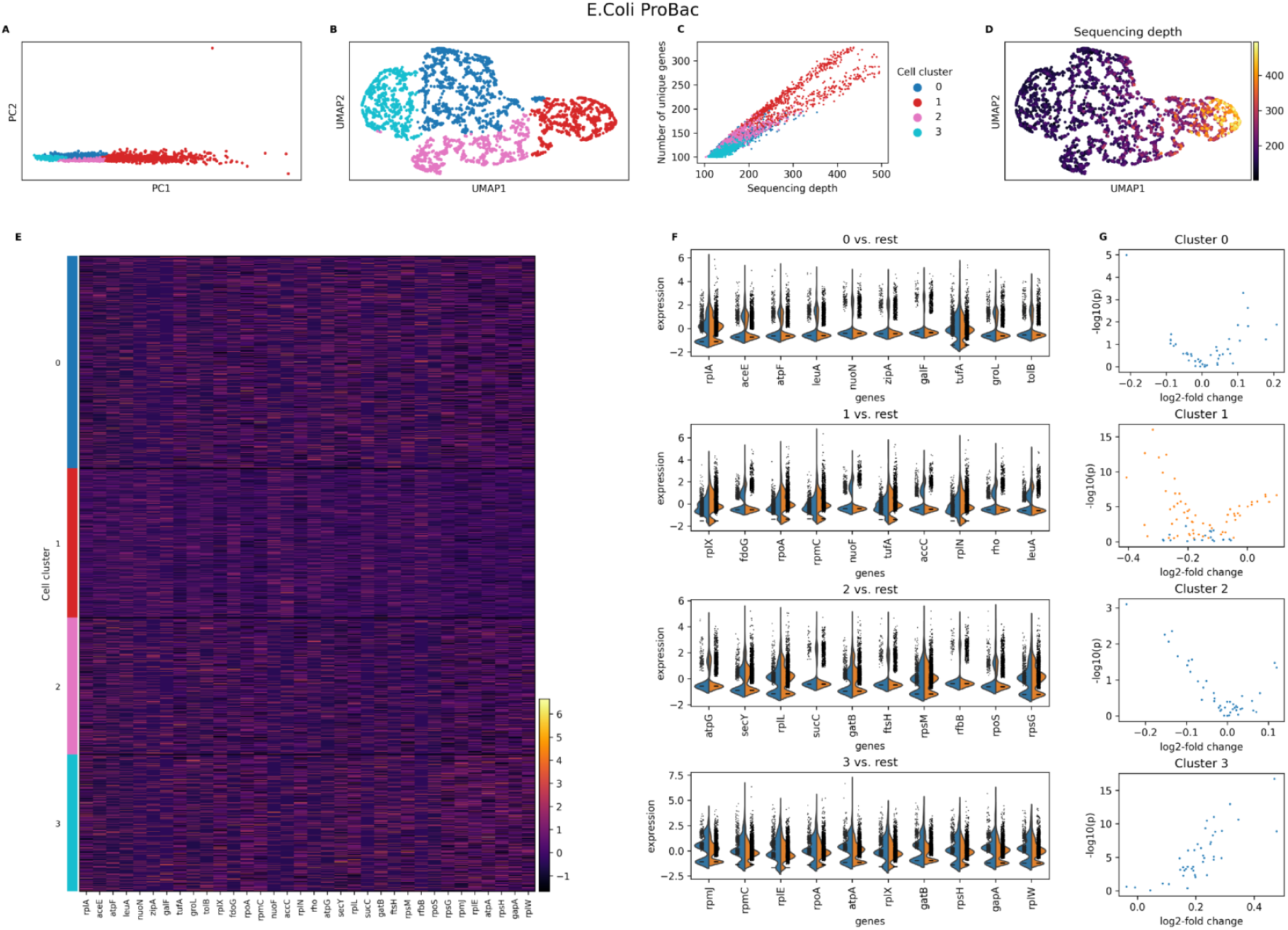
Analysis of the *Ecoli_balanced_PB* dataset with BacSC. **(A)** Scatterplot of first two dimensions of PCA embedding with cell type clusters highlighted. **(B)** UMAP plot based on the parameters determined by BacSC, colored by cell cluster. **(C)** Scatterplot of sequencing depth versus number of unique genes per cell, colored by cell cluster. **(D)** Umap plot as in (B), colored by sequencing depth. **(E)** Heatmap of normalized gene expression for all cells and characteristic genes for cell types. For each cluster, the 10 genes with the highest contrast scores are shown. **(F)** Violin plots of normalized gene expression for DE tests of each cell type (blue) against the rest of the cell population (orange). For each cluster, the 10 genes with the highest contrast score are shown. **(G)** Volcano plots for DE tests as in (F). The x-axis shows the log-fold change for gene expression, the y-axis shows the - log_10_-transformed (uncorrected) p-value. Only genes that are expressed in at least 80% of cells in the respective cluster are shown. Orange genes are differentially expressed at the a= 0.05-level.

**Fig. C10.**
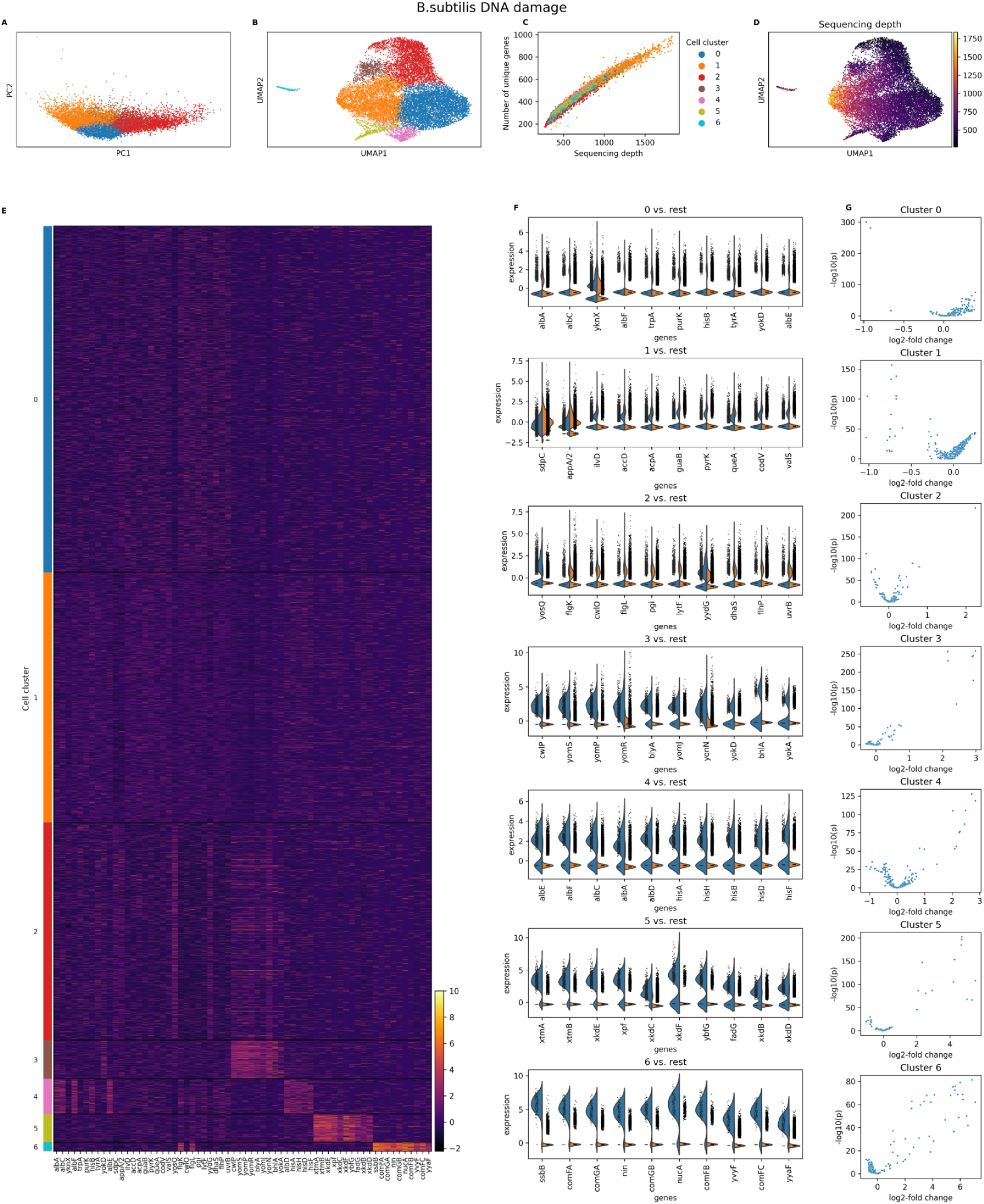
Analysis of the *Bsub_damage_PB* dataset with BacSC. **(A)** Scatterplot of first two dimensions of PCA embedding with cell type clusters highlighted. **(B)** UMAP plot based on the parameters determined by BacSC, colored by cell cluster. **(C)** Scatterplot of sequencing depth versus number of unique genes per cell, colored by cell cluster. **(D)** Umap plot as in (B), colored by sequencing depth. **(E)** Heatmap of normalized gene expression for all cells and characteristic genes for cell types. For each cluster, the 10 genes with the highest contrast scores are shown. For each cluster, the 10 genes with the highest contrast scores are shown. **(F)** Violin plots of normalized gene expression for DE tests of each cell type (blue) against the rest of the cell population (orange). For each cluster, the 10 genes with the highest contrast score are shown. **(G)** Volcano plots for DE tests as in **(F).** The x-axis shows the log-fold change for gene expression, the y-axis shows the - log_10_-transformed (uncorrected) p-value. Only genes that are expressed in at least 80% of cells in the respective cluster are shown. Orange genes are differentially expressed at the *a* = 0.05-leveL

**Fig. C11.**
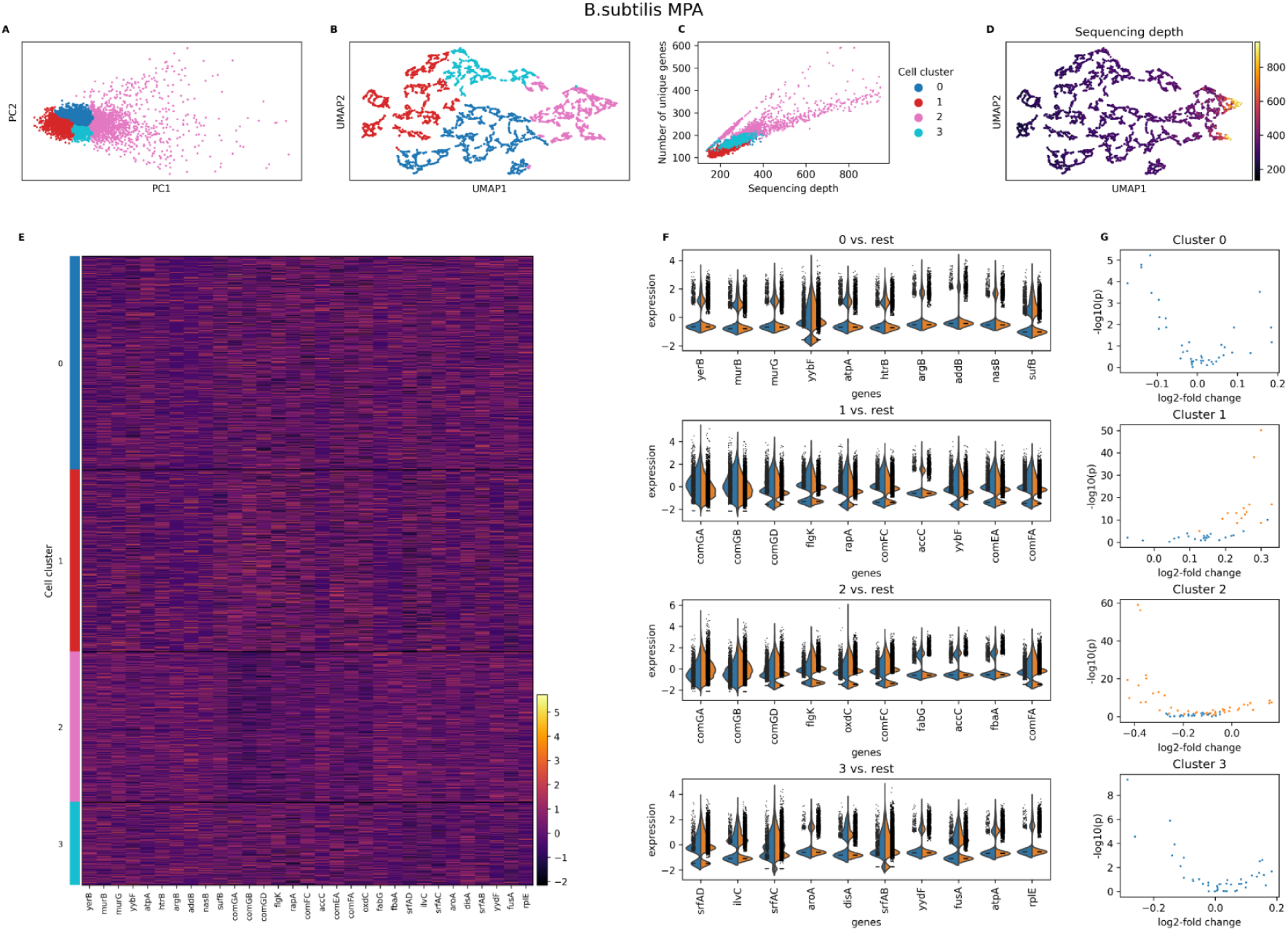
Analysis of the *Bsub_MPA_PB* dataset with BacSC. **(A)** Scatterplot of first two dimensions of PCA embedding with cell type clusters highlighted. **(B)** UMAP plot based on the parameters determined by BacSC, colored by cell cluster. **(C)** Scatterplot of sequencing depth versus number of unique genes per cell, colored by cell cluster. **(D)** Umap plot as in (B), colored by sequencing depth. **(E)** Heatmap of normalized gene expression for all cells and characteristic genes for cell types. For each cluster, the 10 genes with the highest contrast scores are shown. **(F)** Violin plots of normalized gene expression for DE tests of each cell type (blue) against the rest of the cell population (orange). For each cluster, the 10 genes with the highest contrast score are shown. **(G)** Volcano plots for DE tests as in (F). The x-axis shows the log-fold change for gene expression, the y-axis shows the - log_10_-transformed (uncorrected) p-value. Only genes that are expressed **in** at least 80% of cells in the respective cluster are shown. Orange genes are differentially expressed at the a= 0.05-level.

**Fig. C12.**
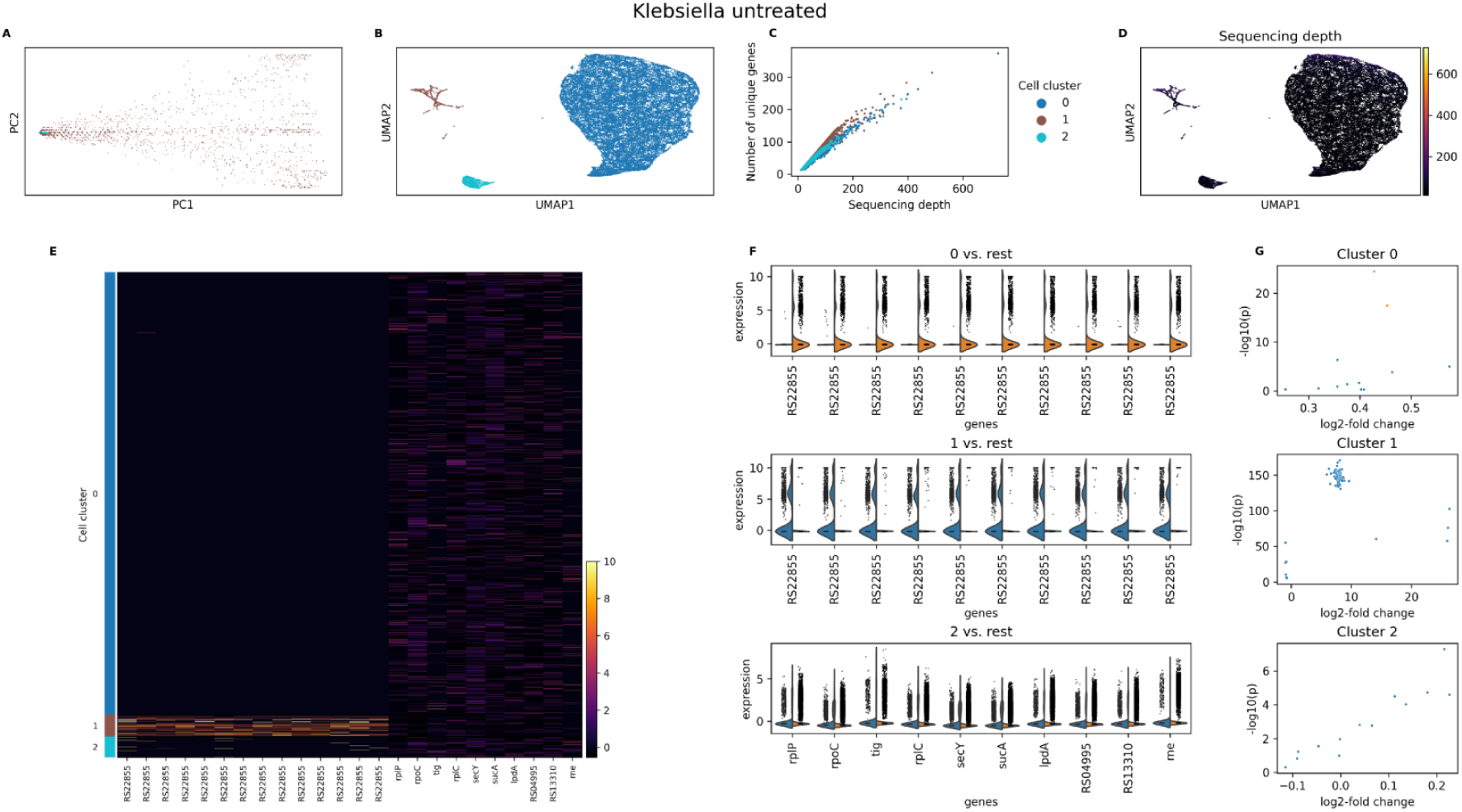
Analysis of the *Klebs_untreated_BD* dataset with BacSC. **(A)** Scatterplot of first two dimensions of PCA embedding with cell type clusters highlighted. **(B)** UMAP plot based on the parameters determined by BacSC, colored by cell cluster. **(C)** Scatterplot of sequencing depth versus number of unique genes per cell, colored by cell cluster. **(D)** Umap plot as **in** (B), colored by sequencing depth. **(E)** Heatmap of normalized gene expression for all cells and characteristic genes for cell types. For each cluster, the 10 genes with the highest contrast scores are shown. **(F)** Violin plots of normalized gene expression for DE tests of each cell type (blue) against the rest of the cell population (orange). For each cluster, the 10 genes with the highest contrast score are shown. Genes in (E) and (F) are annotated with gene symbols wherever possible, otherwise locus tags are shown. (G) Volcano plots for DE tests as in (F). The x-axis shows the log-fold change for gene expression, the y-axis shows the - log_10_-transformed (uncorrected) p-value. Only genes that are expressed in at least 50% of cells in the respective cluster are shown. Orange genes are differentially expressed at the *a* = 0.05-level.

**Fig. C13.**
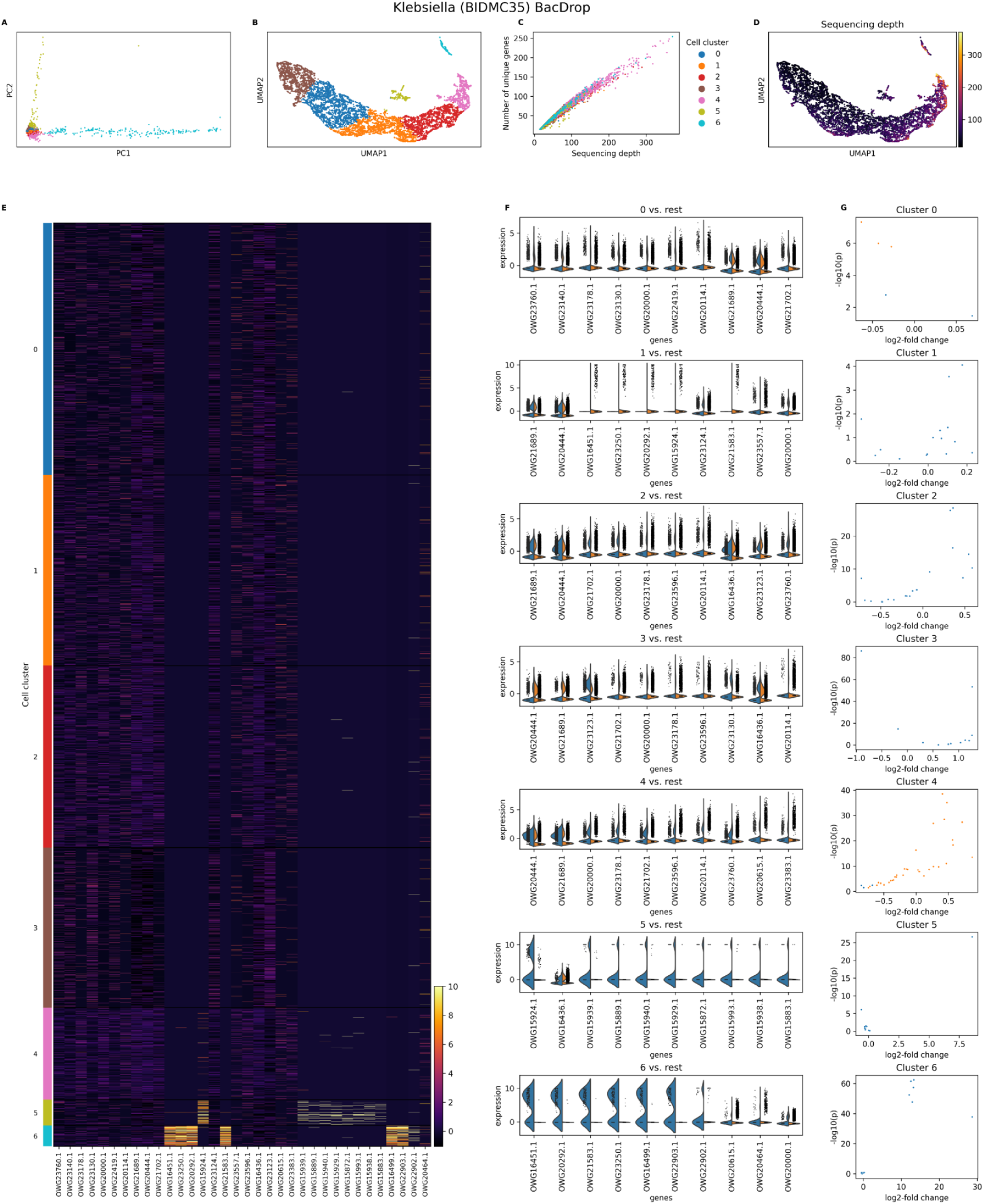
Analysis of the *Klebs_BJDMC35_BD* dataset with BacSC. **(A)** Scatterplot of first two dimensions of PCA embedding with cell type clusters highlighted. **(B)** UMAP plot based on the parameters determined by BacSC, colored by cell cluster. **(C)** Scatterplot of sequencing depth versus number of unique genes per cell, colored by cell cluster. **(D)** Umap plot as in (B), colored by sequencing depth. **(E)** Heatmap of normalized gene expression for all cells and characteristic genes for cell types. For each cluster, the 10 genes with the highest contrast scores are shown. **(F)** Violin plots of normalized gene expression for DE tests of each cell type (blue) against the rest of the cell population (orange). For each cluster, the 10 genes with the highest contrast score are shown. Genes in (E) and (F) are annotated with gene symbols wherever possible, otherwise locus tags are shown. **(G)** Volcano plots for DE tests as in (F). The x-axis shows the log-fold change for gene expression, the y-axis shows the - log_10_-transformed (uncorrected) p-value. Only genes that are expressed in at least 50% of cells in the respective cluster are shown. Orange genes are differentially expressed at the *a* = 0.05-leveL

**Fig. C14.**
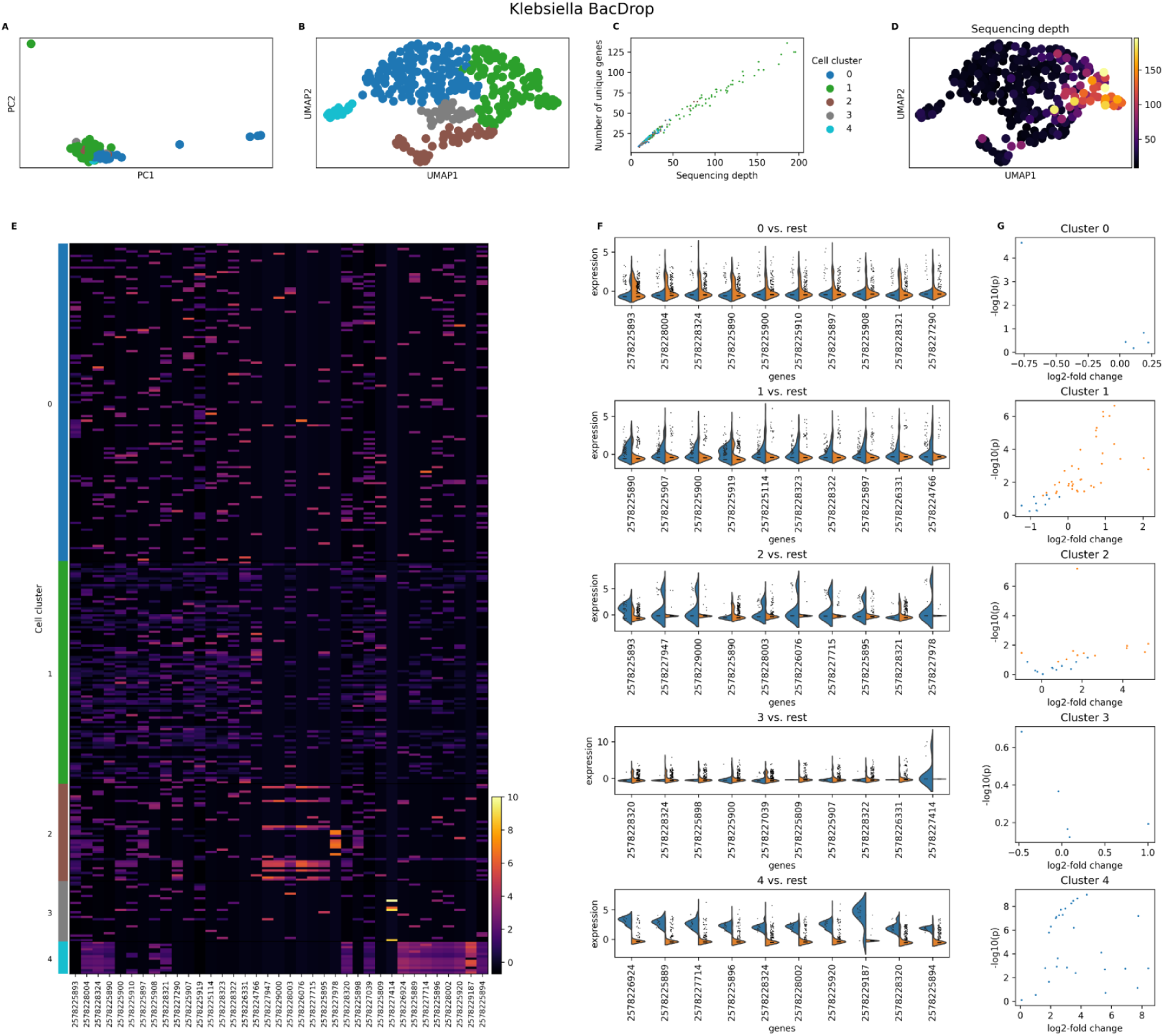
Analysis of the *Klebs_J,species_BD* dataset with BacSC. **(A)** Scatterplot of first two dimensions of PCA embedding with cell type clusters highlighted. **(B)** UMAP plot based on the parameters determined by BacSC, colored by cell cluster. **(C)** Scatterplot of sequencing depth versus number of unique genes per cell, colored by cell cluster. **(D)** Umap plot as in (B), colored by sequencing depth. **(E)** Heatmap of normalized gene expression for all cells and characteristic genes for cell types. For each cluster, the 10 genes with the highest contrast scores are shown. **(F)** Violin plots of normalized gene expression for DE tests of each cell type (blue) against the rest of the cell population (orange). For each cluster, the 10 genes with the highest contrast score are shown. Genes in (E) and (F) are annotated with gene symbols wherever possible, otherwise locus tags are shown. (G) Volcano plots for DE tests as in (F). The x-axis shows the log-fold change for gene expression, the y-axis shows the - log_10_-transformed (uncorrected) p-value. Only genes that are expressed in at least 50% of cells in the respective cluster are shown. Orange genes are differentially expressed at the a = 0.05-level.

**Fig. C15.**
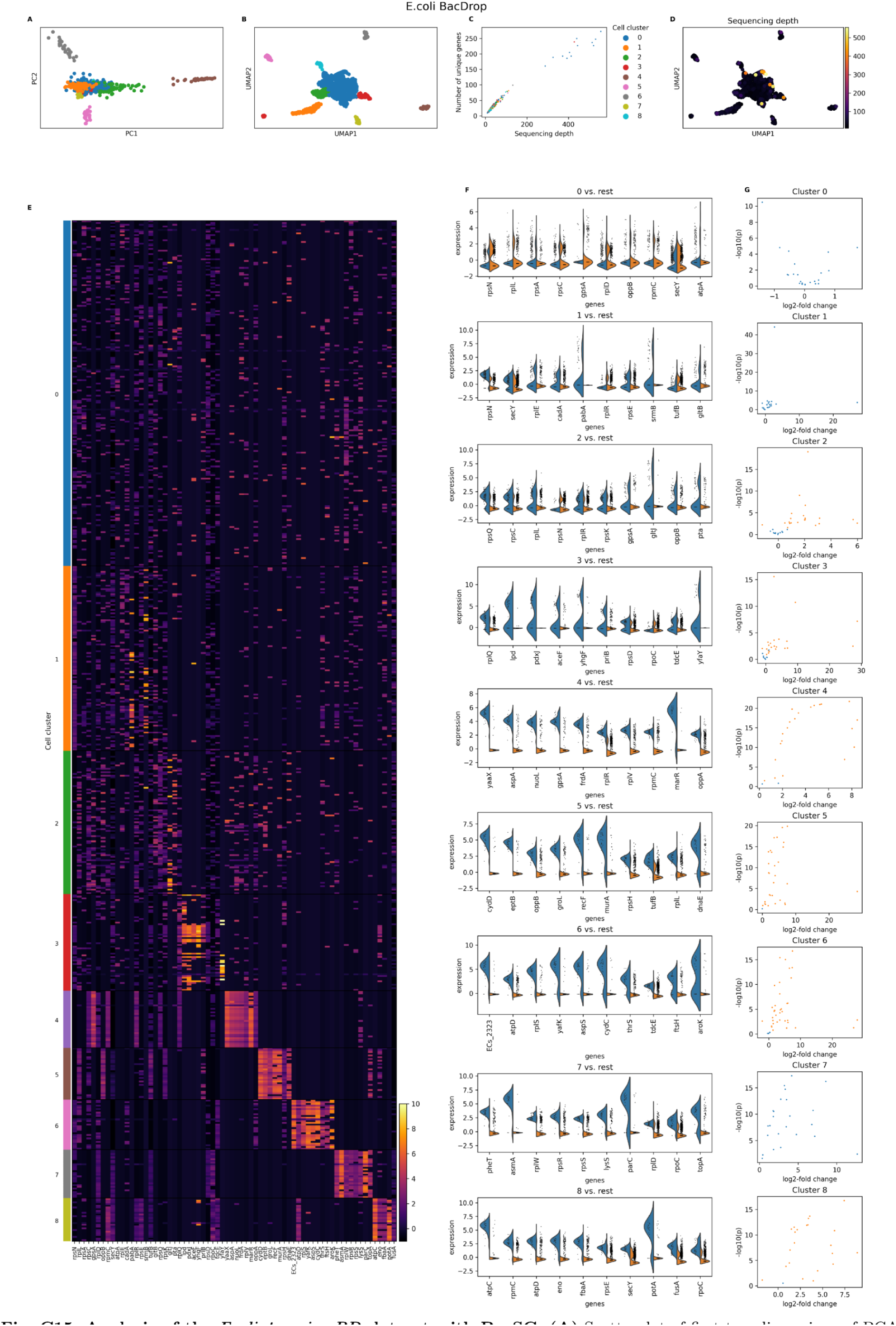
Analysis of the *Ecoli_4species_BD* dataset with BacSC. **(A)** Scatterplot of first two dimensions of PCA embedding with cell type clusters highlighted. **(B)** UMAP plot based on the parameters determined by BacSC, colored by cell cluster. **(C)** Scatterplot of sequencing depth versus number of unique genes per cell, colored by cell cluster. **(D)** Umap plot as in (B), colored by sequencing depth. **(E)** Heatmap of normalized gene expression for all cells and characteristic genes for cell types. For each cluster, the 10 genes with the highest contrast scores are shown. **(F)** Violin plots of normalized gene expression for DE tests of each cell type (blue) against the rest of the cell population (orange). For each cluster, the 10 genes with the highest contrast score are shown. Genes in (Elfland (F) are annotated with gene symbols wherever possible, otherwise locus tags are shown. (G) Volcano plots for DE tests as in (F). The x-axis shows the log-fold change for gene expression, the y-axis shows the - log_10_-transformed (uncorrected) p-value. Only genes that are expressed in at least 50% of cells in the respective cluster are shown. Orange genes are differentially expressed at the *a=* 0.05-level.

**Fig. C16.**
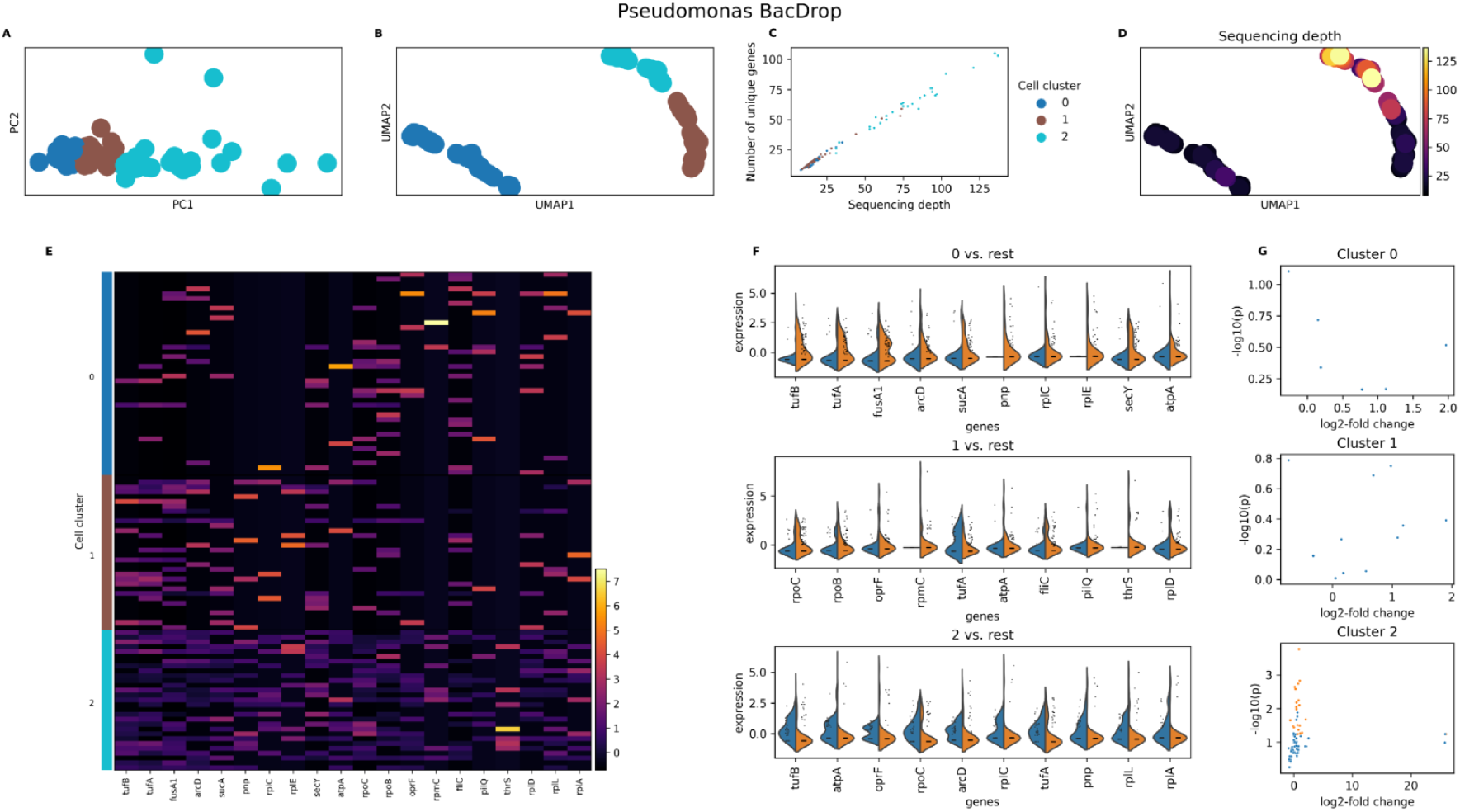
Analysis of the *Pseudomonas_4species_BD* dataset with BacSC. **(A)** Scatterplot of first two dimensions of PCA embedding with cell type clusters highlighted. **(B)** UMAP plot based on the parameters determined by BacSC, colored by cell cluster. (C) Scatterplot of sequencing depth versus number of unique genes per cell, colored by cell cluster. **(D)** Umap plot as in (B), colored by sequencing depth. (E) Heatmap of normalized gene expression for all cells and characteristic genes for cell types. For each cluster, the 10 genes with the highest contrast scores are shown. **(F)** Violin plots of normalized gene expression for DE tests of each cell type (blue) against the rest of the cell population (orange). For each cluster, the 10 genes with the highest contrast score are shown. Genes in (E) and (F) are annotated with gene symbols wherever possible, otherwise locus tags are shown. **(G)** Volcano plots for DE tests as in (F). The x-axis shows the log-fold change for gene expression, the y-axis shows the - log_10_-transformed (uncorrected) p-value. Only genes that are expressed in at least 50% of cells in the respective cluster are shown. Orange genes are differentially expressed at the *a* = 0.05-level.

**Fig. C17.**
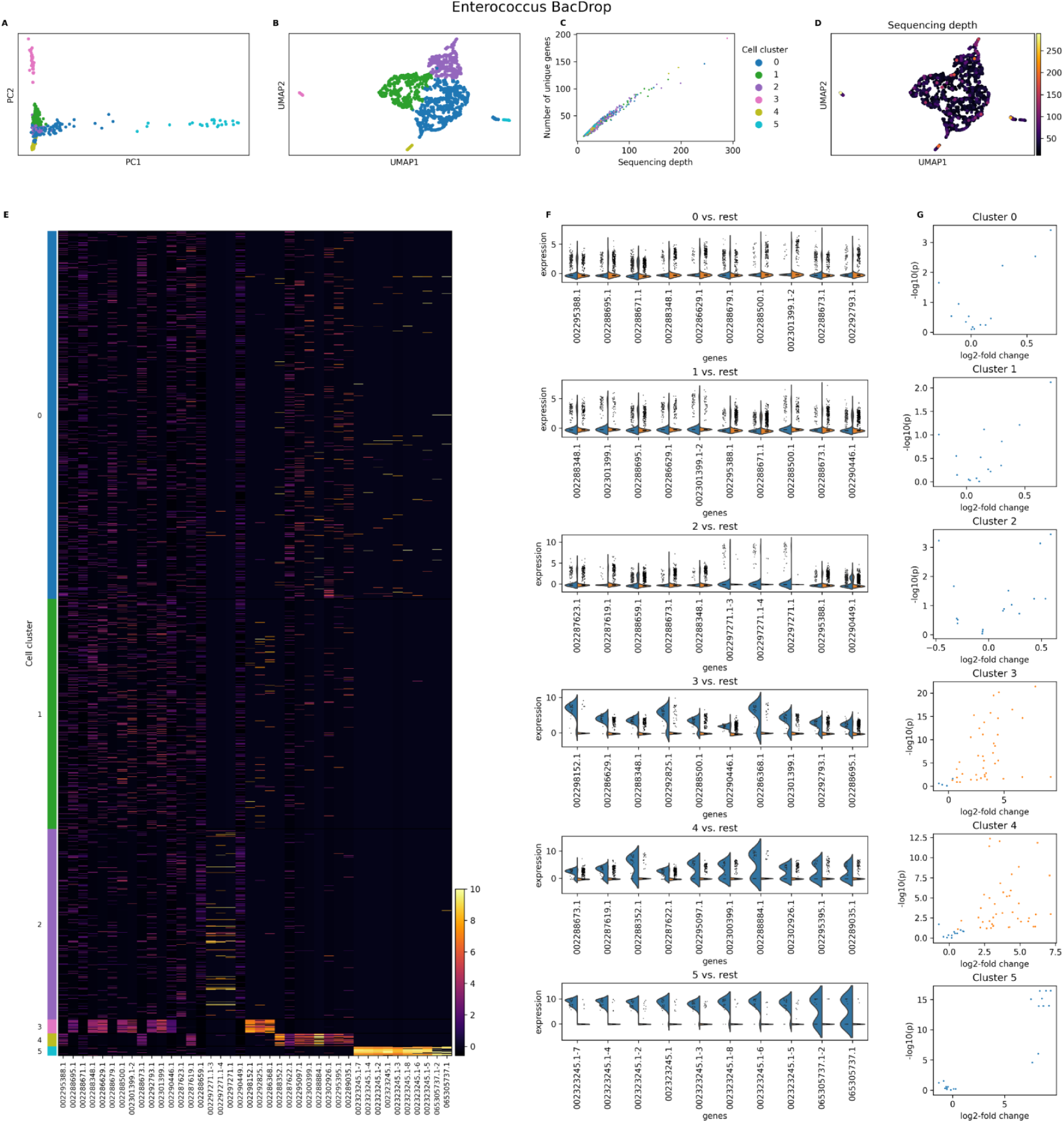
Analysis of the *Efaecium_4species_BD* dataset with BacSC. **(A)** Scatterplot of first two dimensions of PCA embedding with cell type clusters highlighted. **(B)** UMAP plot based on the parameters determined by BacSC, colored by cell cluster. (C) Scatterplot of sequencing depth versus number of unique genes per cell, colored by cell cluster. **(D)** Umap plot as in (B), colored by sequencing depth. **(E)** Heatmap of normalized gene expression for all cells and characteristic genes for cell types. For each cluster, the 10 genes with the highest contrast scores are shown. **(F)** Violin plots of normalized gene expression for DE tests of each cell type (blue) against the rest of the cell population (orange). For each cluster, the 10 genes with the highest contrast score are shown. Genes in (E) and (F) are annotated with gene symbols wherever possible, otherwise locus tags are shown. (G) Volcano plots for DE tests as in (F). The x-axis shows the log-fold change for gene expression, the y-axis shows the - log_10_-transformed (uncorrected) p-value. Only genes that are expressed in at least 50% of cells in the respective cluster are shown. Orange genes are differentially expressed at the *a* = 0.05-leveL

## Appendix D Diagnostic plots for all datasets

This section contains a selection of diagnostic plots from BacSC for each dataset from table 1.

**Fig. D18.**
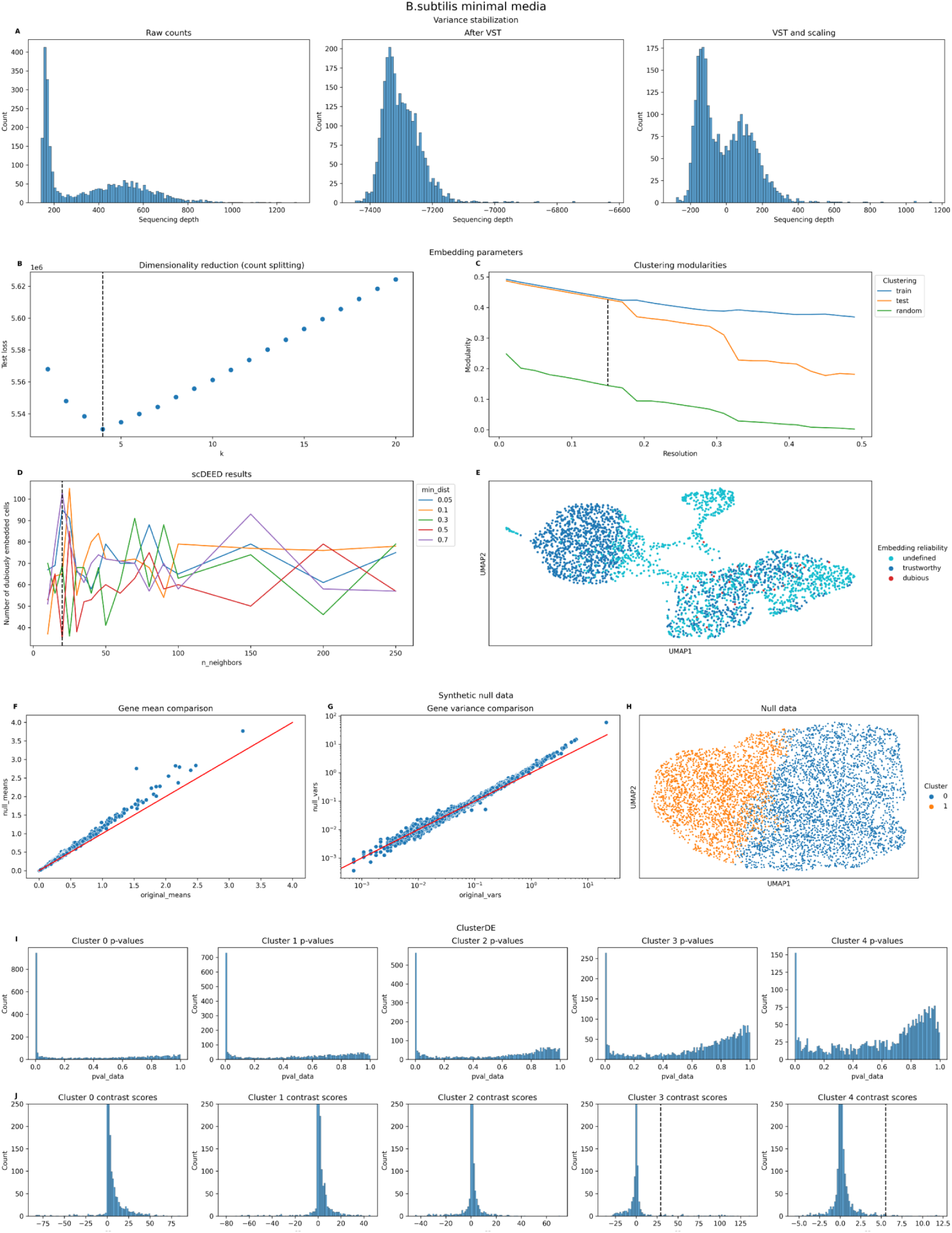
Diagnostic plots generated during the analysis of the *Bsub_minmed_PB* dataset with BacSC. **(A)** Histograms of cell sequencing depth for raw gene expression data (left), after variance-stabilizing transform (middle), and after gene-wise scaling (right). **(B)** Selection of latent dimensionality through count splitting. The dashed line indicates *kopt* selected by BacSC. **(C)** Selection of clustering resolution. The dashed line indicates the maximal gap in modularity between the training data Leiden clustering applied to the test data and a random clustering on the test data. **(D)** Selection of *nneighbors* and *mindist* parameters for UMAP embedding with scDEED. The dashed black line shows the value of *nneighbors* selected by BacSC. **(E)** UMAP embedding generated with BacSC-selected parameters, colored by embedding reliability determined by scDEED with the same parameters. **(F)** Comparison of gene means of original and synthetic null data for DE testing. **(G)** Comparison of gene variances of original and synthetic null data for DE testing. **(H)** UMAP of synthetic null data for DE testing of each cell type against all other cells, colored by the two-group clustering determined for calculation of contrast scores. (I) Histograms of uncorrected p-values for DE testing of each cell type against all other cells. **(J)** Histograms of contrast scores for DE testing of each cell type against all other cells. The dashed lines indicate significance threshold values at the FDR level *a* = 0.05.

**Fig. D19.**
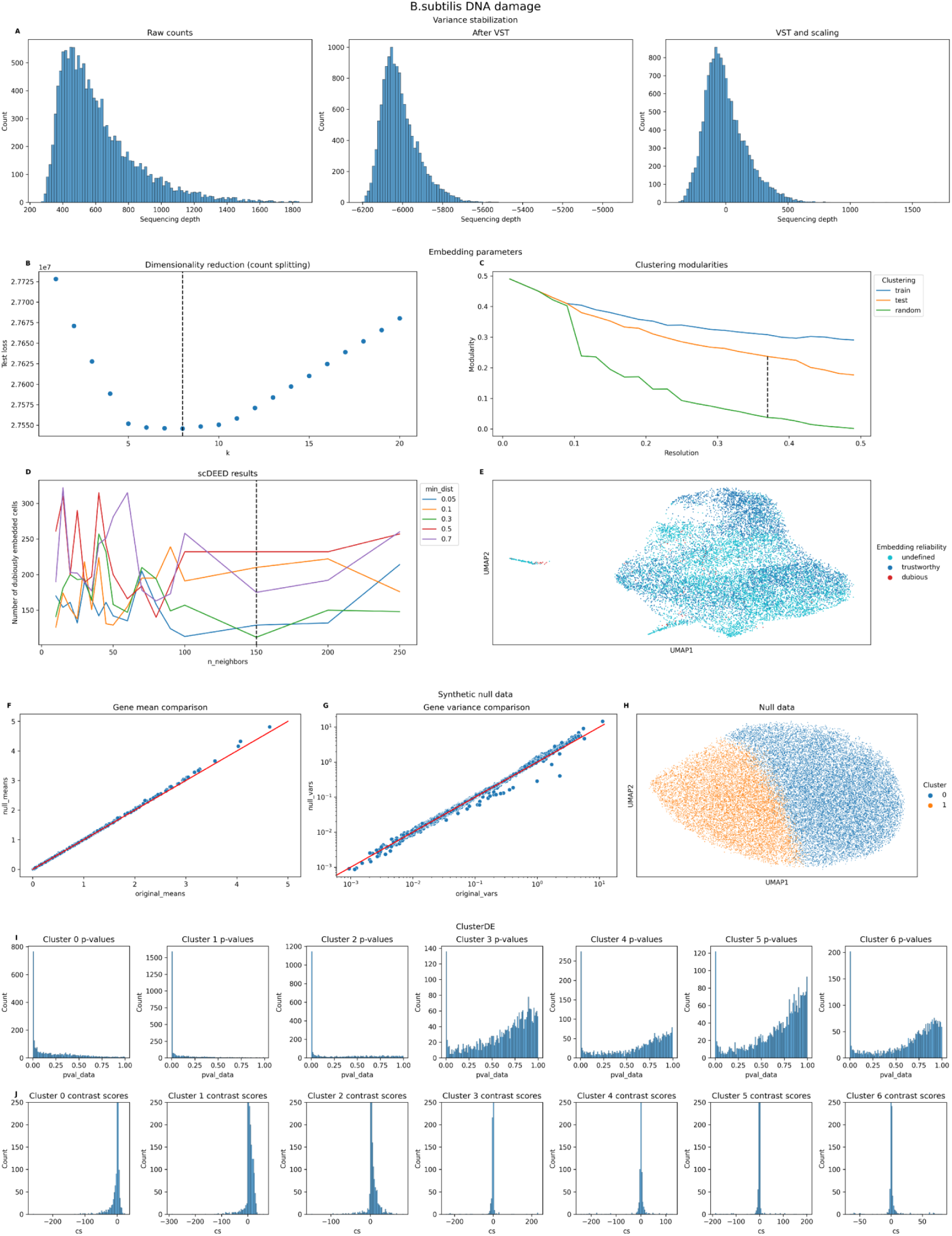
Diagnostic plots generated during the analysis of the *Bsub_damage_PB* dataset with BacSC. **(A)** Histograms of cell sequencing depth for raw gene expression data (left), after variance-stabilizing transform (middle), and after gene-wise scaling (right). **(B)** Selection of latent dimensionality through count splitting. The dashed line indicates *kopt* selected by BacSC. **(C)** Selection of clustering resolution. The dashed line indicates the maximal gap in modularity between the training data Leiden clustering applied to the test data and a random clustering on the test data. **(D)** Selection of *nneighbors* and *mindist* parameters for UMAP embedding with scDEED. The dashed black line shows the value of *nneighbors* selected by BacSC. **(E)** UMAP embedding generated with BacSC-selected parameters, colored by embedding reliability determined by scDEED with the same parameters. **(F)** Comparison of gene means of original and synthetic null data for DE testing. **(G)** Comparison of gene variances of original and synthetic null data for DE testing. **(H)** UMAP of synthetic null data for DE testing of each cell type against all other cells, colored by the two-group clustering determined for calculation of contrast scores. (I) Histograms of uncorrected p-values for DE testing of each cell type against all other cells. **(J)** Histograms of contrast scores for DE testing of each cell type against all other cells. The dashed lines indicate significance threshold values at the FDR level *a* = 0.05.

**Fig. D20.**
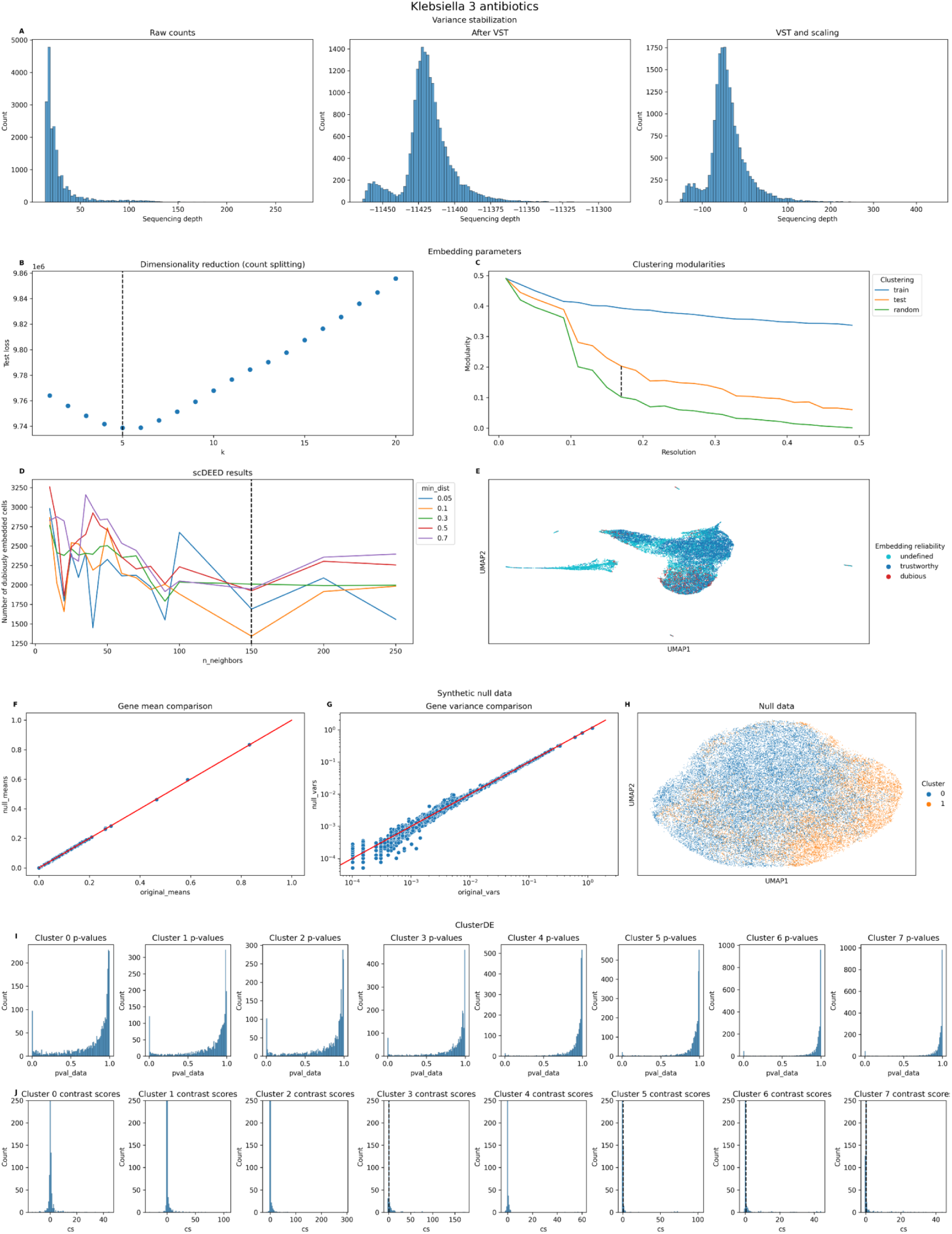
Diagnostic plots generated during the analysis of the *Klebs_antibiotics_BD* dataset with BacSC. **(A)** Histograms of cell sequencing depth for raw gene expression data (left), after variance-stabilizing transform (middle), and after gene-wise scaling (right). **(B)** Selection of latent dimensionality through count splitting. The dashed line indicates *kopt* selected by BacSC. **(C)** Selection of clustering resolution. The dashed line indicates the maximal gap in modularity between the training data Leiden clustering applied to the test data and a random clustering on the test data. **(D)** Selection of *nneighbors* and *mindist* parameters for UMAP embedding with scDEED. The dashed black line shows the value of *nneighbors* selected by BacSC. **(E)** UMAP embedding generated with BacSC-selected parameters, colored by embedding reliability determined by scDEED with the same parameters. **(F)** Comparison of gene means of original and synthetic null data for DE testing. **(G)** Comparison of gene variances of original and synthetic null data for DE testing. **(H)** UMAP of synthetic null data for DE testing of each cell type against all other cells, colored by the two-group clustering determined for calculation of contrast scores. (I) Histograms of uncorrected p-values for DE testing of each cell type against all other cells. **(J)** Histograms of contrast scores for DE testing of each cell type against all other cells. The dashed lines indicate significance threshold values at the FDR level *a* = 0.05.

**Fig. D21.**
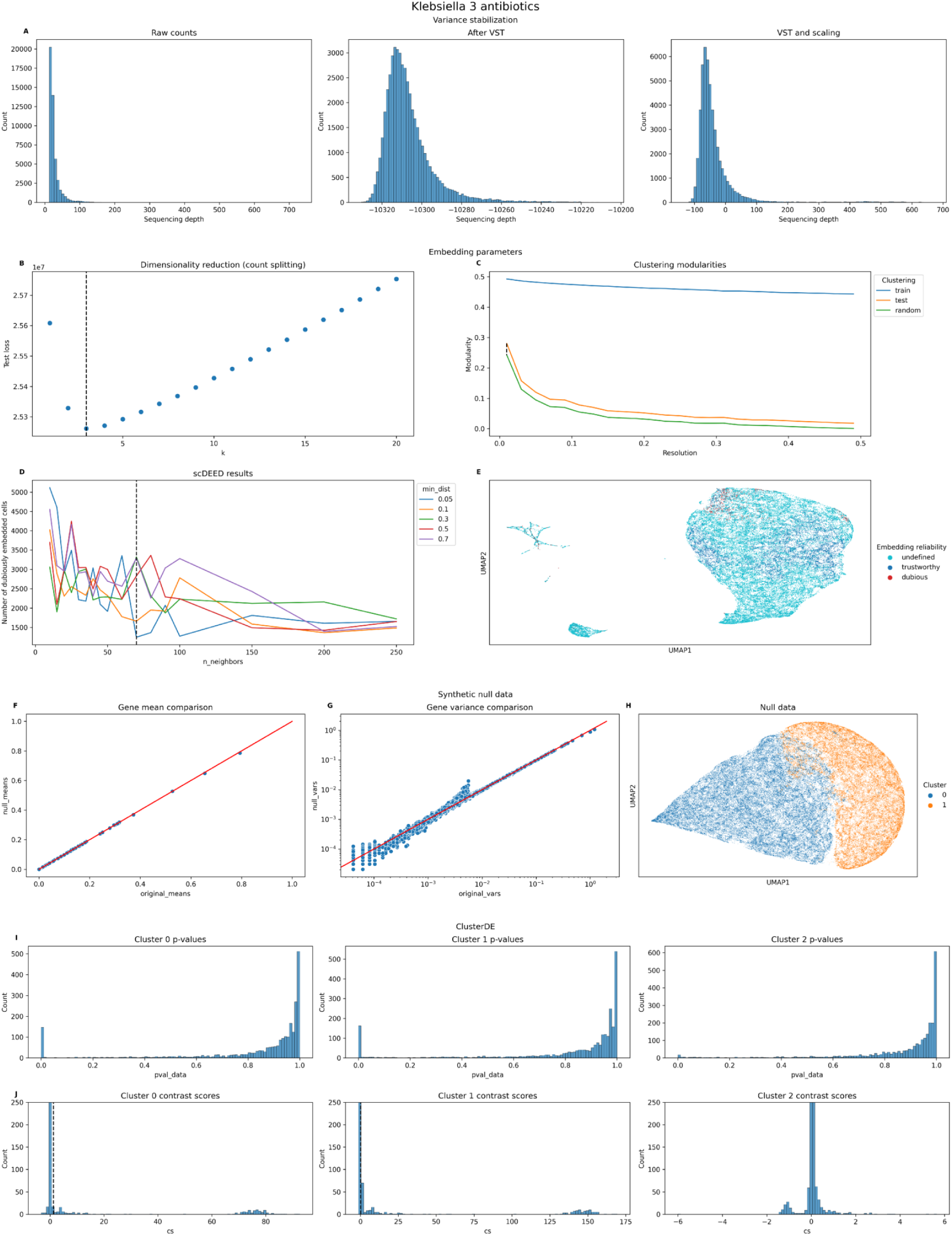
Diagnostic plots generated during the analysis of the *Klebs_untreated_BD* dataset with BacSC. **(A)** Histograms of cell sequencing depth for raw gene expression data (left), after variance-stabilizing transform (middle), and after gene-wise scaling (right). **(B)** Selection of latent dimensionality through count splitting. The dashed line indicates *kopt* selected by BacSC. **(C)** Selection of clustering resolution. The dashed line indicates the maximal gap in modularity between the training data Leiden clustering applied to the test data and a random clustering on the test data. **(D)** Selection of *nneighbors* and *mindist* parameters for UMAP embedding with scDEED. The dashed black line shows the value of *nneighbors* selected by BacSC. **(E)** UMAP embedding generated with BacSC-selected parameters, colored by embedding reliability determined by scDEED with the same parameters. **(F)** Comparison of gene means of original and synthetic null data for DE testing. **(G)** Comparison of gene variances of original and synthetic null data for DE testing. **(H)** UMAP of synthetic null data for DE testing of each cell type against all other cells, colored by the two-group clustering determined for calculation of contrast scores. (I) Histograms of uncorrected p-values for DE testing of each cell type against all other cells. **(J)** Histograms of contrast scores for DE testing of each cell type against all other cells. The dashed lines indicate significance threshold values at the FDR level *a* = 0.05.

**Fig. D22.**
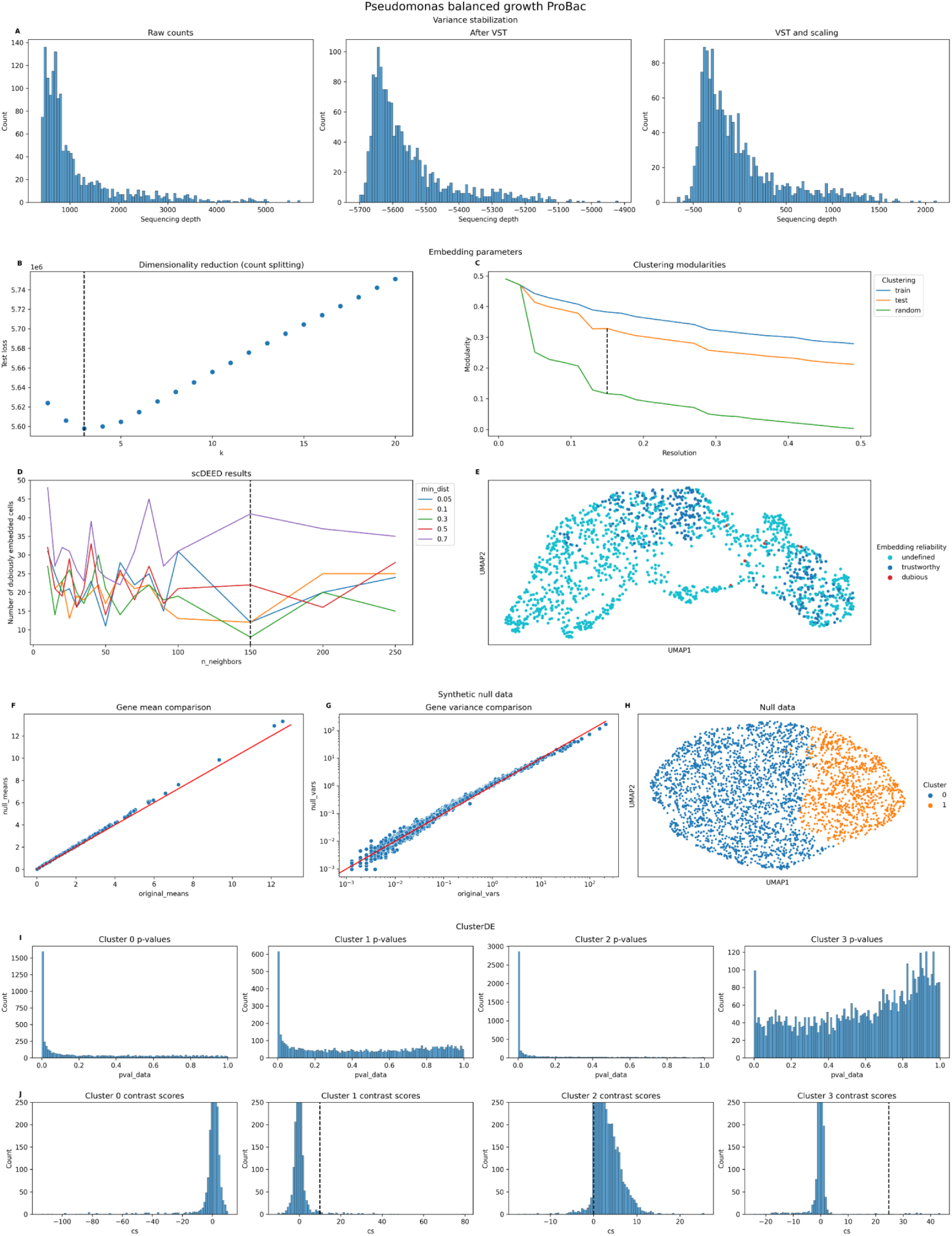
Diagnostic plots generated during the analysis of the *Pseudomonas_balanced_PB* dataset with BacSC. **(A)** Histograms of cell sequencing depth for raw gene expression data (left), after variance-stabilizing transform (middle), and after gene-wise scaling (right). **(B)** Selection of latent dimensionality through count splitting. The dashed line indicates *kopt* selected by BacSC. **(C)** Selection of clustering resolution. The dashed line indicates the maximal gap in modularity between the training data Leiden clustering applied to the test data and a random clustering on the test data. **(D)** Selection of *nneighbors* and *mindist* parameters for UMAP embedding with scDEED. The dashed black line shows the value of *nneighbors* selected by BacSC. **(E)** UMAP embedding generated with BacSC-selected parameters, colored by embedding reliability determined by scDEED with the same parameters. **(F)** Comparison of gene means of original and synthetic null data for DE testing. **(G)** Comparison of gene variances of original and synthetic null data for DE testing. **(H)** UMAP of synthetic null data for DE testing of each cell type against all other cells, colored by the two-group clustering determined for calculation of contrast scores. (I) Histograms of uncorrected p-values for DE testing of each cell type against all other cells. (J) Histograms of contrast scores for DE testing of each cell type against all other cells. The dashed lines indicate significance threshold values at the FDR level °’ = 0.05.

**Fig. D23.**
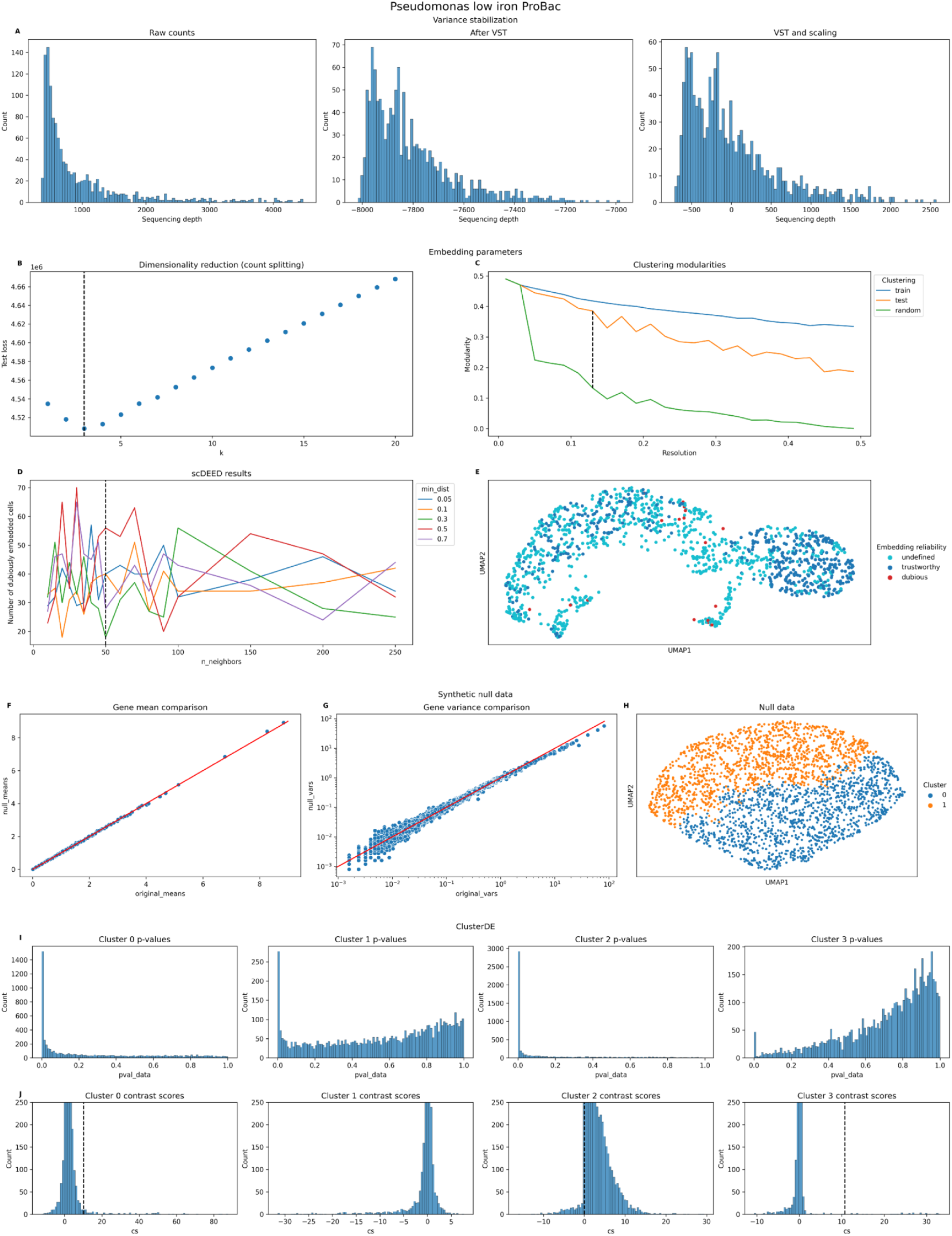
Diagnostic plots generated during the analysis of the *Pseudomonas_/i_PB* dataset with BacSC. **(A)** Histograms of cell sequencing depth for raw gene expression data (left), after variance-stabilizing transform (middle), and after gene-wise scaling (right). **(B)** Selection of latent dimensionality through count splitting. The dashed line indicates *kopt* selected by BacSC. **(C)** Selection of clustering resolution. The dashed line indicates the maximal gap in modularity between the training data Leiden clustering applied to the test data and a random clustering on the test data. **(D)** Selection of *nneighbors* and *mindist* parameters for UMAP embedding with scDEED. The dashed black line shows the value of *nneighbors* selected by BacSC. **(E)** UMAP embedding generated with BacSC-selected parameters, colored by embedding reliability determined by scDEED with the same parameters. **(F)** Comparison of gene means of original and synthetic null data for DE testing. **(G)** Comparison of gene variances of original and synthetic null data for DE testing. **(H)** UMAP of synthetic null data for DE testing of each cell type against all other cells, colored by the two-group clustering determined for calculation of contrast scores. (I) Histograms of uncorrected p-values for DE testing of each cell type against all other cells. **(J)** Histograms of contrast scores for DE testing of each cell type against all other cells. The dashed lines indicate significance threshold values at the FDR level *a* = 0.05.

**Fig. D24.**
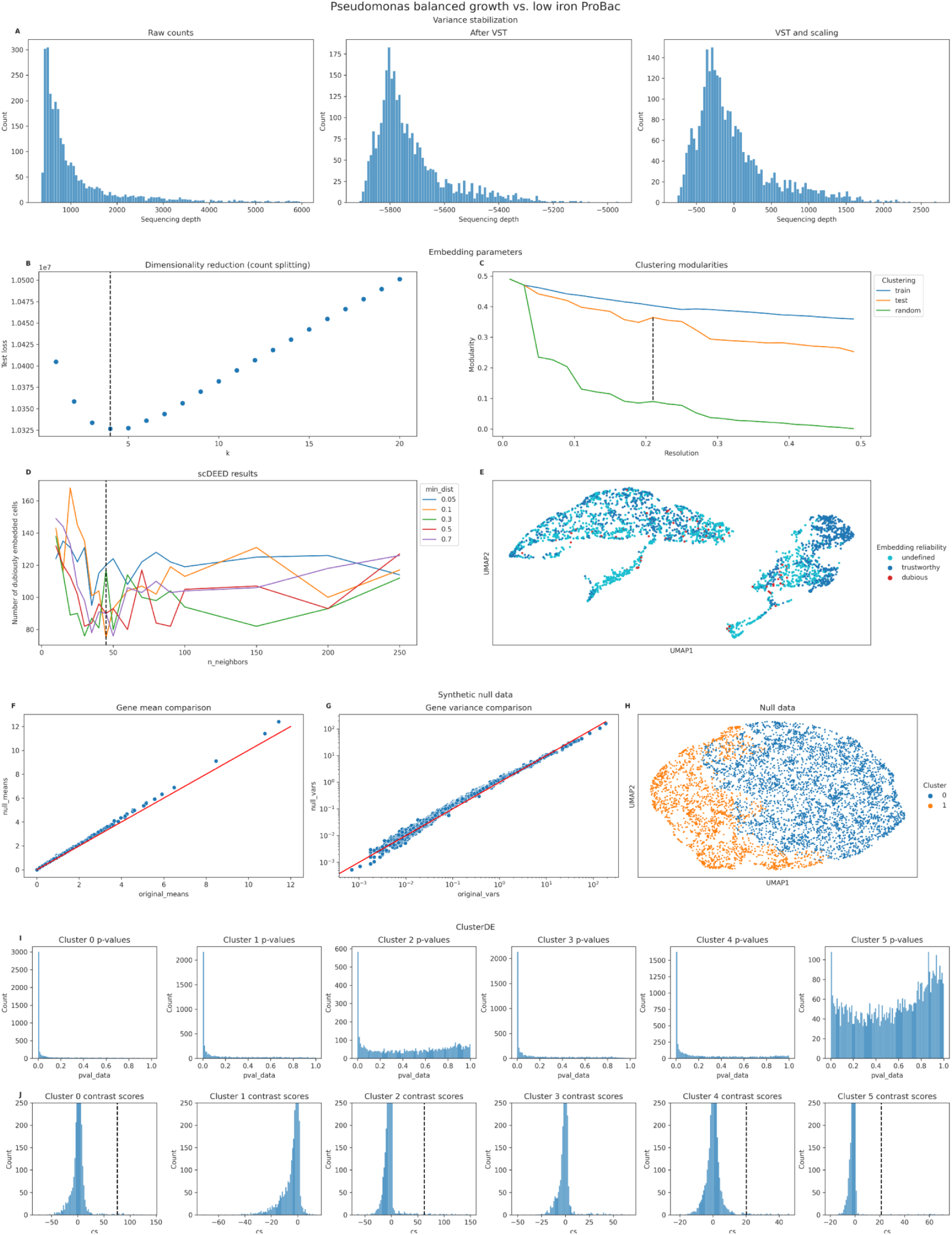
Diagnostic plots generated during the analysis of the combined *Pseudomonas_balanced_PB* and *Pseudomonas_li_PB* dataset with BacSC. **(A)** Histograms of cell sequencing depth for raw gene expression data (left), after variance-stabilizing transform (middle), and after gene-wise scaling (right). **(B)** Selection of latent dimensionality through count splitting. The dashed line indicates *kopt* selected by BacSC. **(C)** Selection of clustering resolution. The dashed line indicates the maximal gap in modularity between the training data Leiden clustering applied to the test data and a random clustering on the test data. **(D)** Selection of *nneighbors* and *mindist* parameters for UMAP embedding with scDEED. The dashed black line shows the value of *nneighbors* selected by BacSC. **(E)** UMAP embedding generated with BacSC-selected parameters, colored by embedding reliability determined by scDEED with the same parameters. **(F)** Comparison of gene means of original and synthetic null data for DE testing. **(G)** Comparison of gene variances of original and synthetic null data for DE testing. **(H)** UMAP of synthetic null data for DE testing of each cell type against all other cells, colored by the two-group clustering determined for calculation of contrast scores. (I) Histograms of uncorrected p­ values for DE testing of each cell type against all other cells. **(J)** Histograms of contrast scores for DE testing of each cell type against all other cells. The dashed lines indicate significance threshold values at the FDR level °’ = 0.05.

**Fig. D25.**
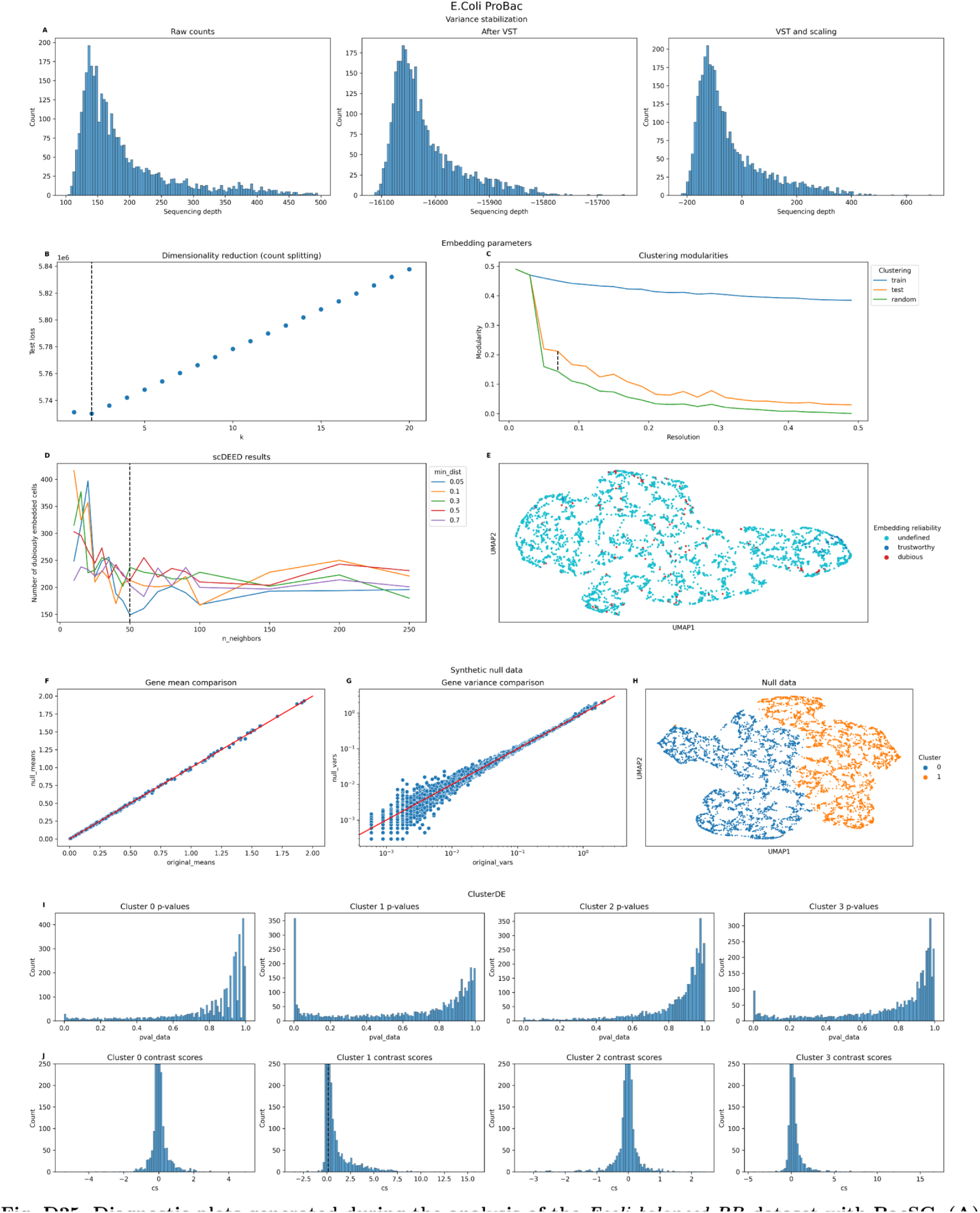
Diagnostic plots generated during the analysis of the *Ecoli_balanced_PB* dataset with BacSC. **(A)** Histograms of cell sequencing depth for raw gene expression data (left), after variance-stabilizing transform (middle), and after gene-wise scaling (right). **(B)** Selection of latent dimensionality through count splitting. The dashed line indicates *kopt* selected by BacSC. **(C)** Selection of clustering resolution. The dashed line indicates the maximal gap in modularity between the training data Leiden clustering applied to the test data and a random clustering on the test data. **(D)** Selection of *nneighbors* and *mindist* parameters for UMAP embedding with scDEED. The dashed black line shows the value of *nneighbors* selected by BacSC. **(E)** UMAP embedding generated with BacSC-selected parameters, colored by embedding reliability determined by scDEED with the same parameters. **(F)** Comparison of gene means of original and synthetic null data for DE testing. **(G)** Comparison of gene variances of original and synthetic null data for DE testing. **(H)** UMAP of synthetic null data for DE testing of each cell type against all other cells, colored by the two-group clustering determined for calculation of contrast scores. (I) Histograms of uncorrected p-values for DE testing of each cell type against all other cells. **(J)** Histograms of contrast scores for DE testing of each cell type against all other cells. The dashed lines indicate significance threshold values at the FDR level *a* = 0.05.

**Fig. D26.**
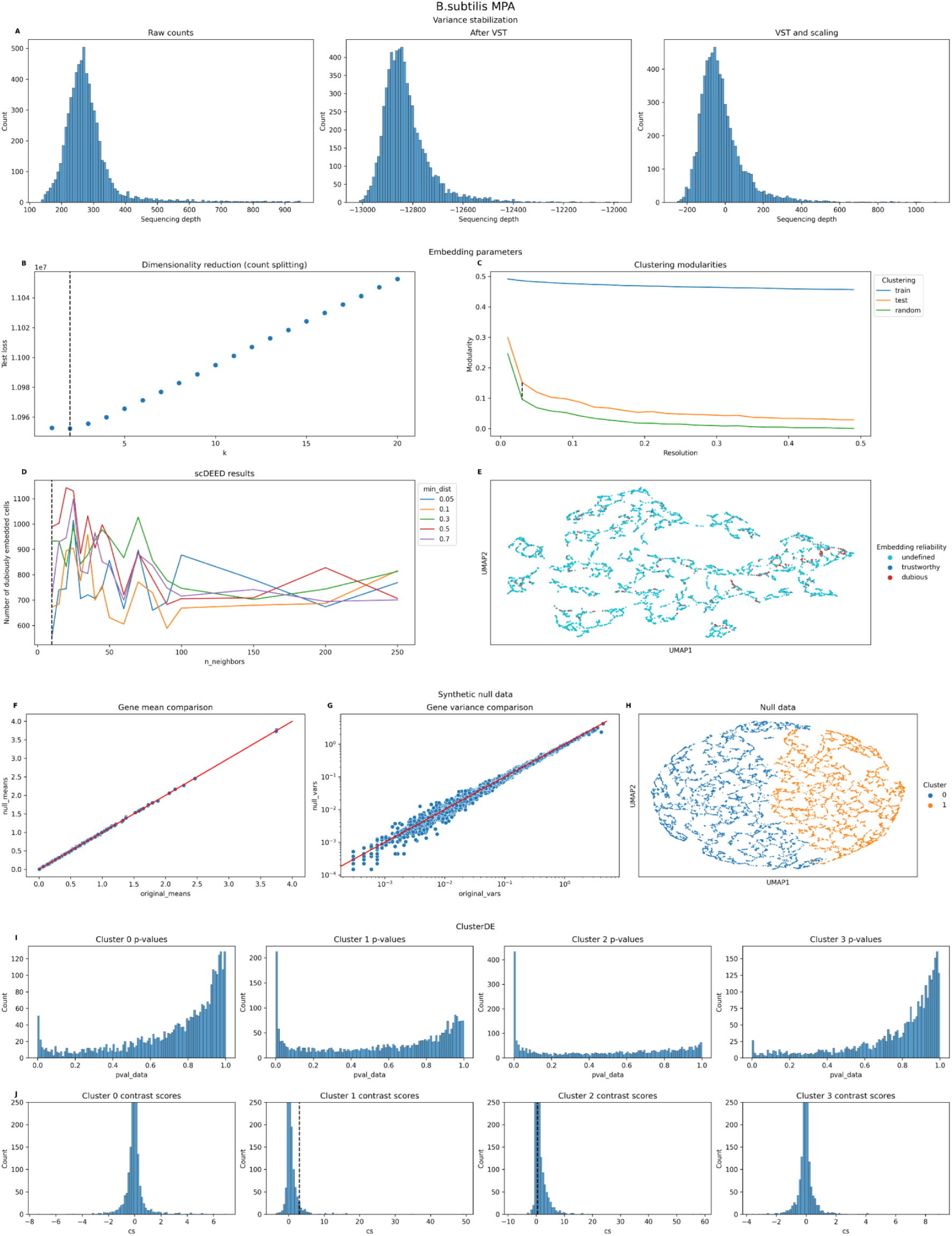
Diagnostic plots generated during the analysis of the *Bsub_MPA_PB* dataset with BacSC. **(A)** Histograms of cell sequencing depth for raw gene expression data (left), after variance-stabilizing transform (middle), and after gene-wise scaling (right). **(B)** Selection of latent dimensionality through count splitting. The dashed line indicates *kopt* selected by BacSC. **(C)** Selection of clustering resolution. The dashed line indicates the maximal gap in modularity between the training data Leiden clustering applied to the test data and a random clustering on the test data. **(D)** Selection of *nneighbors* and *mindist* parameters for UMAP embedding with scDEED. The dashed black line shows the value of *nneighbors* selected by BacSC. **(E)** UMAP embedding generated with BacSC-selected parameters, colored by embedding reliability determined by scDEED with the same parameters. **(F)** Comparison of gene means of original and synthetic null data for DE testing. **(G)** Comparison of gene variances of original and synthetic null data for DE testing. **(H)** UMAP of synthetic null data for DE testing of each cell type against all other cells, colored by the two-group clustering determined for calculation of contrast scores. (I) Histograms of uncorrected p-values for DE testing of each cell type against all other cells. **(J)** Histograms of contrast scores for DE testing of each cell type against all other cells. The dashed lines indicate significance threshold values at the FDR level *a* = 0.05. Klebsiella (BIDMC35) BacDrop

**Fig. D27.**
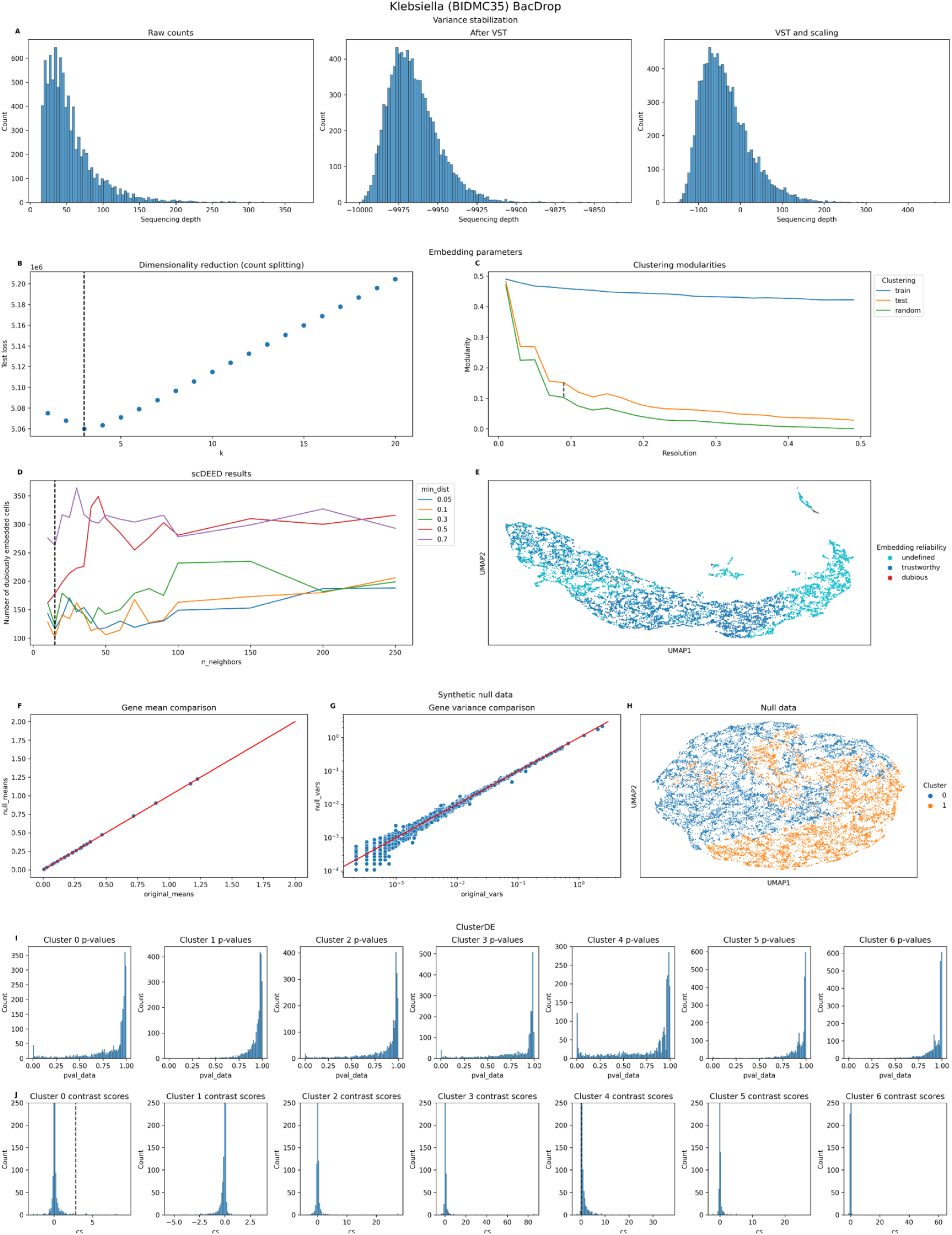
Diagnostic plots generated during the analysis of the *Klebs_BJDMC35_BD* dataset with BacSC. **(A)** Histograms of cell sequencing depth for raw gene expression data (left), after variance-stabilizing transform (middle), and after gene-wise scaling (right). **(B)** Selection of latent dimensionality through count splitting. The dashed line indicates *kopt* selected by BacSC. **(C)** Selection of clustering resolution. The dashed line indicates the maximal gap in modularity between the training data Leiden clustering applied to the test data and a random clustering on the test data. **(D)** Selection of *nneighbors* and *mindist* parameters for UMAP embedding with scDEED. The dashed black line shows the value of *nneighbors* selected by BacSC. **(E)** UMAP embedding generated with BacSC-selected parameters, colored by embedding reliability determined by scDEED with the same parameters. **(F)** Comparison of gene means of original and synthetic null data for DE testing. **(G)** Comparison of gene variances of original and synthetic null data for DE testing. **(H)** UMAP of synthetic null data for DE testing of each cell type against all other cells, colored by the two-group clustering determined for calculation of contrast scores. (I) Histograms of uncorrected p-values for DE testing of each cell type against all other cells. **(J)** Histograms of contrast scores for DE testing of each cell type against all other cells. The dashed lines indicate significance threshold values at the FDR level *a* = 0.05.

**Fig. D28.**
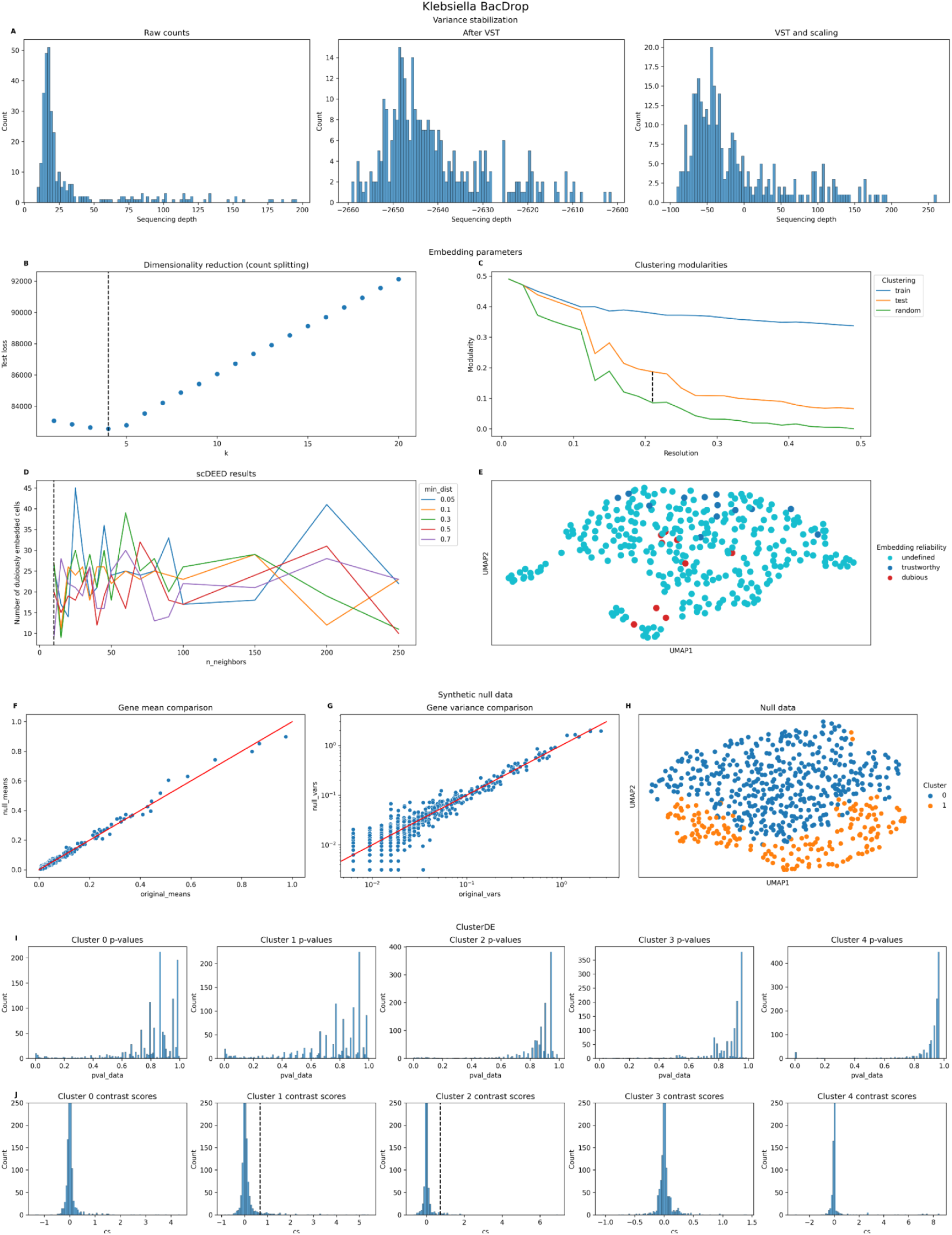
Diagnostic plots generated during the analysis of the *Klebs_J,species_BD* dataset with BacSC. **(A)** Histograms of cell sequencing depth for raw gene expression data (left), after variance-stabilizing transform (middle), and after gene-wise scaling (right). **(B)** Selection of latent dimensionality through count splitting. The dashed line indicates *kopt* selected by BacSC. **(C)** Selection of clustering resolution. The dashed line indicates the maximal gap in modularity between the training data Leiden clustering applied to the test data and a random clustering on the test data. **(D)** Selection of *nneighbors* and *mindist* parameters for UMAP embedding with scDEED. The dashed black line shows the value of *nneighbors* selected by BacSC. **(E)** UMAP embedding generated with BacSC-selected parameters, colored by embedding reliability determined by scDEED with the same parameters. **(F)** Comparison of gene means of original and synthetic null data for DE testing. **(G)** Comparison of gene variances of original and synthetic null data for DE testing. **(H)** UMAP of synthetic null data for DE testing of each cell type against all other cells, colored by the two-group clustering determined for calculation of contrast scores. (I) Histograms of uncorrected p-values for DE testing of each cell type against all other cells. **(J)** Histograms of contrast scores for DE testing of each cell type against all other cells. The dashed lines indicate significance threshold values at the FDR level *a* = 0.05.

**Fig. D29.**
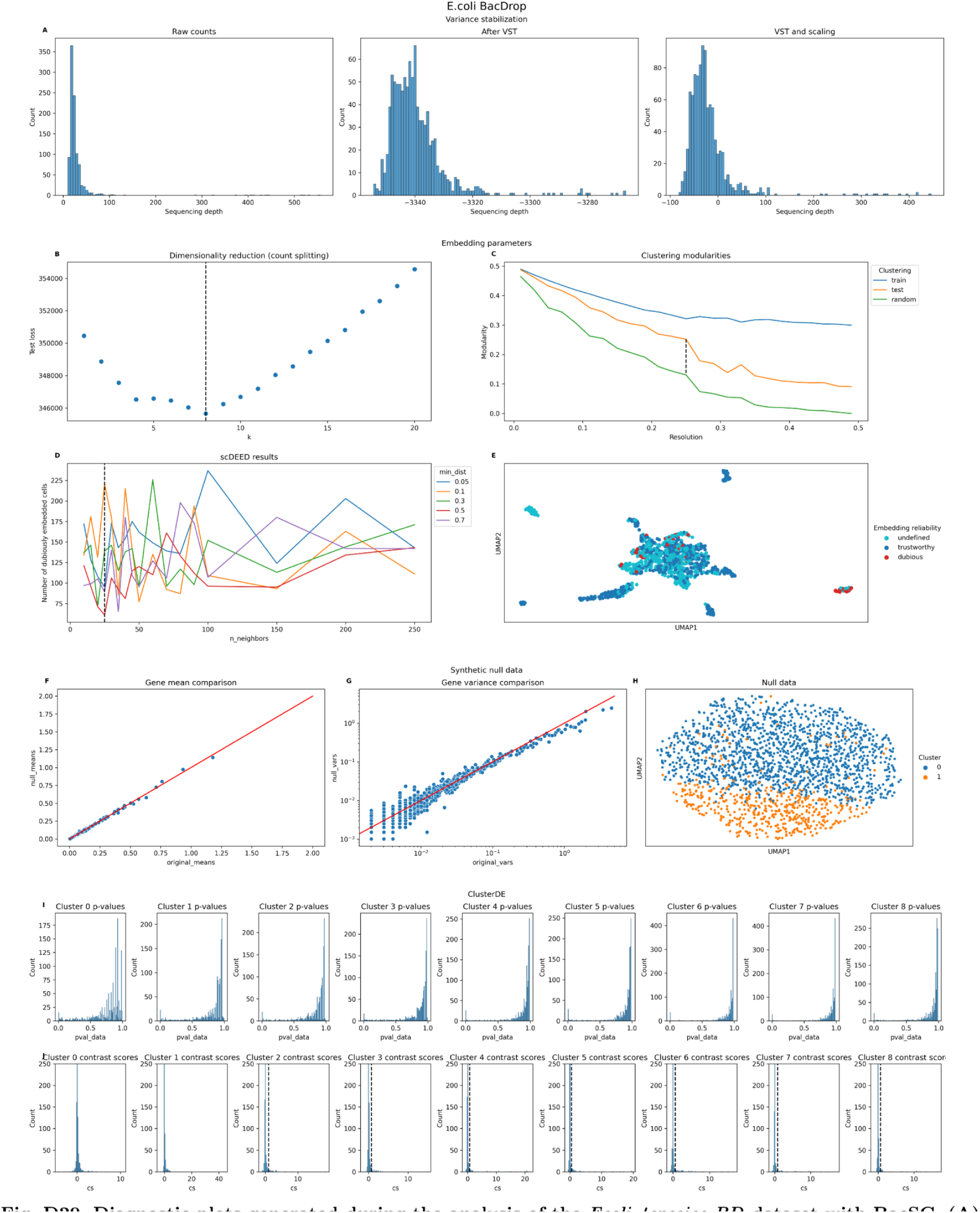
Diagnostic plots generated during the analysis of the *Ecoli_J,species_BD* dataset with BacSC. **(A)** Histograms of cell sequencing depth for raw gene expression data (left), after variance-stabilizing transform (middle), and after gene-wise scaling (right). **(B)** Selection of latent dimensionality through count splitting. The dashed line indicates *kopt* selected by BacSC. **(C)** Selection of clustering resolution. The dashed line indicates the maximal gap in modularity between the training data Leiden clustering applied to the test data and a random clustering on the test data. **(D)** Selection of *nneighbors* and *mindist* parameters for UMAP embedding with scDEED. The dashed black line shows the value of *nneighbors* selected by BacSC. **(E)** UMAP embedding generated with BacSC-selected parameters, colored by embedding reliability determined by scDEED with the same parameters. **(F)** Comparison of gene means of original and synthetic null data for DE testing. **(G)** Comparison of gene variances of original and synthetic null data for DE testing. **(H)** UMAP of synthetic null data for DE testing of each cell type against all other cells, colored by the two-group clustering determined for calculation of contrast scores. (I) Histograms of uncorrected p-values for DE testing of each cell type against all other cells. **(J)** Histograms of contrast scores for DE testing of each cell type against all other cells. The dashed lines indicate significance threshold values at the FDR level *a* = 0.05.

**Fig. D30.**
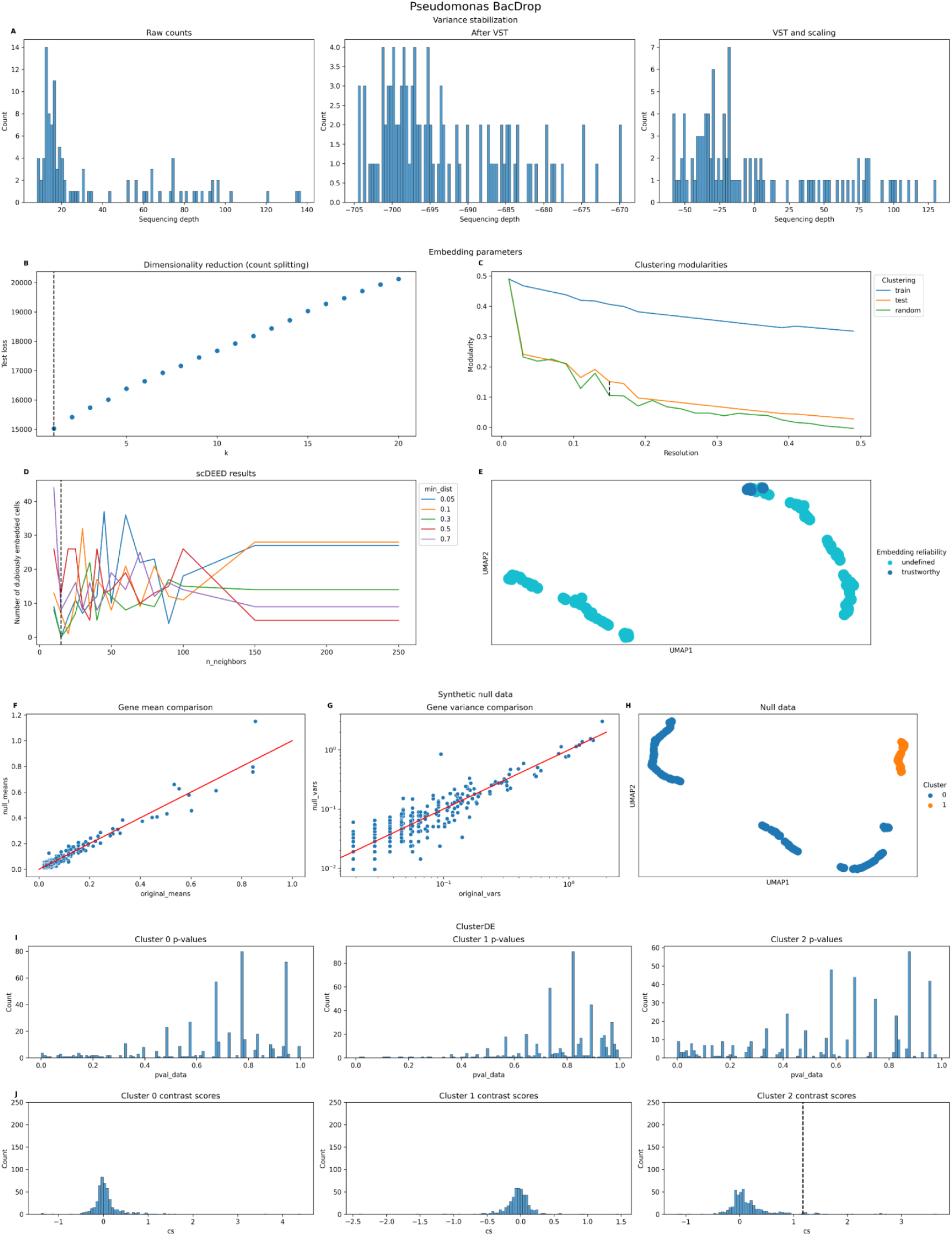
Diagnostic plots generated during the analysis of the *Pseudomonas_4species_BD* dataset with BacSC. **(A)** Histograms of cell sequencing depth for raw gene expression data (left), after variance-stabilizing transform (middle), and after gene-wise scaling (right). **(B)** Selection of latent dimensionality through count splitting. The dashed line indicates *kopt* selected by BacSC. **(C)** Selection of clustering resolution. The dashed line indicates the maximal gap in modularity between the training data Leiden clustering applied to the test data and a random clustering on the test data. **(D)** Selection of *nneighbors* and *mindist* parameters for UMAP embedding with scDEED. The dashed black line shows the value of *nneighbors* selected by BacSC. **(E)** UMAP embedding generated with BacSC-selected parameters, colored by embedding reliability determined by scDEED with the same parameters. **(F)** Comparison of gene means of original and synthetic null data for DE testing. **(G)** Comparison of gene variances of original and synthetic null data for DE testing. **(H)** UMAP of synthetic null data for DE testing of each cell type against all other cells, colored by the two-group clustering determined for calculation of contrast scores. (I) Histograms of uncorrected p-values for DE testing of each cell type against all other cells. **(J)** Histograms of contrast scores for DE testing of each cell type against all other cells. The dashed lines indicate significance threshold values at the FDR level °’ = 0.05.

**Fig. D31.**
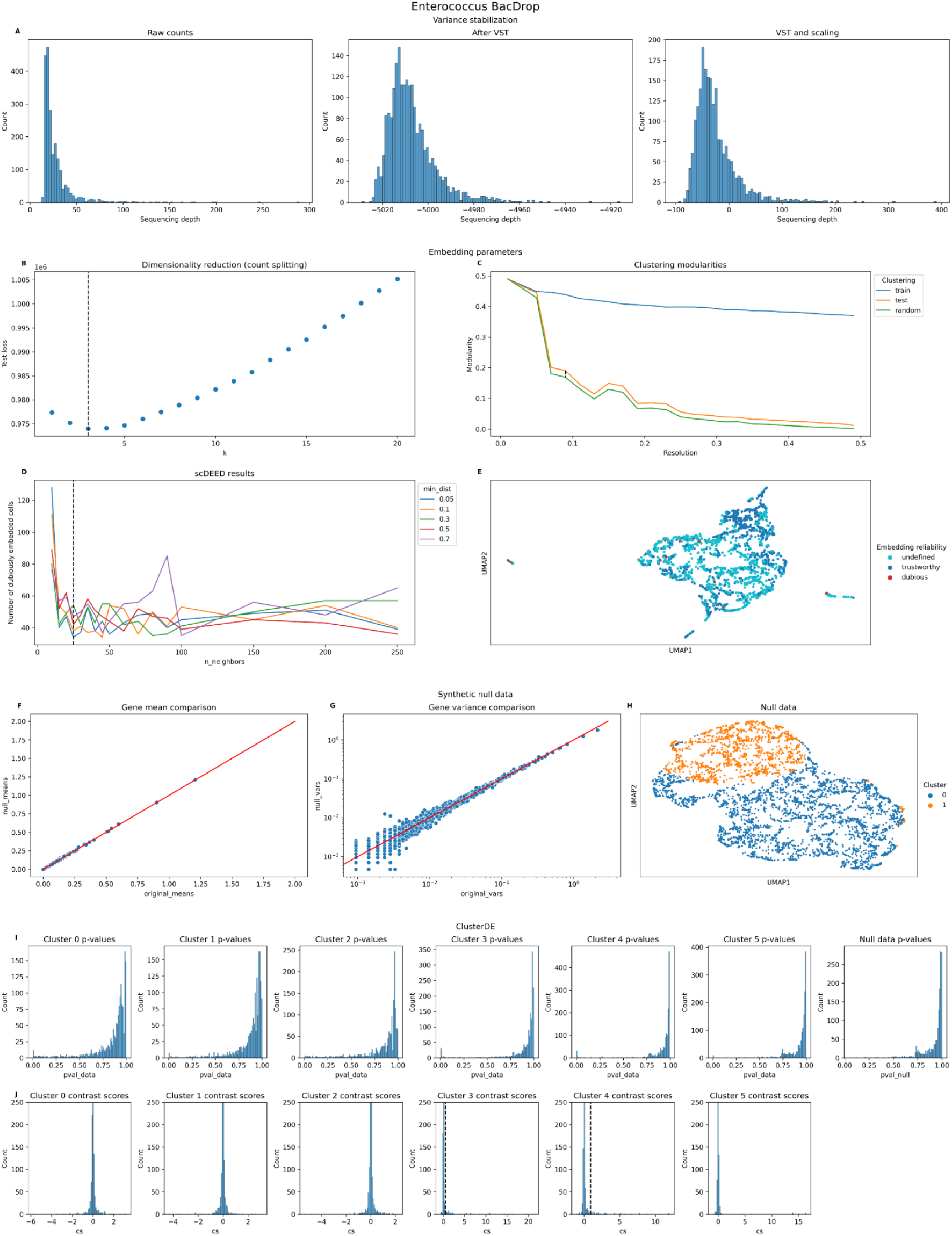
Diagnostic plots generated during the analysis of the *Efaecium_4species_BD* dataset with BacSC. **(A)** Histograms of cell sequencing depth for raw gene expression data (left), after variance-stabilizing transform (middle), and after gene-wise scaling (right). **(B)** Selection of latent dimensionality through count splitting. The dashed line indicates *kopt* selected by BacSC. **(C)** Selection of clustering resolution. The dashed line indicates the maximal gap in modularity between the training data Leiden clustering applied to the test data and a random clustering on the test data. **(D)** Selection of *nneighbors* and *mindist* parameters for UMAP embedding with scDEED. The dashed black line shows the value of *nneighbors* selected by BacSC. **(E)** UMAP embedding generated with BacSC-selected parameters, colored by embedding reliability determined by scDEED with the same parameters. **(F)** Comparison of gene means of original and synthetic null data for DE testing. **(G)** Comparison of gene variances of original and synthetic null data for DE testing. **(H)** UMAP of synthetic null data for DE testing of each cell type against all other cells, colored by the two-group clustering determined for calculation of contrast scores. (I) Histograms of uncorrected p-values for DE testing of each cell type against all other cells. **(J)** Histograms of contrast scores for DE testing of each cell type against all other cells. The dashed lines indicate significance threshold values at the FDR level *a* = 0.05.

## Appendix E Supplementary tables

**Table E1.** Dimensionality and summary statistics of datasets after quality control with BacSC. If not stated otherwise, statistics are in terms of counts/absolute values.

| Dataset | Cells | Genes | Minimum<br>seq. depth | Maximum<br>seq. depth | Median<br>seq.<br>depth | Zero counts<br>(percentage) | Maximum<br>count | 95% quantile | 99% quantile |
| --- | --- | --- | --- | --- | --- | --- | --- | --- | --- |
| Pseudomonas_balanced_PB | 1544 | 5553 | 413 | 5704 | 794.5 | 0.862 | 136.0 | 1.0 | 3.0 |
| Pseudomonas_li_PB | 1255 | 5540 | 360 | 4464 | 647.0 | 0.881 | 80.0 | 1.0 | 2.0 |
| Ecoli_balanced_PB | 3386 | 3968 | 103 | 495 | 163.0 | 0.963 | 14.0 | 0.0 | 1.0 |
| Bsub_minmed_PB | 2784 | 2952 | 141 | 1289 | 325.0 | 0.911 | 45.0 | 1.0 | 2.0 |
| Bsub_damage_PB | 13801 | 2959 | 268 | 1839 | 555.0 | 0.861 | 110.0 | 1.0 | 3.0 |
| Bsub_MPA_PB | 6703 | 2937 | 136 | 948 | 267.0 | 0.940 | 105.0 | 1.0 | 2.0 |
| Klebs_anitbiotics_BD | 19638 | 2500 | 14 | 275 | 21.0 | 0.992 | 13.0 | 0.0 | 0.0 |
| Klebs_untreated_BD | 48511 | 2500 | 12 | 728 | 21.0 | 0.991 | 30.0 | 0.0 | 0.0 |
| Klebs_BIDMC35_BD | 9168 | 2500 | 15 | 371 | 45.0 | 0.990 | 26.0 | 0.0 | 0.0 |
| Klebs_4species_BD | 315 | 1265 | 9 | 196 | 19.0 | 0.978 | 10.0 | 0.0 | 1.0 |
| Ecoli_4species_BD | 983 | 1301 | 10 | 556 | 21.0 | 0.981 | 35.0 | 0.0 | 1.0 |
| Pseudomonas_4species_BD | 103 | 628 | 8 | 137 | 18.0 | 0.953 | 7.0 | 0.0 | 1.0 |
| Efaecium_4species_BD | 2113 | 1606 | 12 | 289 | 22.0 | 0.985 | 19.0 | 0.0 | 1.0 |

**Table E2.** Overview over filtering thresholds used for quality control, number of removed barcodes, and hyperparameters determined during the course of BacSC in each dataset. Both *P.aeruginosa* datasets generated with ProBac-seq were already quality-controlled in CellRanger and therefore needed no further cell filtering for minimal sequencing depth. The ”Data distribution” column denotes the data distribution determined for count splitting (see Methods). ”NB” stands for the Negative Binomial distribution, ”Poi” denotes the Poisson distribution.

| Dataset | Minimum sequencing depth | Minimum cells per gene | Median absolute deviation cutoff (*nmads*) | Number of removed barcodes | Data distribution | Latent dimension ($k_{opt}$) | n_neighbors | min_dist | clustering resolution |
| --- | --- | --- | --- | --- | --- | --- | --- | --- | --- |
| Pseudomonas_balanced_PB | - | 2 | 5 | 108 | NB | 3 | 150 | 0.30 | 0.15 |
| Pseudomonas_li_PB | - | 2 | 5 | 71 | NB | 3 | 50 | 0.30 | 0.13 |
| Ecoli_balanced_PB | 100 | 2 | 5 | 1376 | Poi | 2 | 50 | 0.05 | 0.07 |
| Bsub_minmed_PB | 100 | 2 | 5 | 0 | Poi | 4 | 20 | 0.50 | 0.15 |
| Bsub_damage_PB | 100 | 2 | 5 | 61 | Poi | 8 | 150 | 0.30 | 0.37 |
| Bsub_sporulation_PB | 50 | 2 | 30 | 10204 | Poi | 4 | 250 | 0.30 | 0.29 |
| Bsub_MPA_PB | 100 | 2 | 10 | 197 | Poi | 2 | 10 | 0.05 | 0.03 |
| Klebs_anitbiotics_BD | 15 | 2 | 15 | 1214846 | Poi | 5 | 150 | 0.10 | 0.17 |
| Klebs_untreated_BD | 15 | 2 | 15 | 409547 | Poi | 3 | 70 | 0.05 | 0.01 |
| Klebs_BIDMC35_BD | 15 | 2 | 5 | 768 | Poi | 3 | 15 | 0.10 | 0.09 |
| Klebs_4species_BD | 15 | 2 | 10 | 8335 | Poi | 4 | 10 | 0.70 | 0.21 |
| Ecoli_4species_BD | 15 | 2 | 10 | 8671 | NB | 7 | 25 | 0.50 | 0.25 |
| Pseudomonas_4species_BD | 15 | 2 | 10 | 8089 | Poi | 1 | 15 | 0.05 | 0.15 |
| Efaecium_4species_BD | 15 | 2 | 10 | 7862 | Poi | 3 | 25 | 0.05 | 0.09 |

**Table E3.** Description of genes and rank of p-value from DE testing balanced growth versus low-iron in the combined *Pseudomonas balanced PB* and *Pseudomonas li PB* dataset. Only genes that are DE in the Copathogenex dataset for at least one of the three DE tests performed on that data are shown

| Gene | Symbol | Name | PGFam | Rank<br>(Wilcoxon<br>test) |
| --- | --- | --- | --- | --- |
| PA4514 | NaN | iron transport outer membrane recep-<br>tor | NaN | 1 |
| PA4370 | icmP | insulin-cleaving metalloproteinase<br>outer membrane protein | NaN | 2 |
| PA4515 | NaN | hydroxylase | NaN | 4 |
| PA5531 | tonB1 | transporter TonB | NaN | 6 |
| PA4709 | NaN | hemin degrading factor | NaN | 9 |
| PA4710 | phuR | heme/hemoglobin uptake outer mem-<br>brane receptor PhuR | NaN | 10 |
| PA4516 | NaN | hypothetical protein | NaN | 11 |
| PA4707 | NaN | ABC transporter permease | NaN | 13 |
| PA0472 | NaN | RNA polymerase sigma factor | RNA polymerase ECF-type sigma fac-<br>tor | 14 |
| PA0672 | hemO | heme oxygenase | Heme oxygenase HemO, associated<br>with heme uptake | 16 |
| PA2468 | foxI | ECF sigma factor FoxI | FIG006045: Sigma factor, ECF sub-<br>family | 17 |
| PA2426 | pvdS | extracytoplasmic-function sigma-70<br>factor | Sigma factor PvdS, controlling<br>pyoverdin biosynthesis | 18 |
| PA4371 | NaN | hypothetical protein | NaN | 19 |
| PA4513 | NaN | oxidoreductase | NaN | 20 |
| PA0929 | NaN | two-component response regulator | Two-component transcriptional<br>response regulator, LuxR family | 21 |
| PA2467 | foxR | anti-sigma factor FoxR | Iron siderophore sensor protein | 24 |
| PA4468 | sodM | superoxide dismutase | NaN | 26 |
| PA3530 | NaN | hypothetical protein | NaN | 28 |
| PA0931 | pirA | outer membrane receptor FepA | TonB-dependent receptor; Outer<br>membrane receptor for ferric enter-<br>obactin and colicins B, D | 31 |
| PA5217 | NaN | iron ABC transporter substrate-<br>binding protein | NaN | 34 |
| PA3899 | NaN | RNA polymerase sigma factor | NaN | 36 |
| PA4470 | fumC1 | fumarate hydratase | NaN | 39 |
| PA4708 | phuT | heme-transporter PhuT | NaN | 40 |
| PA4227 | pchR | transcriptional regulator PchR | NaN | 42 |
| PA1911 | femR | sigma factor regulator FemR | Iron siderophore sensor protein | 43 |
| PA4168 | fpvB | second ferric pyoverdine receptor FpvB | NaN | 45 |
| PA0930 | NaN | two-component sensor | two-component sensor | 55 |
| PA1912 | femI | ECF sigma factor FemI | FIG006045: Sigma factor, ECF sub-<br>family | 59 |
| PA3900 | NaN | transmembrane sensor | NaN | 71 |
| PA1300 | NaN | ECF subfamily sigma-70 factor | FIG006045: Sigma factor, ECF sub-<br>family | 73 |
| PA0471 | NaN | transmembrane sensor | Putative transmembrane sensor | 79 |
| PA4706 | NaN | hemin importer ATP-binding subunit | NaN | 81 |
| PA2033 | NaN | hypothetical protein | Siderophore-interacting protein | 86 |
| PA1365 | NaN | siderophore receptor | Ferrichrome-iron receptor @ Iron<br>siderophore receptor protein | 99 |
| PA4471 | NaN | hypothetical protein | NaN | 105 |
| PA4705 | NaN | hypothetical protein | NaN | 108 |
| PA1802 | clpX | ATP-dependent protease ATP-binding<br>subunit ClpX | ATP-dependent Clp protease ATP-binding subunit ClpX | 113 |
| PA1301 | NaN | transmembrane sensor | Iron siderophore sensor protein | 137 |
| PA4467 | NaN | hypothetical protein | NaN | 153 |
| PA0800 | NaN | hypothetical protein | FIG024006: iron uptake protein | 154 |
| PA4469 | NaN | hypothetical protein | NaN | 155 |
| PA5148 | NaN | hypothetical protein | NaN | 158 |

**Table E4.** Description of clusters for the *Bsub minmed PB* dataset. The table shows number of cells, minimal FDR (q value) over all genes, and number of differentially expressed genes at three different FDR levels.

| Cell Type | Number of cells | minimal q | DE genes, FDR = 0.05 | DE genes, FDR = 0.1 | DE genes, FDR = 0.2 |
| --- | --- | --- | --- | --- | --- |
| 0 | 1219 | 0.059649 | 0 | 1804 | 2195 |
| 1 | 897 | 0.148019 | 0 | 0 | 1517 |
| 2 | 386 | 0.257908 | 0 | 0 | 0 |
| 3 | 262 | 0.027027 | 47 | 50 | 50 |
| 4 | 20 | 0.035714 | 28 | 34 | 62 |

**Table E5.** Description of clusters for the *Klebs antibiotics BD* dataset. The table shows number of cells, minimal FDR (q value) over all genes, and number of differentially expressed genes at three different FDR levels.

| Cell Type | Number of cells | minimal q | DE genes, FDR = 0.05 | DE genes, FDR = 0.1 | DE genes, FDR = 0.2 |
| --- | --- | --- | --- | --- | --- |
| 0 | 7954 | 0.076923 | 0 | 13 | 25 |
| 1 | 5960 | 0.102564 | 0 | 0 | 111 |
| 2 | 3262 | 0.052632 | 0 | 62 | 667 |
| 3 | 1843 | 0.016129 | 96 | 122 | 673 |
| 4 | 255 | 0.111111 | 0 | 0 | 41 |
| 5 | 223 | 0.029412 | 69 | 83 | 113 |
| 6 | 74 | 0.016667 | 102 | 121 | 160 |
| 7 | 67 | 0.012987 | 102 | 118 | 133 |

**Table E6.** Description of clusters for the *Pseudomonas balanced PB* dataset. The table shows number of cells, minimal FDR (q value) over all genes, and number of differentially expressed genes at three different FDR levels.

| Cell Type | Number of cells | minimal q | DE genes, FDR = 0.05 | DE genes, FDR = 0.1 | DE genes, FDR = 0.2 |
| --- | --- | --- | --- | --- | --- |
| 0 | 571 | 0.273290 | 0 | 0 | 0 |
| 1 | 484 | 0.020833 | 51 | 71 | 81 |
| 2 | 415 | 0.028825 | 5056 | 5209 | 5209 |
| 3 | 74 | 0.045455 | 22 | 23 | 27 |

**Table E7.** Description of clusters for the *Pseudomonas li PB* dataset. The table shows number of cells, minimal FDR (q value) over all genes, and number of differentially expressed genes at three different FDR levels.

| Cell Type | Number of cells | minimal q | DE genes, FDR = 0.05 | DE genes, FDR = 0.1 | DE genes, FDR = 0.2 |
| --- | --- | --- | --- | --- | --- |
| 0 | 516 | 0.014925 | 82 | 4462 | 4850 |
| 1 | 446 | 0.798621 | 0 | 0 | 0 |
| 2 | 239 | 0.030303 | 5105 | 5210 | 5210 |
| 3 | 54 | 0.029412 | 34 | 35 | 36 |

**Table E8.** Description of clusters for the combined *Pseudomonas balanced PB* and *Pseudomonas li PB* dataset. The table shows number of cells, minimal FDR (q value) over all genes, and number of differentially expressed genes at three different FDR levels.

| Cell Type | Number of cells | minimal q | DE genes, FDR = 0.05 | DE genes, FDR = 0.1 | DE genes, FDR = 0.2 |
| --- | --- | --- | --- | --- | --- |
| 0 | 777 | 0.041667 | 24 | 31 | 44 |
| 1 | 773 | 1.000000 | 0 | 0 | 0 |
| 2 | 576 | 0.025000 | 43 | 54 | 66 |
| 3 | 396 | 0.120000 | 0 | 0 | 34 |
| 4 | 194 | 0.025000 | 50 | 66 | 71 |
| 5 | 124 | 0.029412 | 34 | 36 | 36 |

**Table E9.** Description of clusters for the *Ecoli balanced PB* dataset. The table shows number of cells, minimal FDR (q value) over all genes, and number of differentially expressed genes at three different FDR levels.

| Cell Type | Number of cells | minimal q | DE genes, FDR = 0.05 | DE genes, FDR = 0.1 | DE genes, FDR = 0.2 |
| --- | --- | --- | --- | --- | --- |
| 0 | 1132 | 0.416667 | 0 | 0 | 0 |
| 1 | 796 | 0.006452 | 1423 | 1821 | 2374 |
| 2 | 729 | 1.000000 | 0 | 0 | 0 |
| 3 | 729 | 0.055556 | 0 | 281 | 562 |

**Table E10.** Description of clusters for the *Bsub damage PB* dataset. The table shows number of cells, minimal FDR (q value) over all genes, and number of differentially expressed genes at three different FDR levels.

| Cell Type | Number of cells | minimal q | DE genes, FDR = 0.05 | DE genes, FDR = 0.1 | DE genes, FDR = 0.2 |
| --- | --- | --- | --- | --- | --- |
| 0 | 5166 | 0.263282 | 0 | 0 | 0 |
| 1 | 3734 | 0.086339 | 0 | 2694 | 2694 |
| 2 | 3246 | 0.083924 | 0 | 2049 | 2467 |
| 3 | 576 | 1.000000 | 0 | 0 | 0 |
| 4 | 526 | 0.778626 | 0 | 0 | 0 |
| 5 | 422 | 0.500000 | 0 | 0 | 0 |
| 6 | 131 | 0.100000 | 0 | 0 | 11 |

**Table E11.**
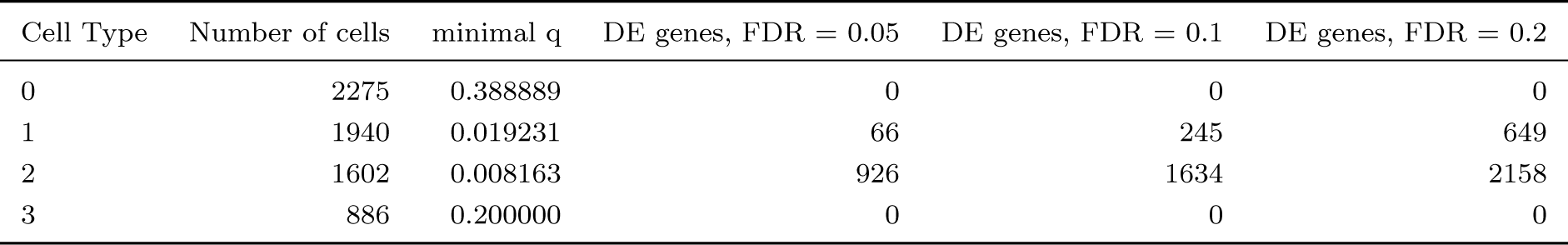
Description of clusters for the *Bsub MPA PB* dataset. The table shows number of cells, minimal FDR (q value) over all genes, and number of differentially expressed genes at three different FDR levels.

| Cell Type | Number of cells | minimal q | DE genes, FDR = 0.05 | DE genes, FDR = 0.1 | DE genes, FDR = 0.2 |
| --- | --- | --- | --- | --- | --- |
| 0 | 2275 | 0.388889 | 0 | 0 | 0 |
| 1 | 1940 | 0.019231 | 66 | 245 | 649 |
| 2 | 1602 | 0.008163 | 926 | 1634 | 2158 |
| 3 | 886 | 0.200000 | 0 | 0 | 0 |

**Table E12.** Description of clusters for the *Klebs untreated BD* dataset. The table shows number of cells, minimal FDR (q value) over all genes, and number of differentially expressed genes at three different FDR levels.

| Cell Type | Number of cells | minimal q | DE genes, FDR = 0.05 | DE genes, FDR = 0.1 | DE genes, FDR = 0.2 |
| --- | --- | --- | --- | --- | --- |
| 0 | 44236 | 0.007299 | 148 | 161 | 362 |
| 1 | 2194 | 0.005988 | 324 | 412 | 676 |
| 2 | 2081 | 0.095238 | 0 | 21 | 21 |

**Table E13.** Description of clusters for the *Klebs BIDMC35 BD* dataset. The table shows number of cells, minimal FDR (q value) over all genes, and number of differentially expressed genes at three different FDR levels.

| Cell Type | Number of cells | minimal q | DE genes, FDR = 0.05 | DE genes, FDR = 0.1 | DE genes, FDR = 0.2 |
| --- | --- | --- | --- | --- | --- |
| 0 | 2504 | 0.050000 | 0 | 20 | 26 |
| 1 | 1892 | 1.000000 | 0 | 0 | 0 |
| 2 | 1807 | 0.125000 | 0 | 0 | 8 |
| 3 | 1589 | 0.066667 | 0 | 15 | 86 |
| 4 | 914 | 0.008696 | 1047 | 1237 | 1561 |
| 5 | 255 | 0.142857 | 0 | 0 | 41 |
| 6 | 207 | 0.142857 | 0 | 0 | 7 |

**Table E14.** Description of clusters for the *Klebs 4species PB* dataset. The table shows number of cells, minimal FDR (q value) over all genes, and number of differentially expressed genes at three different FDR levels.

| Cell Type | Number of cells | minimal q | DE genes, FDR = 0.05 | DE genes, FDR = 0.1 | DE genes, FDR = 0.2 |
| --- | --- | --- | --- | --- | --- |
| 0 | 137 | 0.071429 | 0 | 14 | 53 |
| 1 | 96 | 0.019608 | 62 | 86 | 111 |
| 2 | 42 | 0.041667 | 24 | 31 | 36 |
| 3 | 26 | 0.333333 | 0 | 0 | 0 |
| 4 | 14 | 0.062500 | 0 | 29 | 30 |

**Table E15.** Description of clusters for the *Ecoli 4species PB* dataset. The table shows number of cells, minimal FDR (q value) over all genes, and number of differentially expressed genes at three different FDR levels.

| Cell Type | Number of cells | minimal q | DE genes, FDR = 0.05 | DE genes, FDR = 0.1 | DE genes, FDR = 0.2 |
| --- | --- | --- | --- | --- | --- |
| 0 | 592 | 0.258621 | 0 | 0 | 0 |
| 1 | 107 | 0.062500 | 0 | 97 | 148 |
| 2 | 83 | 0.030303 | 33 | 41 | 82 |
| 3 | 56 | 0.040000 | 43 | 72 | 84 |
| 4 | 33 | 0.025641 | 39 | 58 | 69 |
| 5 | 30 | 0.018519 | 56 | 62 | 72 |
| 6 | 29 | 0.021277 | 52 | 54 | 63 |
| 7 | 28 | 0.027027 | 37 | 37 | 53 |
| 8 | 25 | 0.043478 | 43 | 47 | 53 |

**Table E16.** Description of clusters for the *Pseudomonas 4species PB* dataset. The table shows number of cells, minimal FDR (q value) over all genes, and number of differentially expressed genes at three different FDR levels.

| Cell Type | Number of cells | minimal q | DE genes, FDR = 0.05 | DE genes, FDR = 0.1 | DE genes, FDR = 0.2 |
| --- | --- | --- | --- | --- | --- |
| 0 | 42 | 0.206897 | 0 | 0 | 0 |
| 1 | 32 | 1.000000 | 0 | 0 | 0 |
| 2 | 29 | 0.043478 | 23 | 65 | 144 |

**Table E17.** Description of clusters for the *Efaecium 4species PB* dataset. The table shows number of cells, minimal FDR (q value) over all genes, and number of differentially expressed genes at three different FDR levels.

| Cell Type | Number of cells | minimal q | DE genes, FDR = 0.05 | DE genes, FDR = 0.1 | DE genes, FDR = 0.2 |
| --- | --- | --- | --- | --- | --- |
| 0 | 943 | 0.755102 | 0 | 0 | 0 |
| 1 | 589 | 1.000000 | 0 | 0 | 0 |
| 2 | 488 | 0.571429 | 0 | 0 | 0 |
| 3 | 36 | 0.018868 | 63 | 73 | 99 |
| 4 | 33 | 0.022727 | 48 | 53 | 89 |
| 5 | 24 | 0.100000 | 0 | 0 | 11 |

